# Structures of the human transcription factor brachyury offer insights into DNA recognition, and identify small molecule binders for the development of degraders for cancer therapy

**DOI:** 10.1101/2024.06.06.597736

**Authors:** Joseph A Newman, Angeline E Gavard, Nergis Imprachim, Hazel Aitkenhead, Hadley E. Sheppard, Paul A. Clarke, Mohammad Anwar Hossain, Louisa Temme, Hans J. Oh, Carrow I. Wells, Zachary W. Davis-Gilbert, Paul Workman, Opher Gileadi, David H. Drewry

## Abstract

The transcription factor brachyury is a member of the T-Box family of transcription factors. It is active during embryogenesis and is required for the formation of the posterior mesoderm and the notochord in vertebrates. Aside from its role in embryogenesis, brachyury plays an essential role in tumour growth of the rare chordoma bone cancer and is implicated in other solid tumours. Given that brachyury is minimally expressed in healthy tissues, these findings suggest that brachyury is a potential therapeutic target in cancer. Unfortunately, as a ligandless transcription factor, brachyury has historically been considered undruggable. To investigate direct targeting of brachyury by small molecules, we initially determined the structure of human brachyury both in complex with its cognate DNA and in the absence of DNA. Analysis of these structures provided insights into brachyury DNA binding and the structural context of the G177D variant which is strongly associated with chordoma risk. We used these structures to perform a crystallographic fragment screen of brachyury and identify hotspot regions on numerous pockets on the brachyury surface. Finally, we have performed follow-up chemistry on fragment hits and describe the structure-based progression of a thiazole-containing chemical series. Excitingly, we have produced brachyury binders with low µM potency that can serve as starting point for further medicinal chemistry efforts. These data show that brachyury is ligandable and provides an example of how crystallographic fragment screening may be used to find ligands to target protein classes that are traditionally difficult to address using other approaches.

## Introduction

The T-box family of transcription factors comprises 18 members in humans and is found in a wide range of other animal species. T-box genes can function as both transcriptional activators and repressors and are required for the development of multiple cell types. Defects in these genes can result in a variety of developmental disorders^1^. They encode a highly conserved sequence-specific DNA binding domain of around 200 residues and a transcriptional regulatory domain in the C-terminus. The DNA binding domain forms an immunoglobulin-like fold and contacts to DNA via an unusual insertion of an α-helix into the minor groove. All T-box proteins recognize a common consensus sequence named the T-box binding element (AGGTGTGAAA) with family member-specific variations found in the preferences for repeats, orientations (inverted or tandem) and spacing of such repeats^2^. It is still not clearly understood how these preferences relate to *in vivo* target gene promoter specificity as binding sites identified by Chip-Seq analysis generally only show single T-box elements^3,4^ and further specificity may be achieved by interactions with other factors.

Brachyury (encoded by the *TBXT* gene, also formerly known as *T*) is the founding member of the family and was first identified in 1927^5^. It has subsequently been studied in detail due to its roles in development of the notochord and posterior mesoderm^6^. Brachyury is minimally expressed after day 13 of human development except for a select few tissues (thyroid, testes, and pituitary gland), however aberrant expression is found in various cancers. The most well-established association is with the bone cancer chordoma where brachyury is used as a diagnostic marker. Chordoma is a rare cancer (affecting around 1 in 1,000,000 people per year) that occurs along the spinal-cord and is thought to originate from remnants of the embryonic notochord. Chordoma currently lacks effective targeted therapies, with primary treatment by surgical resection followed by adjuvant radiotherapy and cytotoxic chemotherapy. Reoccurrence is common and further surgical intervention may become debilitating, with the median survival of chordoma patients being 7.7 years^7^. The *TBXT* (*brachyury*) gene is frequently duplicated in chordoma^8^, is required for growth in chordoma disease models^3,9^ and has been identified by genome-scale CRISPR-Cas9 screening as the top selectively essential gene in chordoma ^10^. Further associations of brachyury in chordoma are found with the observation of a common single nucleotide polymorphism rs2305089 being strongly associated with the risk of developing chordoma in European populations^11^ (odds ratio of 6.1). This variant encodes a glycine to aspartate substitution in the brachyury DNA binding domain. It is not clear at present what precise role this substitution plays in the protein function and role in chordoma, although previous *in vitro* assays suggest an impact on DNA binding ^12^.

Further to its role as a genetic driver of chordoma, there is evidence that brachyury plays a role in various epithelial cancers where it promotes growth, induces an Epithelial to Mesenchymal transition (EMT) and subsequently metastasis from the primary tumour site ^13-15^. This is thought to be due to the presence of a T-box DNA site in the E-cadherin promoter^14^ which is a key player in cell adhesion. Brachyury expression in these cancers has also been correlated with resistance to chemo- and radiotherapy^16^.

Overall, these data are consistent with brachyury being the genetic driver of the malignant phenotype that is expressed almost uniquely in the tumour cells, making it a biologically ideal therapeutic target in chordoma. However, such ligandless transcription factors have traditionally been thought to be difficult to inhibit with small molecules due to their lack of druggable pockets and polar nature of the DNA binding interface. In this study, we have addressed whether brachyury can be targeted with small molecules with sufficient affinity and specificity. We initially determined the crystal structures of human brachyury, and its chordoma-associated variant, bound to two different T-box binding element containing DNA molecules. Examination and comparison of the structures provided insights into DNA recognition and allowed direct comparison of WT and variant structures bound to DNA. We have also developed crystal systems of both WT and variant DNA binding domains in the absence of DNA that diffract to high resolution and used these crystals in a high-throughput crystallographic fragment screen to identify ligandable pockets in brachyury. We found 29 fragments bound in 6 clusters which we used as starting points for development of more potent binders. Here, we describe preliminary structure-guided optimization of compounds from a thiazole-containing series to a low µM level of potency which have the potential for further medicinal chemistry optimization. These compounds could lead to chemical probes of brachyury functions or be further developed into warheads for protein degradation.

### Results

### Crystal structure of WT and G177D brachyury in complex with DNA

Like many transcription factors, nearly half of the brachyury protein is intrinsically disordered (Figure S 1) and therefore not amenable to crystallography to capture reliable structures. We therefore aimed to crystallize the DNA binding domain of human brachyury. Using the previously determined structure of brachyury from *Xenopus laevis* as a guide^17^, we designed oligonucleotides containing a palindromic arrangement of T-box binding elements of varying length (22-30 nucleotides) for crystallization of human brachyury. We were able to crystallise the WT human brachyury DNA complex to 2.25 Å resolution using a construct spanning residues 41-224 with a 24 base pair DNA sequence. The chordoma-associated G177D variant brachyury was crystallized with a 26 base pair DNA sequence and diffracted to 2.15 Å resolution. Both structures feature generally high-quality electron density maps and have been restrained to standard bond lengths and angles. A summary of the data collection and refinement statistics can be found in Table 1.

**Table 1.** Data collection and refinement statistics for brachyury DNA complexes.

|  |  |  |  |
| --- | --- | --- | --- |
|  | **TBXT + DNA** | **TBXT(D177) + DNA** | **TBXT + ssDNA** |
| Space group | P 21 | P 43 2 2 | P 61 2 2 |
| Cell dimensions, *a, b, c* (Å) | 75.0 37.1 110.8 | 75.3 75.3 288.5 | 63.0 63.0 218.5 |
| Wavelength (Å) | 0.92 | 0.92 | 0.97 |
| Resolution (Å) | 29.6 – 2.25 (2.33 – 2.25) | 72.0 – 2.15 (2.21 – 2.15) | 109 – 2.55 (2.62 – 2.55) |
| R<sub>merge</sub> | 0.123 (0.689) | 0.113 (2.560) | 0.109 (2.810) |
| I/σI | 6.1 (1.5) | 12.9 (1.2) | 15.9 (1.0) |
| CC1/2 | 0.992 (0.653) | 0.997 (0.777) | 1.00 (0.593) |
| Completeness % | 97.6 (94.1) | 99.9 (98.7) | 100 (100) |
| Multiplicity | 3.4 (3.4) | 25.5 (23.1) | 17.4 (15.7) |
| No. Unique reflections | 27866 (2466) | 46387 (3687) | 9158 (666) |
| **Refinement statistics** |  |  |  |
| Resolution | 29.6 – 2.25 | 72.0 – 2.15 | 109 – 2.55 |
| R<sub>work</sub>/R<sub>free</sub> (%) | 22.56/28.82 | 22.14/25.20 | 27.33/30.18 |
| No. atoms | 4131 | 4105 | 1937 |
| Protein | 2843 | 2924 | 1449 |
| Solvent | 335 | 112 | 2 |
| DNA | 952 | 1060 | 486 |
| Average B factors (Å<sup>2</sup>) |  |  |  |
| All atoms | 38 | 69 | 82 |
| Protein | 37 | 70 | 82 |
| Solvent | 42 | 61 | 62 |
| DNA | 30 | 65 | 82 |
| Wilson B | 34 | 59 | 71 |
| R.M.S. deviations |  |  |  |
| Bond lengths (Å) | 0.007 | 0.008 | 0.006 |
| Bond angles (°) | 0.90 | 0.96 | 1.055 |
| Ramachandran plot |  |  |  |
| Favoured (%) | 99 | 99 | 92 |
| Allowed (%) | 1 | 1 | 7 |
| PDB ID | 6F58 | 6F59 | 8CDN |

**Table II.**
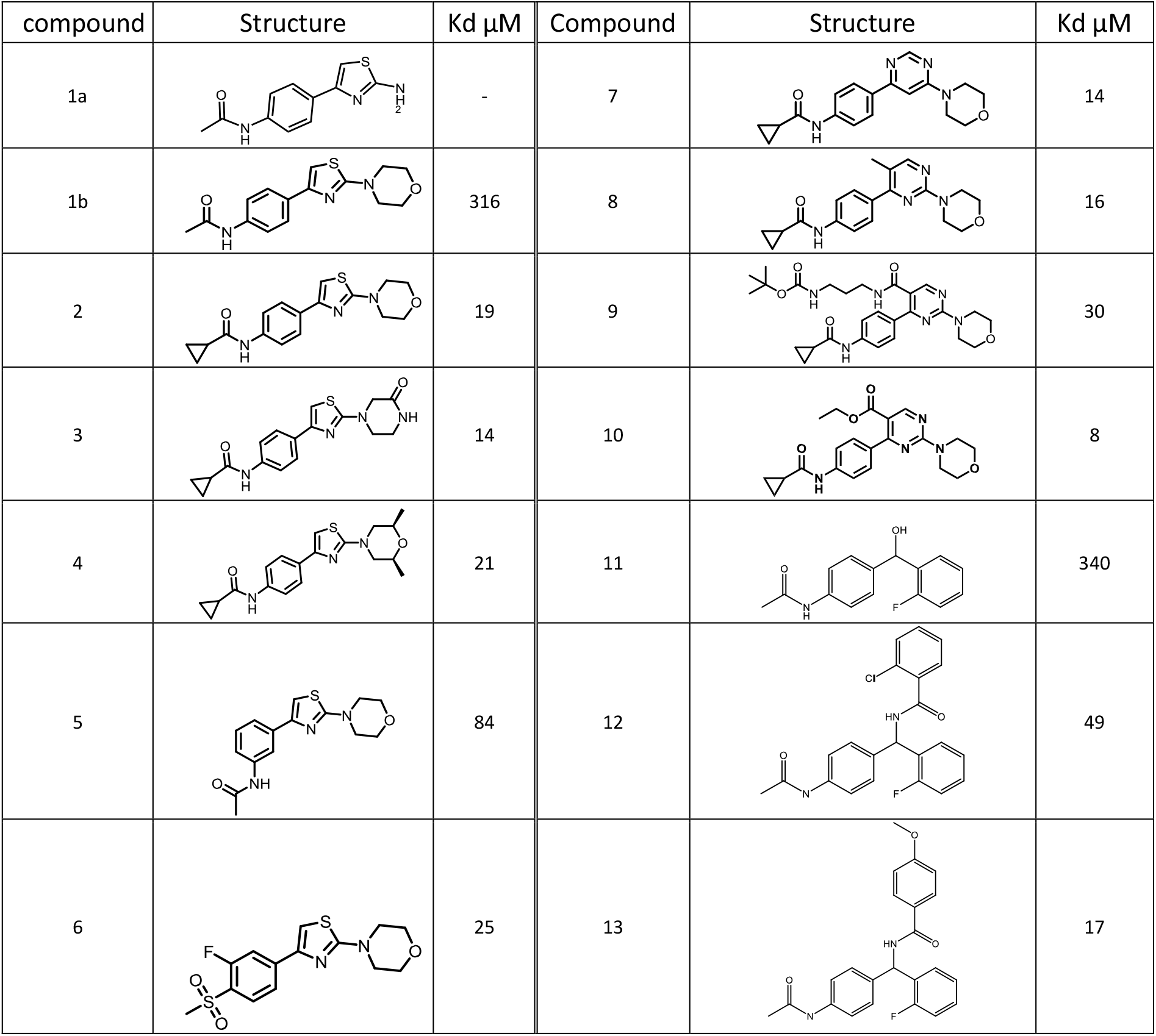
Chemical structures and binding affinity of compounds measured by SPR.

| compound | Structure | Kd μM | Compound | Structure | Kd μM |
| --- | --- | --- | --- | --- | --- |
| 1a |  | - | 7 |  | 14 |
| 1b |  | 316 | 8 |  | 16 |
| 2 |  | 19 | 9 |  | 30 |
| 3 |  | 14 | 10 |  | 8 |
| 4 |  | 21 | 11 |  | 340 |
| 5 |  | 84 | 12 |  | 49 |
| 6 |  | 25 | 13 |  | 17 |

Overall, the structures are very similar to previous T-box family DNA complexes, including the highly related Xenopus brachyury structure (92 % sequence identities and 0.9 Å RMSD). The structures feature a modified immunoglobulin-like β-sandwich fold with additional helical elements between the first and second strands and at the C-terminus (Figure 1A). As for other T-box structures which have been obtained with near identical DNA sequences, two copies of brachyury bind to the DNA in a 2-fold symmetrical arrangement with a small interface between subunits located towards the N-terminal end of the first β-sheet. Contacts to the DNA are made via loops between strands A and B, c and c’, and two α-helices following on from strand G at the C-terminal end of the DNA binding domain (Figure 1B). Unusually, compared to other transcription factor families which recognize sequences via major groove interactions, the final helix is inserted into the minor groove of the DNA with two conserved aromatic side chains F213 and F217 inserting deep into the groove and making contacts with the bases. As has been noted previously^17^, only two direct hydrogen bonds are made between protein and nucleobases, R69 contacting the N7 of the guanine at position 5 of the motif, and the main chain carbonyl of F213 contacting the guanine at position 7 of the motif (Figure 1C). It has been speculated that these contacts are not sufficient to explain the observed pattern of recognition observed in *in vitro* site selection experiments^18^. The contact area between the two protein subunits is small (in the region of 200 Å^2^) and the contribution of this interface towards the possible cooperative binding on similarly spaced palindromic sites has been questioned following structures of related T-box binding proteins on the same DNA sequence which do not feature this interface^19,20^. Furthermore, the fact that palindromic arrangements of T-Box binding elements have not been identified in any known T-box target gene promoters has led to questions of what if any role cooperativity may play *in vivo*^3,4^.

**Figure 1.**
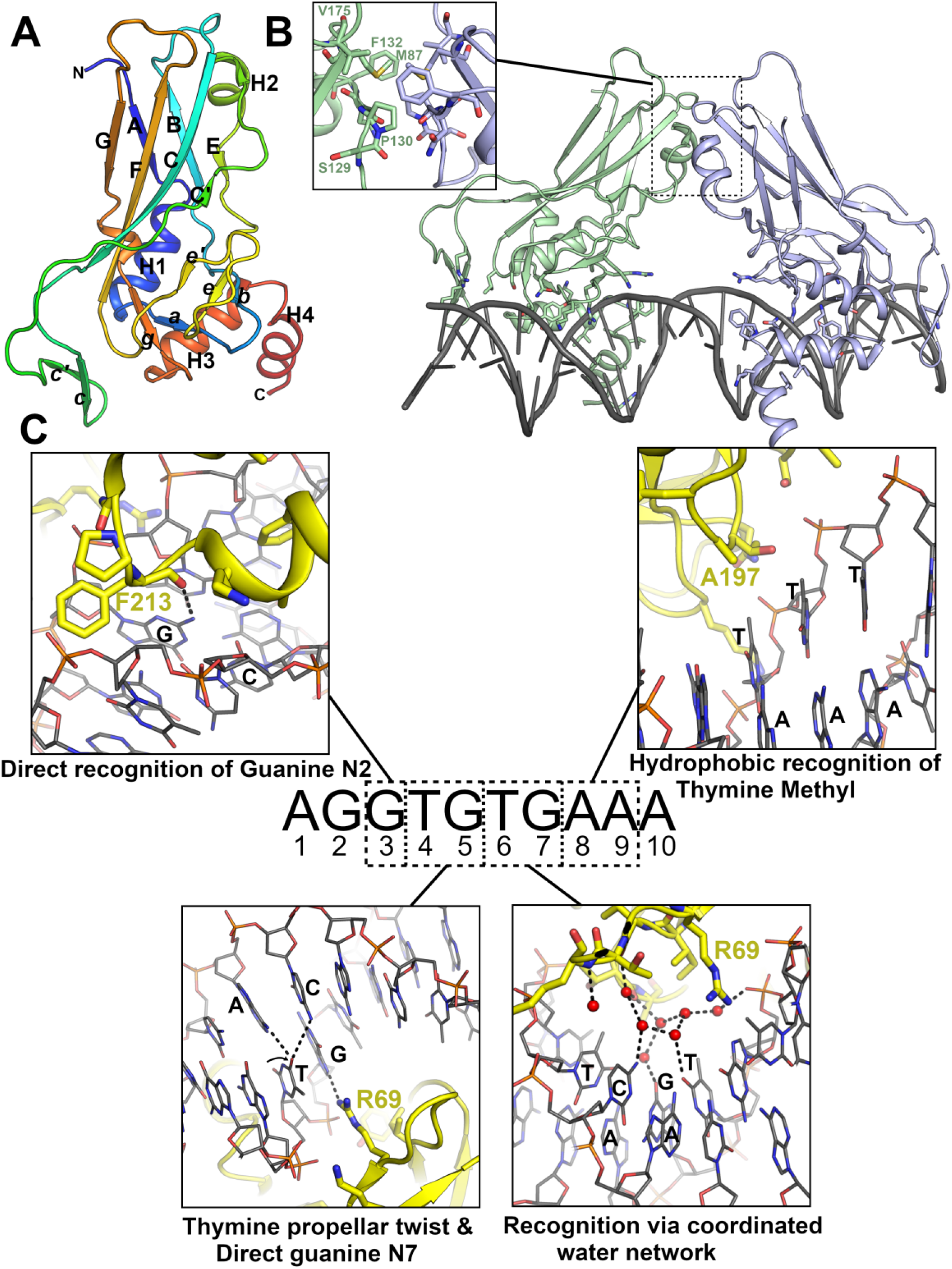
Structure of brachyury in complex with DNA. (**A**) Overall structure of human brachyury DNA binding domain with secondary structure elements labelled in accordance with the immunoglobulin fold nomenclature. (**B**) Structure of human brachyury bound to a palindromic DNA containing an inverted repeat of the T-box recognition element. Two brachyury protomers bind to each DNA half site with an unusual insertion of a helix into the minor groove and form a small interface between the two chains which is shown in the inset. (**C**) Details of the recognition of specific DNA base pairs by brachyury. For each nucleotide position in the T-box recognition element the mode of recognition is indicated in the inset box with key residues and nucleotides labelled.

### Structural basis of T-box binding element recognition

The relatively high resolution of our DNA complex structures has allowed us to examine the DNA protein interface in detail including identification of water-mediated interactions. We have also obtained a crystal structure of WT brachyury (at 2.7Å resolution) in complex with a single T-box binding element half-site oligo which allows us to directly compare binding interfaces and DNA distortions between single and palindromic sites. In addition to the direct contacts that define specificities at bases 3 and 5 (detailed above), recognition of a Thymine at base 4 has been previously attributed to the possibility of formation of a bifurcated hydrogen bond with the Cytosine from base 5 which stabilizes the significant buckle of this base pair^19^ (Figure 1C). Hydrophobic interactions with Thymine methyl groups at bases 8 and 9 to A197 and T196 likely contribute to specificities for those sites (Figure 1C). Recognition of bases at positions 1 and 2 has been attributed largely to indirect mechanisms (intrinsic deformability of the DNA) as these bases are distant from the protein, which is consistent with the more relaxed specificity at these sites. The means of recognition of bases at the sixth and seventh positions has so far been more elusive, as no direct contacts are made to the protein, yet these bases are conserved in most T-box binding sites. We note that a cluster of water molecules are found in this region in conserved positions when comparing our structure with the TBX3 DNA complex obtained at 1.7 Å resolution^19^. The water network contacts the O4 of Thymine at position 6, the Guanine O6, and Cytosine N4 at position 7. We accept that the configuration of this water network may vary depending on the DNA sequence but using known rules of hydrogen bonding donors and acceptors we can deduce that some degree of recognition is possible, for example the presence of a H-bond donor at Cytosine N7 of base 7 would specify through the water network a H-bond acceptor at O4 of base 6 (Figure 1C).

As has been found in previous T-box family DNA structures, the DNA contains some significant distortions from regular B-form geometry, most notably a widening of the minor groove to accommodate the insertion of α-4 (Figure S2). Both halves of the palindromic site display similar distortions, and both ends show a significant narrowing of the minor groove (from 11.7 Å in canonical B-form DNA to around 9 Å at both ends). This narrowing may be related to the presence of an A-tract (defined as a stretch of 4 or more A-T base pairs without a TpA step) on either end of the palindromic sequence, which is known to facilitate DNA bending and narrowing of the minor groove^21^. Given that only the last two T-A bases of the A-tract are within the T-box motif it is not clear if this narrowing is a feature in T-box family DNA recognition. Comparing the structures of brachyury bound to a palindromic DNA with a 12-base pair single site reveals that most of the contacts at the interface are maintained (Figure S3) although the end narrowing of the minor groove is not present in the single site DNA which also does not contain a full A-tract sequence (Figure S3), suggesting perhaps that this may just be a feature of the palindrome used for crystallization.

### Comparison of WT and G177D structures

As would be expected with only a single substitution, the structures of the WT and G177D variant are very similar (RMSD 0.8 Å) with only minor differences seen in the ordering and conformations of various loops that are presumably flexible (Figure 2A). The G177D variant itself lies in a loop between strands F and G and is situated away from the DNA interface on the opposite side of the protein. The conformation of this loop is significantly altered (Figure 2A) with a new salt-bridge formed between the substituted D177 residue and R174 possibly explaining the altered conformation in the variant structure. Furthermore, the G177D substitution is within a GGP peptide sequence that lies in a restricted region of Ramachandran space. The PHI and PSI angles adopted by the Glycine are not permitted for a non-glycine residue, particularly in the case of a pre-proline which is more restrictive due to steric hindrance around the proline Cδ. Whilst the differences in this loop may be significant on a local level, we do not see a possible way for these changes to be transmitted through the protein to the DNA binding site. The G177D substitution is however close to the site of the small protein interface that is created between subunits when binding on DNA containing T-box binding sites in a palindromic arrangement. Thus, it is plausible that the substitution may affect the cooperativity of DNA binding as has been suggested previously^12^. Analysis of the interfaces between subunits using the program PISA^22^ reveals some very minor differences in the interface areas (210 Å^2^ for the G177D versus 219 Å^2^ for the WT) and in the calculated energetic contributions towards toward complex formation (-4.8 Kcal/mol versus -4.5 Kcal/Mol).

**Figure 2.**
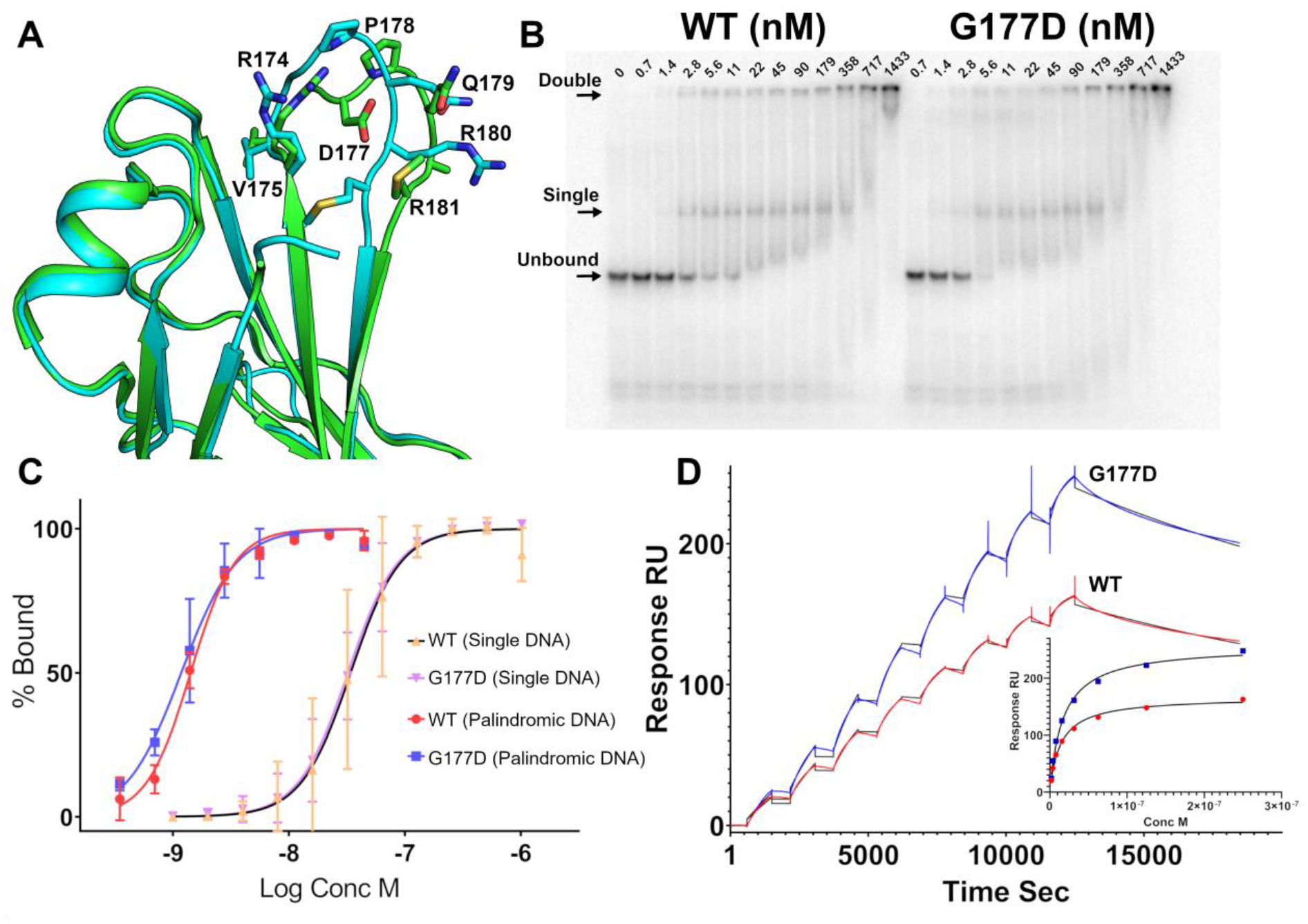
Analysis of the G177D variant on brachyury structure and DNA binding. (**A**) Comparison of the structures of WT (Cyan) and G177D (Green) DNA binding domains in the vicinity of the loop 175-181. (**B**) Representative electrophoretic mobility shift assay (EMSA) gel of WT and G177D full length brachyury binding to a 50 bp DNA with palindromic repeat of the T-box recognition element. (**C**) Quantification of EMSA data comparing WT and G177D brachyury binding to a single site and palindromic repeat of the T-box recognition element. (**D**) Analysis and comparison of WT and G177D brachyury binding to palindromic DNA by surface plasmon resonance. The main plot shows responses generated for sequential injections of DNA with full length WT or G177D brachyury immobilized on the sensor surface. The black line shows the fit to the data using a bivalent analyte model and inset shows the same data fitted using a concentration-response curve.

### The G177D variant does not substantially change DNA binding affinity or cooperativity

Although the structural differences may be small, it is plausible that they contribute towards a difference in the cooperative binding at sites with inverted repeats of the T-box binding element. We tested this using *in vitro* DNA binding assays and compared the full-length WT and the G177D variant by electrophoretic mobility shift assay (EMSA) on a range of DNA probes containing palindromic, single sites and natural promoter sequences. We found, as others have observed, a preference for palindromic repeats with an apparent K_d_ of around 1 nM (Figure 2B, 2C). Binding of a single site probe or a probe from a natural brachyury target promoter (sequence identified from Fibroblast growth factor FGF8) gave a lower apparent K_d_ of 30-40 nM (Figure 2C & S4). Two shifted species could be observed on the gel for the palindromic probe which we presume represent both singly and doubly bound species. From examination of the bands on the EMSA assay there does appear to be some degree of cooperativity as both upper and lower bands appear at approximately the same point in the titration, rather than the upper band lagging behind the lower for a noncooperative independent binding scenario. However, contrary to previous reports, we do not see any significant differences in either the binding affinity or the degree of cooperativity when comparing the WT and variant proteins.

We have also tested the binding of a palindromic DNA sequence to brachyury WT and G177D via Surface Plasmon Resonance (SPR) with the protein immobilized on to a streptavidin sensor surface. A clear high affinity binding interaction can be observed which can be fit as a concentration response with apparent dissociation constants of 14.8 ± 2.4 nM for the WT and 17.9 ± 2.9 nM for the G177D variant. Fitting the data using a kinetic model is only possible with a bivalent binding model consistent with two binding sites on the DNA (Figure 2D). The reason for the cooperative binding has been the subject of some debate within the T-box family, as the small interface formed between brachyury subunits is not present in other T-box family member DNA structures^19,20^ although these family members do show some preference for palindromic DNA in *in vitro* assays^23^. We suggest a possible explanation for this phenomenon, that the widening of the minor groove for binding at one site lowers the energy barrier (due to proximity) for the widening of the minor groove for the nearby binding of a second protomer. The fact that the two T-box sites are arranged in the inverted orientation and in the closest possible proximity for binding both sites without steric clashes would fit with this ‘through DNA’ model. The small interface between subunits may play an additional role in a subset of T-box family members such as brachyury.

### The Chordoma associated variant is expressed equally to WT

While the G177D variant does not substantially change DNA binding *in vitro*, we aimed to further characterize its biological relevance. Previous studies investigated the role of the G177D variant using engineered isogenic cellular systems that expressed only WT or G177D brachyury^3^. Each of these isogenic cell lines was viable, suggesting that the G177D brachyury variant is not a sole chordoma driver. Furthermore, there was no significant difference in the downstream brachyury target genes as identified by ChIP-sequencing. We aimed to understand the consequence of the G177D variant within its endogenous context as other reasons for its association with chordoma could lie in protein stability or expression levels. However, examining the stability of WT and G177D brachyury by DSF shows only a small TM shift of ∼0.7 degrees, with the WT variant appearing to be very slightly more thermostable than the chordoma-associated variant (Figure 3A) and thus likely not an explanation for observed differences in activity. To characterise the prevalence of the G177D variant in chordoma cells, we obtained a panel of 9 chordoma lines and genotyped them for the presence of the G177D variant. These cells included lines derived from both sacral and clival chordomas, primary and metastatic tumour sites, and the paediatric chordoma cell line UM-Chor5. Sanger sequencing of exon 4 of the *TBXT* locus confirmed the presence of the WT (G) or G177D (A) variant (Figure 3B, Figure 3C). We find that the majority of chordoma cell lines (6/9) genotyped are homozygous for the chordoma-associated SNV. The majority of homozygous variants across the chordoma lines mirrors previous patient findings^11^. Interestingly, the chordoma cell line U-CH17M, derived from a metastatic chordoma tumour^24^, does not encode the G177D variant and is the first chordoma cell line without the variant identified to date. SW480/SW620 colorectal cancer cells, control non-chordoma cell lines which have been shown to express brachyury^9^, also do not encode the G177D brachyury variant. Given a minority of chordoma cell lines are also heterozygous for the brachyury variant, we aimed to determine the expression levels of the WT vs G177D alleles. Using publicly available RNA-sequencing data^10^, we confirm that each variant in the heterozygous cell line UM-Chor1 is equally expressed. Together, these data, coupled with previous findings^3^, suggest a brachyury-directed chordoma therapy must target both WT and G177D brachyury.

**Figure 3.**
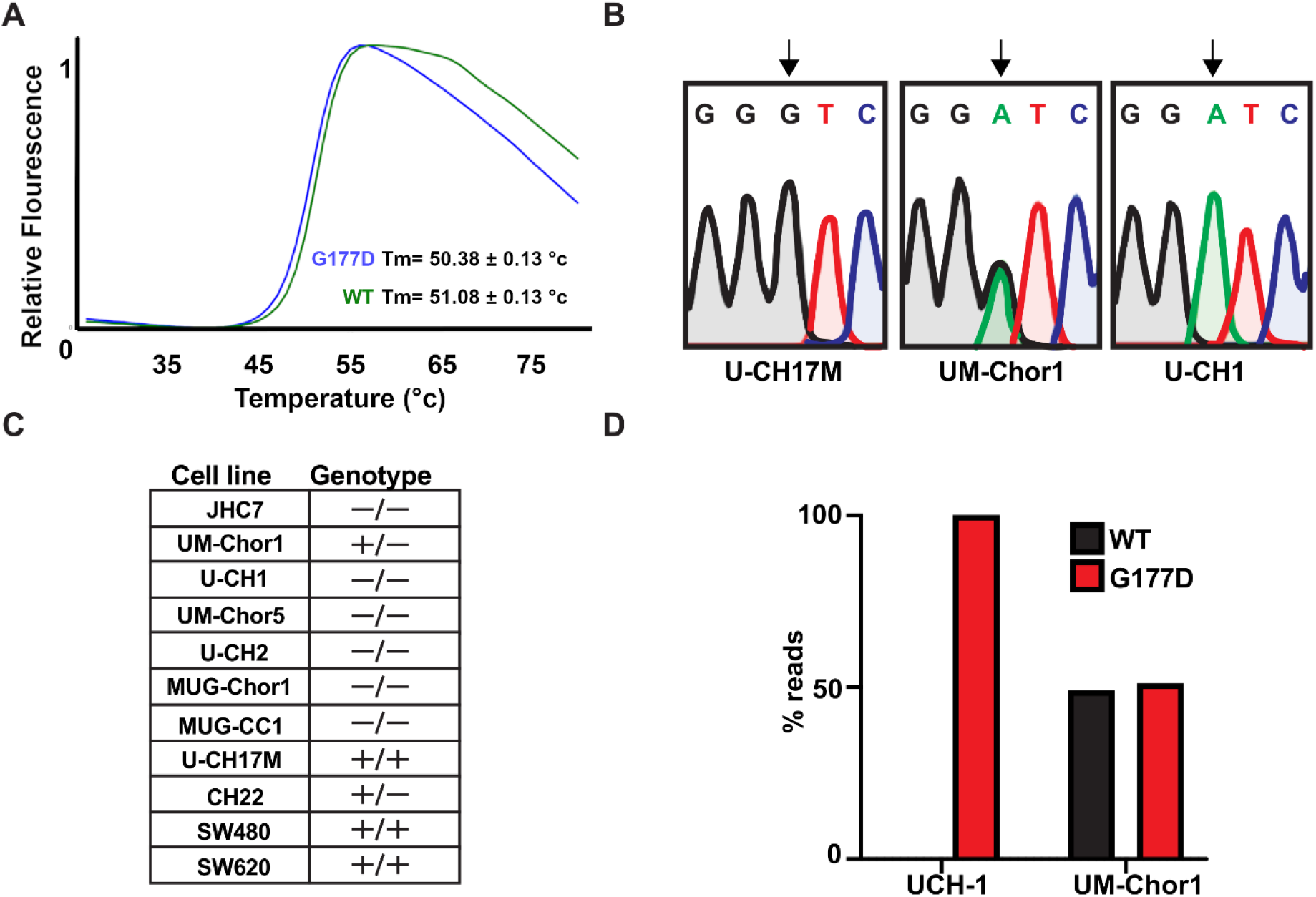
Analysis of the common G177D variant thermal stability and prevalence in human chordoma cell lines. (**A**) Differential Scanning Fluorimetry (DSF) measurement of the thermal stability of WT and G177D full length brachyury proteins. (**B**) DNA sequencing chromatograms of three chordoma cell lines in the vicinity of the polymorphism. WT brachyury is encoded by a Guanine nucleotide at the 3^rd^ position whilst G177D variant has an Adenine. (**C**) Table of the genotypes of various chordoma cell lines (+ signifies WT and – the G177D variant). (**D**) RNA-Seq analysis of two chordoma cell lines showing approximately equal transcription of WT and G177D alleles in a heterozygous human chordoma cell line UM-Chor1.

### Crystallographic fragment screening of brachyury

The causal role of brachyury in human chordoma cancers coupled with the absence of detectable expression of brachyury in most adult human tissues, indicate that, from the point of view of potential effectiveness and therapeutic index, it is an ideal drug target biologically for drug discovery. However, ligandless transcription factors have traditionally been considered very difficult to target due to the presence of intrinsic disorder and lack of well-defined pockets for small molecule binding^25^. We have tested this premise and performed a druggability analysis of our brachyury structures using the ICM pocket finder algorithm (Figure 4A)^26,27^. As expected, only three pockets of moderate volume could be found that were not predicted to be druggable based on the Drug-Like-Density (DLID) metric^28^. Similar results were obtained using the ligandability analysis in the canSAR knowledgebase^29^, with the largest pocket displaying properties on the lower limit for a classically a druggable site (kinase) and similar to a challenging yet druggable target (Bcl 2) (Figure S5). Such calculations can serve as useful barometers, but new inhibition modalities such as PROteolysis Targeting Chimeras (PROTACS) offer promise even for these difficult targets^30,31^.

**Figure 4.**
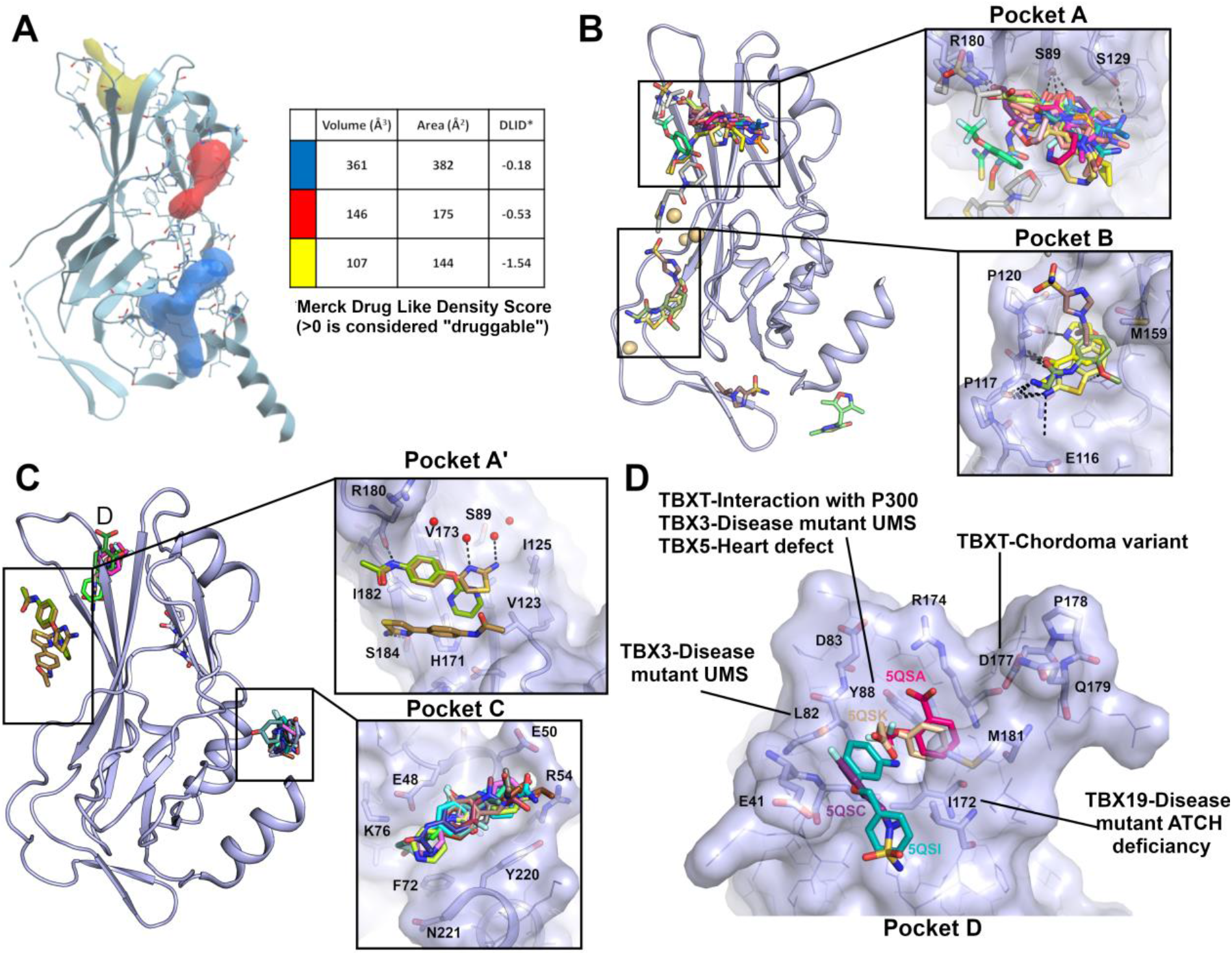
Druggability analysis and fragment screening of brachyury. (**A**) Pocket analysis of brachyury DNA binding domain. Pockets are depicted as solid blobs of red, blue, and yellow with volumes, areas, and predicted druggability scores shown in the inset table. Druggability scores are calculated using ICM-Pro according to the method of Sheridan *et. al*.^*28*^. (**B**) Fragment screening results for WT brachyury DNA binding domain with hotspot pockets shown in the insets in a surface representation with key residues labelled. (**C**) Fragment screening results for G177D brachyury DNA binding domain with hotspot pockets shown in the insets in a surface representation with key residues labelled. (**D**) Close up view of the fragments bound to pocket D with residues involved in disease mutations in other T-box family members labelled.

To further investigate the potential for direct inhibition or degradation of brachyury, and to discover starting points for structure-based drug discovery, we have performed a crystallographic fragment screen on both WT and G177D variant brachyury DNA binding domains. Crystallographic fragment screening relies on direct detection of fragments in electron density maps, the reliability of which is greatly enhanced with crystals that diffract to high resolution. For X-ray fragment screening in the absence of DNA, we used a shorter construct (amino acids 41-211) lacking the C-terminal helix that inserts into the DNA minor groove, which was observed to be flexible in our early DNA-free crystallization efforts. WT and G177D crystals were obtained in different conditions with different crystal forms although both generally diffracted to around 1.6 Å. A total of 609 fragments were soaked into WT crystals and diffraction data were collected and analysed using the PanDDA algorithm^32^, yielding a total of 30 fragment hits across 27 datasets. For the G177D crystals a total of 616 fragments were soaked yielding 17 fragment hits across 16 datasets (Supplementary Table I and II). Comparisons between the two datasets show (perhaps surprisingly) very little overlap between the fragment hits, despite the same fragment library being soaked. Presumably this is due to a combination of the different accessibility of sites in the two crystals forms (Figure S6) and the sensitivity of X-ray fragment binding to minor changes in conformation or chemical conditions.

In the WT crystals a significant hot spot (pocket A) was identified near the N-terminal end of strand *c*, with 24 fragments bound, the majority of which make a hydrogen bond to S89 with additional contacts to R180 and S129 (Figure 4B). A further 4 fragments are bound to a pocket (pocket B) formed between strands *c’* and *e’* which make polar contacts to the main chain of loop 116-120 (which was found to be partially disordered in the DNA complex structures), and side chains of M159 and E116 (Figure 4B). The WT fragment screen was performed in crystallization conditions containing cadmium chloride with typically 5 cadmium ions are bound to surface sites in the protein making contacts to exposed cysteine and histidine residues. We acknowledge that the presence of these ions may have some influence on fragment binding although only a single fragment was observed to bind primarily to a cadmium ion. In the G177D form, a hotspot pocket (pocket C) was located near the C-terminal end of the final helix which contains 9 fragments that make polar contacts to R54, E48 and K76 (Figure 4C). Although this pocket lies on a crystallographic 2-fold symmetry axis, the fragments are close to the DNA interface (∼8 Å away) and the pocket is significantly larger and extends down to the DNA interface in the DNA complex structures with the longer construct. Three fragments were observed to bind in a roughly equivalent site to pocket A in the WT (pocket A’) although these fragments engage in more hydrophobic type interactions with I182, L91 and V123 (Figure 4C). Finally, 4 fragments bound to a pocket near the N-terminus (pocket D) which appears to be induced partially by ligand binding. Two fragments with a benzene ring occupy a relatively buried cavity that was also observed to bind an 2-Methyl-2,4-pentanediol (MPD) molecule in the G177D DNA complex structure (a component of the crystallization solutions), and make contacts to R174, Y88 and M181. This pocket is also lined by G/D 177 and binding may be specific to the G177D variant conformation (Figure 4D). Across both screens only two fragments (PDB entries 5QRM and 5QRW) are bound to the regions containing the DNA binding interface, in line with the general view that such interfaces do not generally contain tractable pockets for small molecules.

It is not known whether any of the pockets identified are sites that upon compound binding will lead to inhibition of brachyury or modulation of its function, although pockets B and C approach to within around 8 Å of the DNA interface with the potential for fragment growth in that direction. Pockets A and A’ are close (within 3 Å) to the potential dimer interface that is formed when binding to palindrome T-box sites and may be extended to disrupt DNA cooperativity without directly having to compete with DNA binding. Finally, pocket D is distant from both dimerization and DNA binding sites but has been implicated to have a likely role in downstream signalling due to the observation of a dependence for a tyrosine residue at position 88 for interaction with P300, a component of the histone modification machinery^33^. Residues lining this pocket are well conserved in the T-box family and have been shown to be important *in vivo* as clusters of point mutations from several different human genetic diseases map to this site (Figure 4D), and mutational analysis in the murine T-bet protein indicates critical roles for this pocket in interactions with permissive chromatin re-modellers including KDM6A and KDM6B^34^.

### Structure-guided optimization of fragments to potent brachyury binders

Encouraged by the discovery of ligandable pockets on brachyury, we have initiated a medicinal chemistry campaign to optimize the potency of fragment-derived molecules. We have used biophysical binding by surface plasmon resonance (SPR) as our primary assay to measure the potency of compounds irrespective of the potential to inhibit brachyury activity. Only compounds that displayed high quality sensograms with concentration-dependent responses were included for analysis (Table S2). This biophysical approach also enables the discovery of potent binders that, even if they are not inhibitors of brachyury function, could be used as warheads to induce the degradation of brachyury through a PROTAC modality. This degradation approach has been validated for brachyury in chordoma cells using the inducible degradation dTAG system^3^.

One of the promising chemical series is based on a thiazole fragment found in pocket A’ (PDB: 5QS9) which bound with two molecules near a solvent-exposed hydrophobic surface patch that is formed by the second β-sheet containing strands c,*c’*,*f* and *g* (Figure 4c). One molecule of thiazole 1a shows binding to R180 and two exposed solvent molecules, and the other molecule of thiazole 1a shows binding to S184 (Figure 5a). Initial work aimed to induce a selectivity bias between the two different binding modes. Chemistry methods and compound characterization is provided in the supplemental information. Replacement of the 2-NH_2_ of the thiazole (Table 2, Entry 1a) with a 2-morpholino moiety led to a compound (Table 2, Entry 1b) with measurable binding affinity on SPR that could be soaked into our crystals, revealing a single mode of binding equivalent to the R180-interacting pose of the original thiazole hit (Figure 5c). This fragment retains the hydrogen bond to R180 with most of the rest of the interaction mediated by hydrophobic interactions with nearby side chains of residues L91, V123, I125, V173 and I182 which form an unusual cluster of surface-exposed hydrophobic residues that are conserved amongst other T-box family structures (Figure S6). Further improvement was obtained by conversion of the acetamide to a cyclopropylacetamide which provided the most notable improvement in this limited series of analogues (Table 2, Entry 2 & Table S2) increasing the apparent binding affinity to the low µM range (14-20 µM) as measured by SPR (Figure S7). The cyclopropyl group has the potential to make favourable hydrophobic interactions with the side chains of residues I182 and M181. We hypothesize that steric clashes with crystallographic neighbours thwarted attempts to generate structures with ligands containing these cyclopropyl groups. Additional modifications at this position were also tolerated, including replacement with an ortho-fluoro methylsulfone (Table 2, Entry 6). The morpholine group could be replaced with piperazin-2-one or 3,5-*cis-*dimethylmorpholine (Table 2, compounds 3 and 4). We were able to employ either 2,4-substituted or 4,6-substituted pyrimidines (Table 2, Entry 7-10) as isosteric replacements of the thiazole ring, and compounds with these modifications retained binding activity (Table S2). Moving from the 5-membered thiazole to 6-membered pyrimidines provides an additional vector for further optimization of the series and an additional position for exploration of exit vectors for bivalent degraders (for example compound 9). In our original pyrimidine fragment (PDB 5QSD, Figure 5Bb) the pyrimidine moiety inserts into a conserved hydrophobic cleft close to L91, V123 and a polar triad of residues (H171, R169 and D93) that form a network of hydrogen bonds (Figure 5B & S6). In our initial fragment screen, we also found a similar N-phenylacetamide in the A’ pocket which contained a diphenylmethanol group (Table 2, Entry 11, Figure 5D). Further exploration of the secondary alcohol with chlorobenzamide (Table 2, Entry 12) or methoxybenzamide (Table 2, Entry 13) groups provided significant increases in potency for this limited series of compounds as measured by SPR (Table S2).

**Figure 5.**
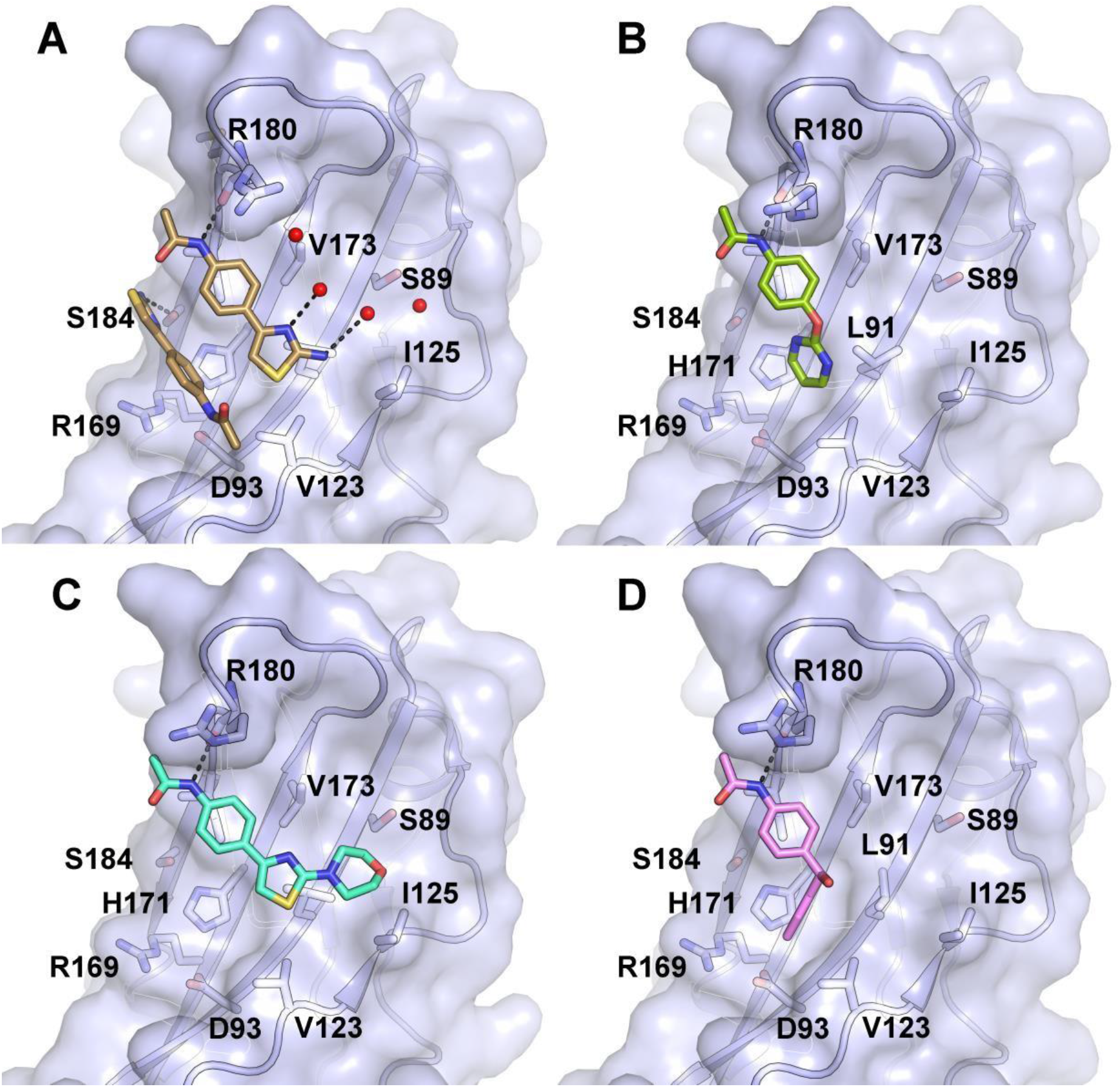
Structures of fragment derived hits binding to pocket A’. (**A**) Crystal Structure of 5QS9 in pocket A’ of brachyury. (**B**) Crystal Structure of 5QSD in pocket A’ of brachyury. These two compounds occupy very similar binding modes in this space. The primary interaction in these fragments is a hydrogen bond to ARG180 from the amide functional group (this can be observed in both 5QS9 and 5QSD). (**C**) Crystal Structure of CSC027898502 in pocket A’ of brachyury. Showing contact to ARG180, and further interactions with the hydrophobic pocket surface. (**D**) Crystal structure of compound 11 in complex with brachyruy forming hydrophobic contacts with a conserved hydrophobic cleft.

## Discussion

Directly targeting ligandless transcription factors as a cancer therapy has huge potential^35^, however, with a few notable exceptions^36-38^ this potential has yet to be realized. Direct targeting of transcription factors with small molecules remains challenging due to the combination of frequent intrinsic disorder, lack of defined druggable pockets and the dynamic nature of transcription factor complexes. In this study, we have investigated the potential for direct targeting of the human brachyury transcription factor, which has a causal role in chordoma cancers, using a structure-guided approach. We have determined structures of human brachyury both alone and in complex with DNA and have used our high-resolution APO form crystals to perform crystallographic fragment screening. This screen identified several ligandable pockets on the surface of brachyury, and we have been able through structure-guided chemical optimization to progress fragment hits to binders with apparent dissociation constants in the low µM range. Very little overlap was observed in the fragment hits between the WT and G177D fragment screens, we attribute this to the fact that the two screens were performed on different crystal forms and the influence of crystal packing and crystallization conditions, rather than the structural differences between the two forms, the exception being some of the hits in pocket D which do contact variant residues.

It is important to note that these compounds have been validated as binders and are not yet inhibitors of brachyury function. As we have shown from our RNA-Seq analysis an effective therapeutic targeting brachyury must be able to target both WT and G177D variant, and we expect this to be the case for our optimized compounds. Structural data generated to date on this series show that they bind a considerable distance from the variant site and DNA interfaces. We have tested compounds from this series using an *in vitro* fluorescence polarization-based DNA binding assay and do not find any evidence for inhibition of DNA binding, although we note that it is possible there are functional consequences for binding at this site in the cell. These compounds may serve as starting points for further development into PROTAC warheads to induce the degradation of brachyury.

The compounds we have identified bind to relatively shallow surface pockets, one of which has an unusual lipophilic character due to several surface exposed hydrophobic residues. Importantly these pockets were not identified as easily druggable by a computational analysis and hits binding to this pocket would be unlikely to be detected by conventional high-throughput screening if using a DNA binding readout. Our results demonstrate the power of the structure-guided fragment-based approach for identifying binders for classes of proteins that are otherwise generally considered intractable for drug discovery. Whilst low µM binders are not sufficiently potent for pharmacological inhibition in a classical occupancy-driven model, such moderate affinity binders have previously been shown with other protein targets to be effective as PROTAC warheads.^39^ As PROTAC warheads they participate in what has been called event-driven pharmacology, and are not required to occupy the binding site for extended periods of time and indeed may exert their effects through transient interactions that allow multiple rounds of degradation^40,41^. The work we have described here adds to the notion that emerging technologies, including fragment screening by x-ray crystallography, will soon move transcription factors out of the undruggable category, and a new reality will emerge in which transcription factors are valuable and tractable drug discovery targets^42^.

## Materials & Methods

### Cloning Expression and Purification

Constructs for WT and G177D brachyury DNA binding domain (residues 41-224) (used for DNA complex crystal structures) in pET28 were obtained as a gift from Michael Miley, University of North Carolina. Constructs for the WT and G177D truncated DNA binding domain (residues 41-211) (used for fragment screening crystals) were cloned into pSUMO-LIC vector using ligation-independent cloning to produce proteins with an N-terminal His-SUMO tag. Full length WT and G177D brachyury (used in the DSF and SPR assays) were cloned into the pHGT-Bio vector by ligation independent cloning for expression of brachyury with a N-terminal 9His-GST tag and C-terminal AVI tag for biotinylation of the target protein by co-expression with BirA enzyme in the presence of biotin^43^. All plasmids generated in this study have been deposited in Addgene.

From glycerol stocks, bacteria were inoculated in 15 ml of 1 x TB in a 50 ml tube with Kanamycin 0.05 mg/ml and 0.034 mg/ml of chloramphenicol and grown overnight in a shaker at 37°C, 250rpm. The following day, 4 ml of the overnight culture were inoculated in 1L of TB (Merck). The bacteria grew in an incubator at 37°C, with shaking at 180 rpm. Once the OD reached 2-3, IPTG (300uM) was added to the media and left overnight at 18°C. The cultures were harvested by centrifugation. For purification of WT and G177D DNA binding domain (used in 6F58 and 6F59) cell pellets were resuspended in 250 ml of Lysis Buffer (50 mM HEPES pH 7.5, 500 mM NaCl, 10 mM Imidazole, 5% Glycerol and 1 mM TCEP). The cells, on ice, were sonicated for 20 minutes with 5 seconds pulse ON and 10 seconds pulse OFF with 35% of amplitude and centrifuged for 25 minutes at 66700 x g. The supernatant was incubated for an hour at 4°C, with Nickel beads pre-washed with Lysis buffer. After one hour of batch-binding, the tubes containing the lysate were centrifuged at 700 x g at 4°C for 5 minutes and the supernatant discarded. This step was repeated twice with, respectively, 100 ml and 50 ml of Lysis Buffer. Beads were loaded on a gravity column with 20 ml of Wash Buffer (50 mM HEPES pH 7.5, 500 mM NaCl, 30 mM Imidazole, 5% Glycerol and 1 mM TCEP) and, followed by two elution of 10 ml each with Elution Buffer (50 mM HEPES pH 7.5, 500 mM NaCl, 300 mM Imidazole, 5% Glycerol and 1 mM TCEP). After an SDS-PAGE gel, the elution containing the protein was concentrated with an Amicon 10kDa concentrator and loaded on a Hi Load 16/600 Superdex 75 pg column at 1 ml/min, collecting 2-ml fractions. The fractions containing the protein were pooled together and concentrated with an Amicon 10kDa concentrator until 10 mg/ml was reached. Protein aliquots were flash frozen in Liquid Nitrogen and stored at -80°C.

For purification of WT and G177D for DNA-free Crystal forms, initial cell lysis and IMAC purification was as above. Following IMAC fractions containing TBXT were pooled, and SUMO protease was added to a final mass ratio 1:150. Cleavage was performed overnight during dialysis into dialysis buffer (50 mM HEPES pH 7.5, 500 mM NaCl, 5 % Glycerol, 1 mM TCEP) using 3500 MWCO snakeskin dialysis tubing. After dialysis the protein was concentrated with an Amicon 10kDa concentrator and loaded on a Hi Load 16/600 Superdex 75 pg column. The flowrate of the gel filtration was 1 ml/min and the volume of the fractions collected was 2 ml. The fractions containing the protein were pulled together and concentrated with an Amicon 10kDa concentrator, until the concentration 12 mg/ml was reached. Protein aliquots were stored at -80°C after being flash frozen in Liquid Nitrogen.

For purification of WT and G177D full length brachyury (used for SPR analysis) initial cell lysis and IMAC purification was as above. Following IMAC fractions containing TBXT were pooled, and TEV protease was added to a final mass ratio 1:40. Cleavage was performed overnight during dialysis into dialysis buffer (50 mM HEPES pH 7.5, 500 mM NaCl, 5 % Glycerol, 1 mM TCEP) using 3500 MWCO snakeskin dialysis tubing. After dialysis the protein was concentrated with an Amicon 30 kDa concentrator and loaded on a Hi Load 16/600 Superdex 200 pg column. All proteins were confirmed by ESI-TOF intact mass spectrometry.

### Crystallization and structure determination

For crystallization of the WT brachyury DNA complex (6F58) a self-complementary DNA oligonucleotide 5’-AATTTCACACCTAGGTGTGAAATT was dissolved to 1mM, heated to 95°C on a heat block and allowed to cool slowly over 2hrs. The protein and DNA were mixed in a 1:1.1 molar ratio (assuming a duplex DNA molecule) and sitting drop vapour diffusion crystallisation trials were set up with a Mosquito (SPT Labtech) crystallisation robot at a final concentration of 6.6 mg/ml. TBXT crystallised at 4°C in conditions containing 40% PEG300, 0.1M citrate pH 4.2. Crystals were loop-mounted and cryo-cooled by plunging directly into liquid nitrogen. Data were collected to 2.2Å resolution at Diamond light source beamline I04-1 and the structure was solved by molecular replacement using the program PHASER^44^ and the structure of Xenopus laevis brachyury^17^ (1XBR) as a search model. Refinement was performed using PHENIX REFINE^45^ to a final Rfactor = 24.2%, Rfree = 28.8%.

For crystallization of the G177D brachyury DNA complex (6F59), a self-complementary DNA oligonucleotide 5’-GAATTTCACACCTAGGTGTGAAATTC was dissolved to 1mM, heated to 95°C on a heat block and allowed to cool slowly over 2hrs. The protein and DNA were mixed in a 1:1.1 molar ratio (assuming a duplex DNA molecule) and sitting drop vapor diffusion crystallization trials were set up with a Mosquito (SPT Labtech) crystallization robot at a final concentration of 8 mg/ml. TBXT crystallised at 4°C in conditions containing 56% MPD, 0.1 M SPG pH 6.0. Crystals were loop-mounted and cryo-cooled by plunging directly into liquid nitrogen. Data were collected to 2.1Å resolution at Diamond light source beamline I04-1 and processed using DIALS^46^. The structure was solved by molecular replacement using the programme PHASER^44^ and 1XBR^17^ as a search model. Refinement was performed using PHENIX REFINE^45^ to a final Rfactor = 22.1%, Rfree = 25.2%. Data collection and refinement parameters for all DNA complex datasets are shown in Table 1.

### X-ray fragment screening

For crystallization of WT brachyury for fragment screens the protein was adjusted to 7.5 mg/ml and sitting drop vapor diffusion crystallisation trials were set up with a Mosquito (SPT Labtech) crystallisation. TBXT crystallised at 4°C in conditions containing 32% PEG400, 0.1M acetate pH 4.5, 0.1 M cadmium chloride. Crystals were loop mounted and cryo-cooled by plunging directly into liquid nitrogen.

For crystallization of G177D brachyury for fragment screens the protein was adjusted to 16 mg/ml and sitting drop vapor diffusion crystallisation trials were set up with a Mosquito (SPT Labtech) crystallisation robot. TBXT crystallised at 4°C in conditions containing 30% PEG1000, 0.1M SPG pH 7.0. Initial seed crystals were obtained from these conditions ad 4-5 crystals were crushed with a glass probe and transferred to a 50 ul solution of well solution containing a PFTE seed bead and vortexed for 3 x 20 seconds. The concentrated seed solution was diluted in well solution 1:1000 and used to seed crystals set up at 10 mg/ml in 300 nL drops with 20nL of seeds (added last).

For the WT crystals a total of 608 fragments from the DSI poised library (500 mM stock concentration dissolved in DMSO) were transferred directly to brachyury crystallization drops using an ECHO liquid handler (10 % or 50 mM nominal final concentration) and soaked for 1–3 h before being loop mounted and flash cooled in liquid nitrogen. A total of 603 crystals were mounted leading to 575 datasets the vast majority of which diffracting to 2.5 Å or higher (96%). For the G177D fragment screen 637 fragments from the DSI poised library were soaked as above. A total of 590 crystals were mounted leading to 486 datasets with the majority diffracting to 2.5 Å or higher (76%). For both crystal forms data were collected at Diamond light source beamline I04-1 and processed using the automated XChemExplorer pipeline. Structures were solved by difference Fourier synthesis using the XChemExplorer pipeline^47^. Fragment hits were identified using the PanDDA^32^ program. Refinement was performed using REFMAC^48^. A view of the electron density maps of each fragment hit is shown in Supplementary Table S1 and a summary of data collection and refinement statistics for all fragment-bound and ligand-bound datasets is shown in Supplementary Data 1.

### Electrophoretic mobility shift assays (EMSA)

To evaluate binding to a palindromic DNA sequence the following oligonucleotide pair was used: TA50-F CATGCATGCAGGGAATTTCACACCTAGGTGTGAAATTCCCATTCGTGCGA, TA50-R TCGCACGAATGGGAATTTCACACCTAGGTGTGAAATTCCCTGCATGCATG. To reduce the formation of hairpin structures, the oligos were annealed at a high concentration (>200 µM each) in 10 mM tris-HCl, pH 7.5, 50 mM NaCl by heating to 95°C in a dry block and leaving to cool to room temperature. The dsDNA was subsequently labelled with T4 polynucleotide kinase (NEB) and γ-32P-ATP. The labelled DNA was separated from the remaining ATP/ADP using a BioRad MicroBiospin P-6 column equilibrated in annealing buffer. EMSA buffer was: 25 mM HEPES, pH 7.4, 10% glycerol, 75 mM NaCl, 0.1% tween 20, 1 mM TCEP. Protein was diluted serially in this buffer and mixed with 1-5 nM of DNA diluted in the same buffer. After 10-minute incubation on ice, the samples (5 µl) were loaded on a pre-run 8% polyacrylamide gel (40:1 acrylamide/bis) in chilled TAE buffer (40 mM TRIS base, 20 mM acetic acid, 1 mM EDTA). The gel tanks were placed in an ice bucket and run for 75 minutes at 150V. The dried gels were exposed overnight using a BioRad phosphorimager screen. Results were quantified using BioRad ImageLab software are plotted using Graphpad Prism as means ± standard deviation from at least 3 independent experiments. Dissociation constants were calculated by fitting the data to a 4-parameter logistic regression equation.

### Surface Plasmon Resonance (SPR)

DNA binding analysis by SPR was performed on a Biacore s200 machine using a Series S SA sensor surface. Full length biotinylated G177D or WT brachyury was immobilized to approximately 1500 RU (immobilization at approximately 20 nM for 100 seconds) in running buffer 10 mM Hepes pH 7.5, 150 mM NaCl, 1 mM DTT, 1% DMSO. Palindromic DNA containing T-box binding sites (TA50-F + TA50-R) was titrated as an 8 x 2-fold dilution series from a high concentration of 250 nM in single cycle mode (90 seconds association with final dissociation of 600 seconds) with the highest concentration last. The data were fit with a bivalent kinetic model (in agreement with each duplex containing two T-box sites) and approximated by dose response analysis using the Biacore S200 evaluation software.

Small molecule binding by SPR was measured using a biacore 8K+ system. Full length human brachyury (G177D) containing a C-terminal AVI-tag and biotinylation site was immobilized at approximately 8000RU to a Biacore SA sensor surface (10 ug/ml for 360 seconds in buffer 10mM HEPES, pH7.5, 150mM NaCl, 1mM TCEP, 0.05%P20). Serial dilutions of compounds (6 x 2-fold dilutions starting from 200 uM or 100 uM) were injected over the surface in multi cycle mode with 60 second association time, 120 second dissociation at a flow rate of 30 ul/min in a buffer containing 10 mM HEPES pH 7.5, 150 mM NaCl, 1 mM TCEP, 0.05 % Tween 20. Solvent correction and reference compound injections were performed every 50 and 100 cycles respectively. After solvent correction compound binding dissociation constants were evaluated using a dose response (equilibrium) analysis.

## Data availability

Crystallographic coordinates and structure factors for all structures have been deposited in the Protein Data Bank with the following accession codes: 6F58, 6F59, 8CDN, 5QS6, 5QS7, 5QS8, 5QS9, 5QSA, 5QSB, 5QSC, 5QSD, 5QSE, 5QSF, 5QSG, 5QSH, 5QSI, 5QSJ, 5QSK, 5QSL, 5QRF, 5QRG, 5QRH, 5QRI, 5QRJ, 5QRK, 5QRL, 5QRM, 5QRN, 5QRO, 5QRP, 5QRQ, 5QRR, 5QRS, 5QT0, 5QRT, 5QRU, 5QRV, 5QRW, 5QRX, 5QRY, 5QRZ, 5QS0, 5QS1, 5QS2, 5QS3, 5QS4, 5QS5, 7ZL2, 8A1O, 8A7N, 7ZKF. SPR sensograms and data fits for all compounds tested in this study have been made publicly available on Zenodo (https://doi.org/10.5281/zenodo.6394811).

## Acknowledgements

This work was funded in part by grants from the Chordoma Foundation and the Mark Foundation for Cancer Research, and generous gifts to the Chordoma Research Fund by supporters of chordoma patients and chordoma research. The Structural Genomics Consortium (SGC) is a registered charity (number 1097737) that receives funds from Bayer AG, Boehringer Ingelheim, the Canada Foundation for Innovation, Eshelman Institute for Innovation, Genentech, Genome Canada through Ontario Genomics Institute, EU/EFPIA/OICR/McGill/KTH/Diamond, Innovative Medicines Initiative 2 Joint Undertaking, Janssen, Merck KGaA (also known as EMD in Canada and USA), Pfizer, the São Paulo Research Foundation-FAPESP, and Takeda. The crystallographic fragment screen was supported by the XChem facility at Diamond Light Source (proposal ID LB18145). Crystallographic data were collected at Diamond Light Source beamlines I04-1, I04 & I03 (proposals mx19301 and mx28172). We thank the CMD Biotechnology group for help with plasmid cloning, test expression, and mass spectrometry. PW acknowledges recent and current support for his research from Cancer Research UK (CRUK Programme Grants C309/A31322 and C309/A11566; Strategic Award C35696/A23187; Infrastructure Award C309/A27413; and funding for the CRUK Childrens Brain Tumour Centre of Excellence C9685/A26398/RG93685), Wellcome (Biomedical Resource and Technology Development Grant 212969/Z/18/Z to support the Chemical Probes Portal), Chordoma Foundation, Mark Foundation, Bone Tumour Research Trust, CRIS Cancer, and The Institute of Cancer Research. PW is a CRUK Life Fellow. Hadley Sheppard was funded by an AACR–CRUK Transatlantic Fellowship.

## Author contributions

D.H.D., O.G., P.W., P.A.C, C.I.W. and J.A.N conceptualization. J.A.N., A.E.G., H.A. N.I., H.E.S., M.A.H., L.T. & Z.W.D-G. Methodology. J.A.N., A.E.G., H.A. N.I., H.E.S., M.A.H., L.T. & Z.W.D-G Investigation and Data analysis. J.A.N. & M.A.H. writing-original draft. D.H.D., O.G., P.W., P.A.C, C.I.W. J.A.N., A.E.G., H.A. N.I., H.E.S., M.A.H., L.T. & Z.W.D-G., writing review and editing final draft.

