## supplementary Information for "Structures of the human transcription factor brachyury offer insights into DNA recognition, and identify small molecule binders for the development of degraders for cancer therapy"

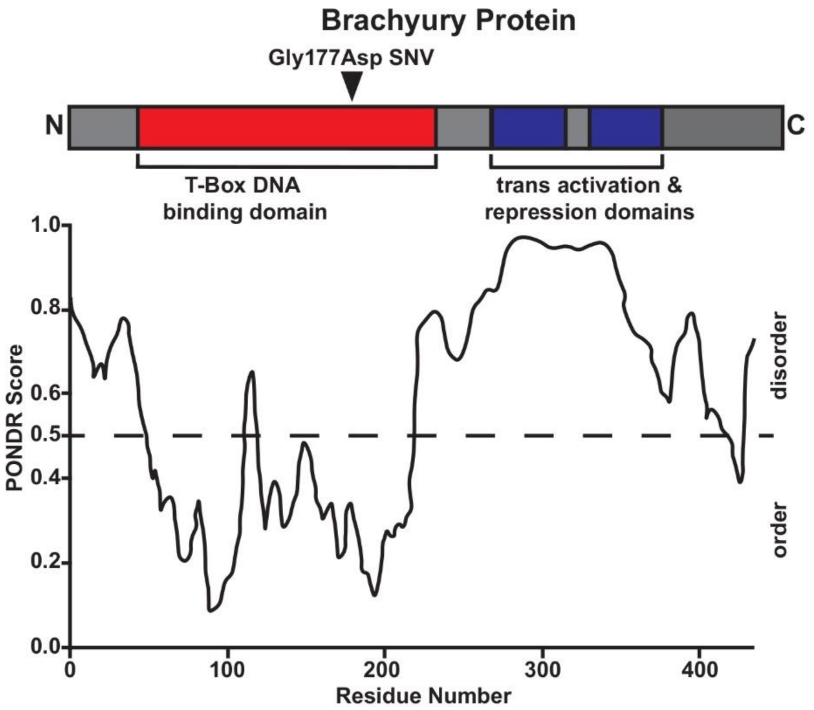


Figure S1 – Analysis of structural disorder in the full length brachyury protein using the PONDR method. Predicted disordered regions map well with the boundaries of the know DNA binding domain and indicate that brachyury is largely disordered outside this domain.

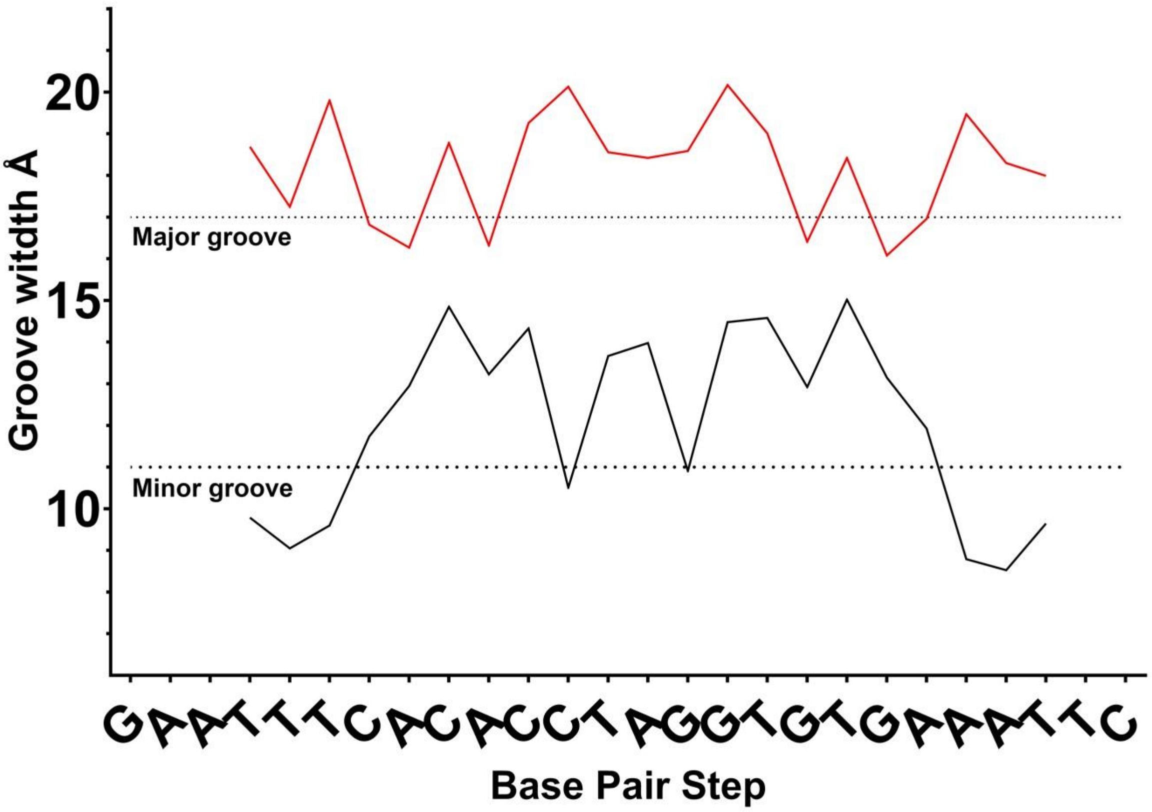


Figure S2 Plot of the measured major and minor groove widths of the DNA in the palindromic brachyury G177D DNA complex (6F59). Major groove widths are measured phosphorus to phosphorus between each base and its partner 3 bases downstream. Minor groove widths are measured phosphorus to phosphorus between each base and its partner 4 bases upstream.

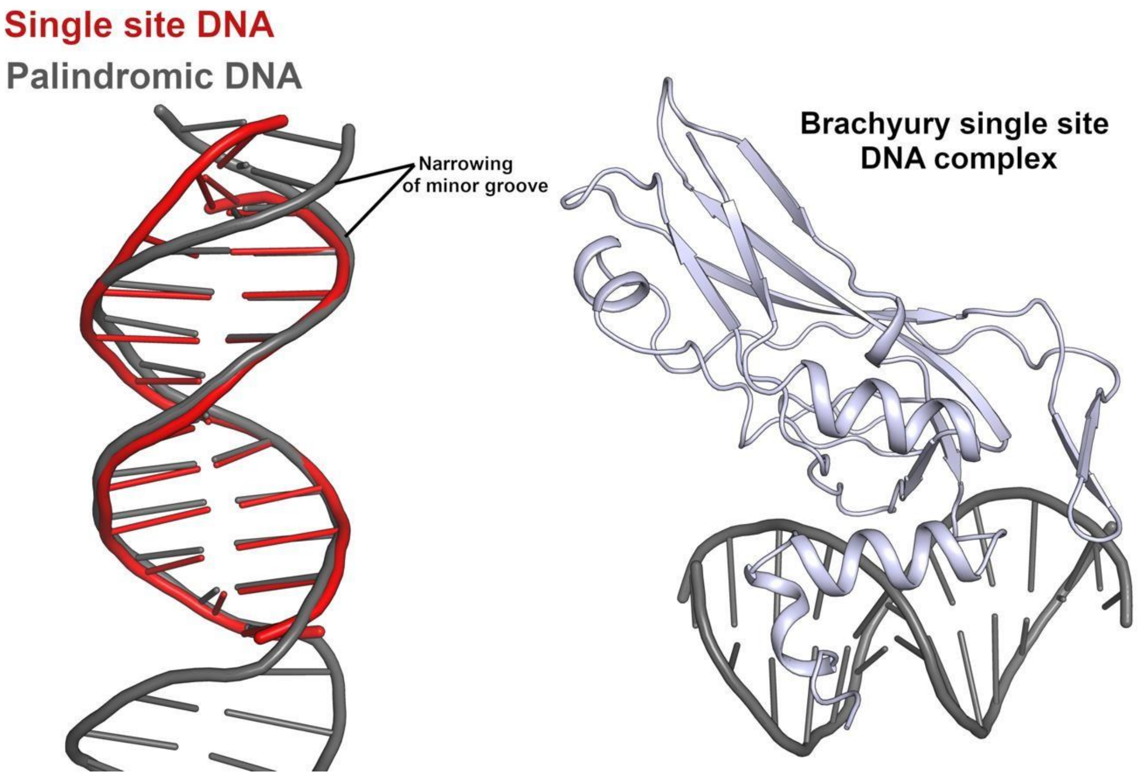


Figure S3 – Structure of WT brachyury in complex with a single site DNA. The right-hand panel shows the overall structure in cartoon form which is very similar to a half site of the palindromic DNA complex. The left-hand panel shows a comparison of the two DNA structures which contain similar distortions from canonical B-form DNA with the exception of the end narrowing of the minor groove which is not present in the single site DNA complex.

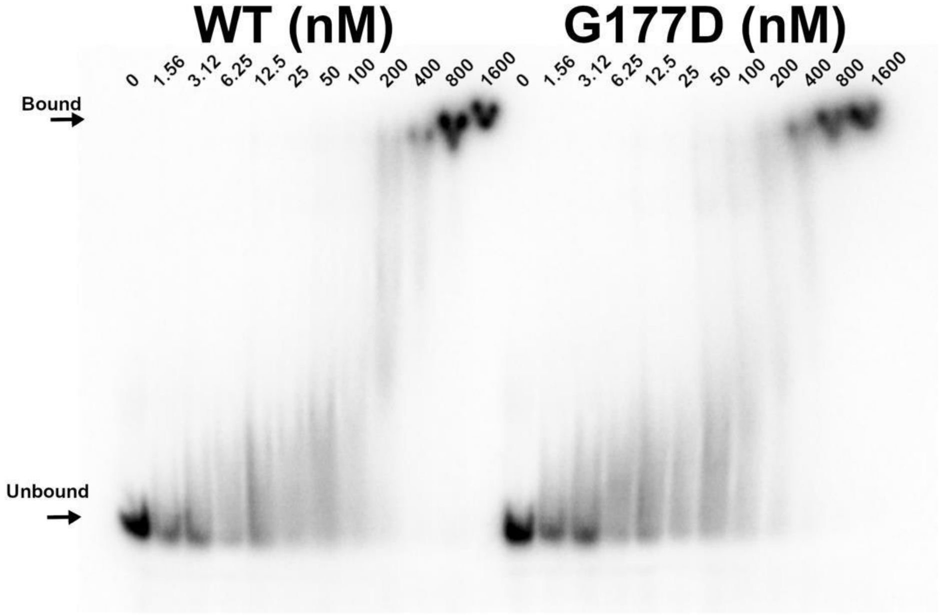


Figure S4 – Representative gel of full-length WT and G177D brachyury binding to an oligonucleotide containing a single T-box binding element. The results of the quantification of three replicates of this gel are shown in Figure 2c.

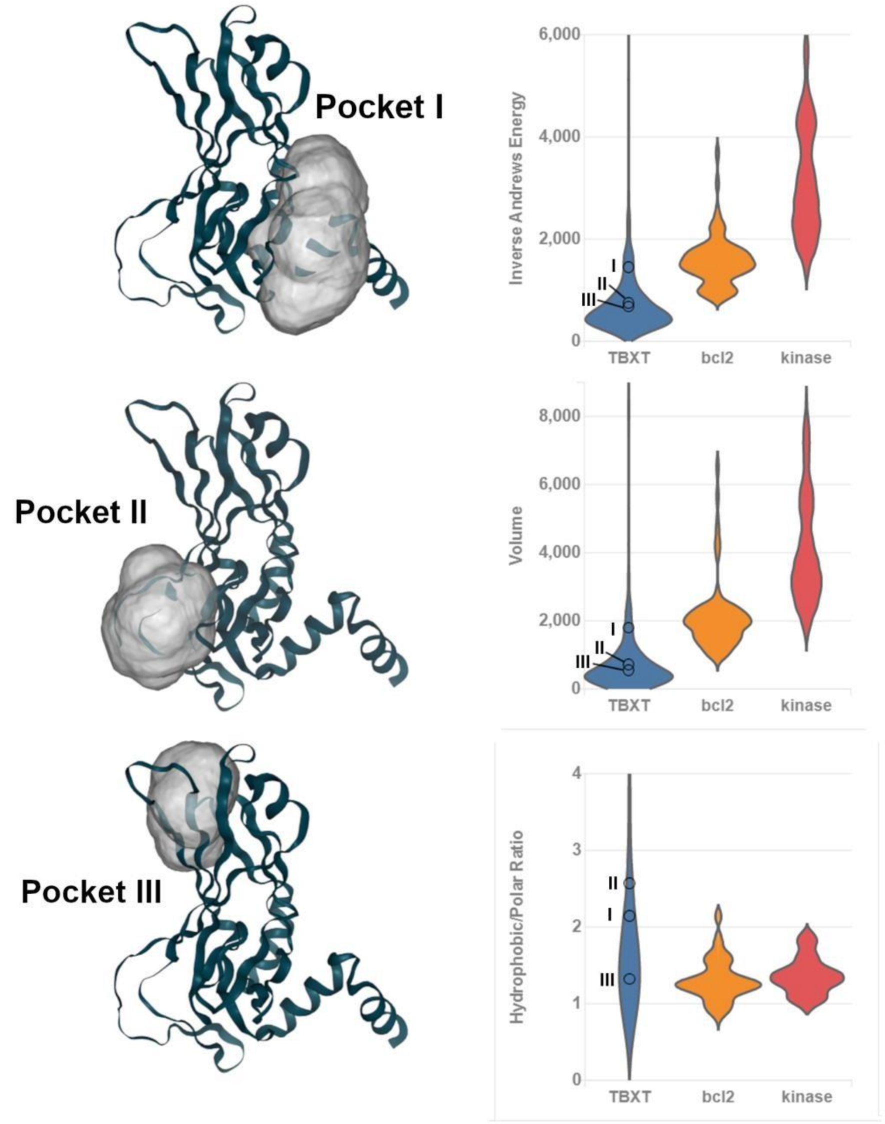


Figure S5 – Analysis of the ligandability potential of pockets on brachyury using the tools available at the CANSAR web server. The left panel shows representations of three pockets identified in the brachyury G177D DNA complex structure (6f59:chainA). The graphs on the right show pocket properties relating to ligandability with the brachyury pockets marked. Representative distributions of challenging (bcl2 – orange) and tractable (kinase – Red) pockets are shown for reference.

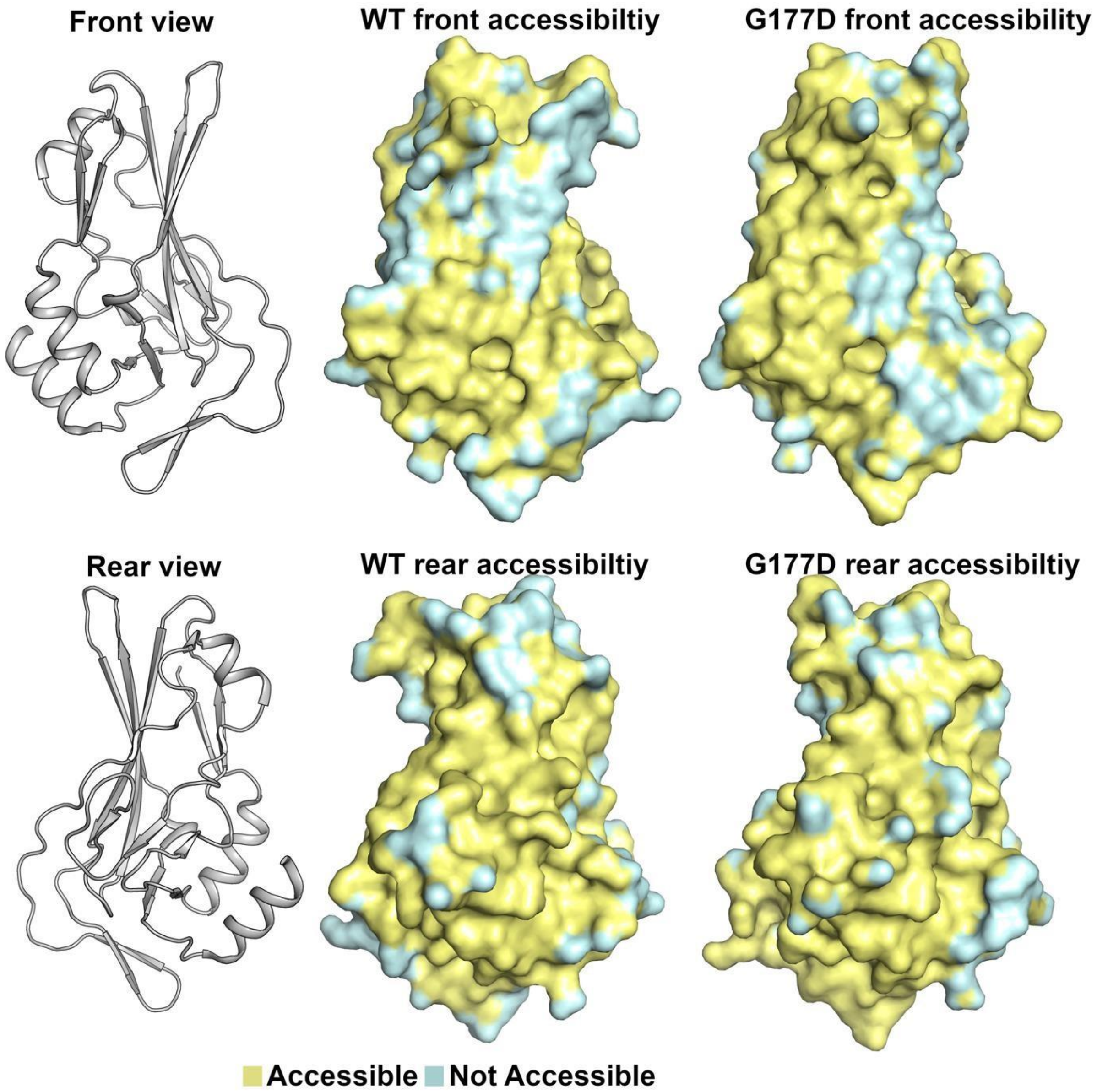


Figure S6 – Accessibility of the various surfaces and pockets in the WT and G177D crystals. Inaccessible regions are defined as surface atoms within 4 Å of an atom in a crystal contact.

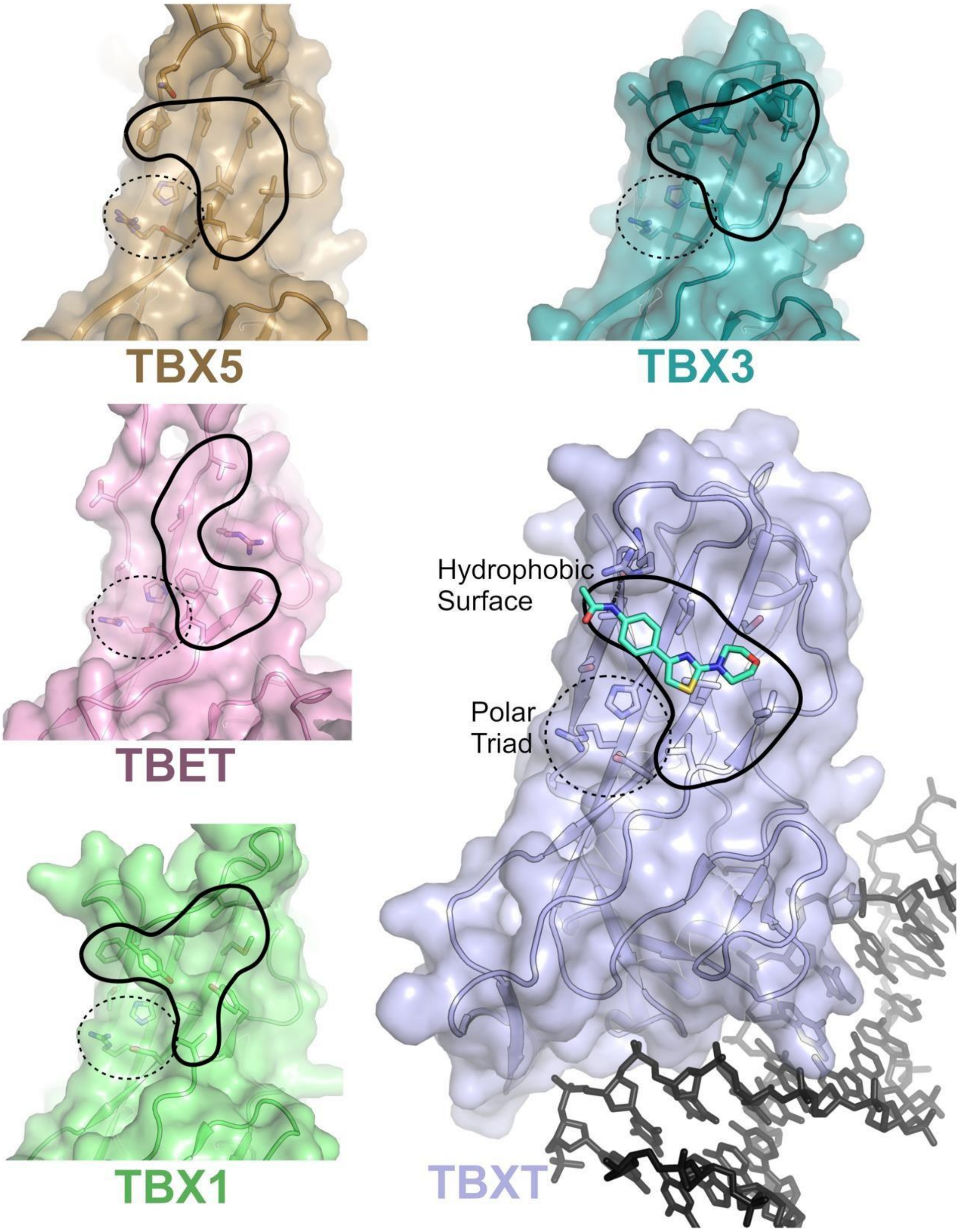


Figure S7 – Overview of pocket A’ in brachyury and other T-Box family members. The TBXT A’ pocket (shown in grey with surface view and ligand for reference) is distant from the DNA binding interface and contains a polar triad and an unusual cluster of surface exposed hydrophobic residues. Both of these features are conserved across the T-Box family.

| PDB | Ligand | Pocket | 2Fo-Fc map (1σ) | Location |
|---|---|---|---|---|
| 5QT0 | 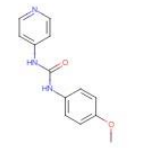 Z321318226 | 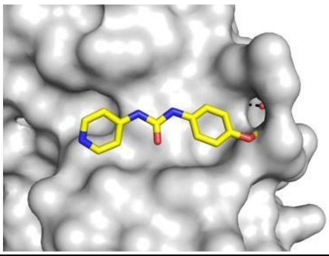 | 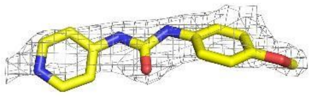 2.10 Å | 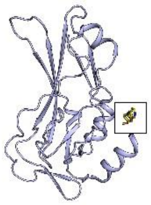 Pocket C |
| 5QS6 | 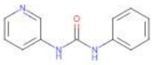 Z44592329 | 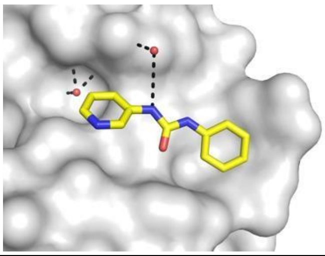 | 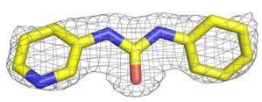 1.67 Å | 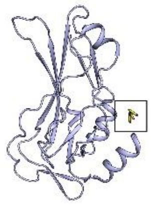 Pocket C |
| 5QS7 | 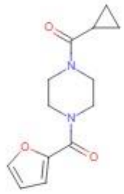 Z32327641 | 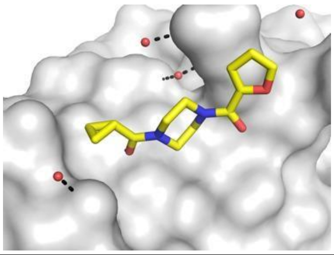 | 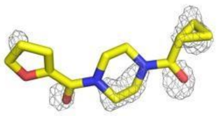 1.66 Å | 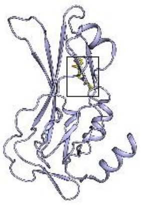 G177D other |
| 5QS8 | 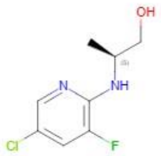 Z1432018343 | 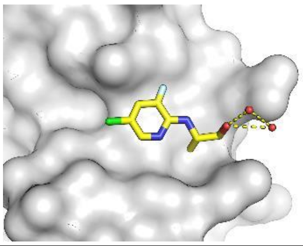 | 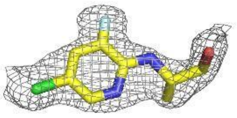 1.47 Å | 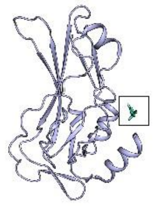 Pocket C |
| 5QS9 | 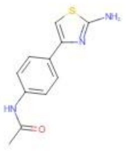 Z48847594 | 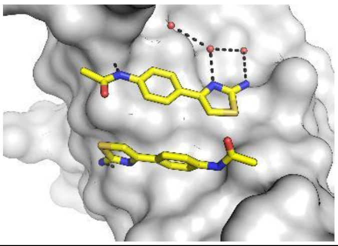 | 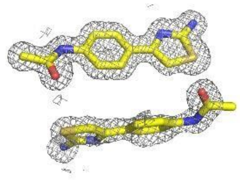 1.43 Å | 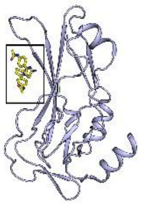 Pocket A’ |
| 5QSA | 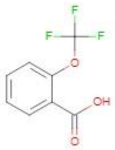 Z2856434778 | 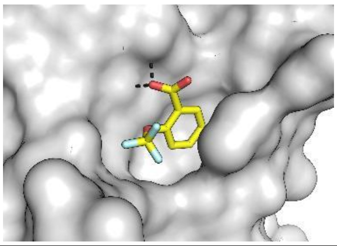 | 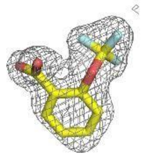 1.55 Å | 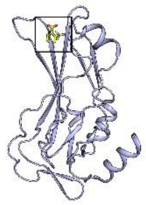 Pocket D |

| PDB | Ligand | Pocket | 2Fo-Fc map (1σ) | Location |
|---|---|---|---|---|
| 5QSB | 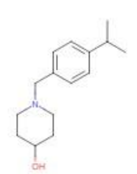 Z2856434874 | 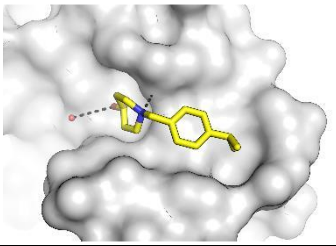 | 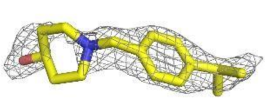 1.82 Å | 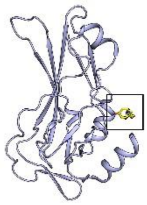 Pocket C |
| 5QSC | 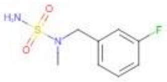 Z300245038 | 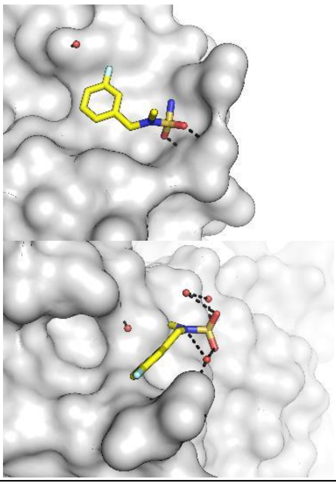 | 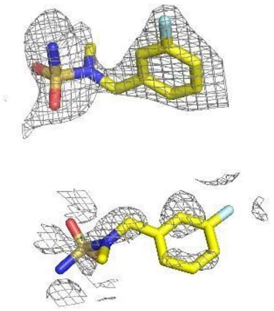 1.62 Å | 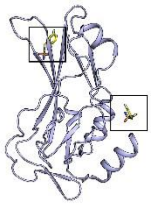 Pocket C & Pocket D |
| 5QSD | 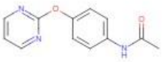 Z54571979 | 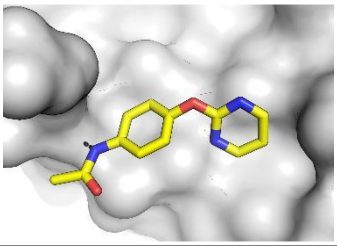 | 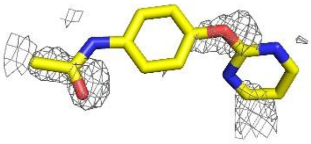 1.87 Å | 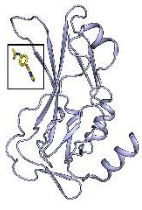 Pocket A' |
| 5QSE | 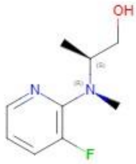 Z2017168803 | 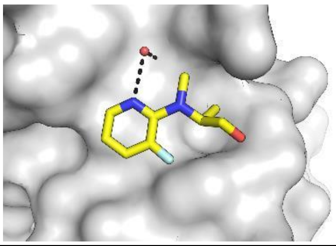 | 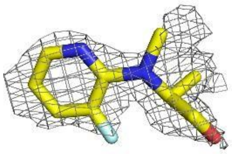 2.01 Å | 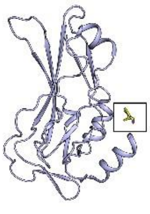 Pocket C |
| 5QSF | 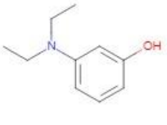 Z2856434814 | 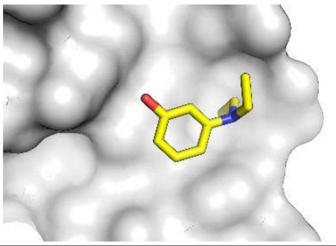 | 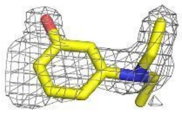 1.96 Å | 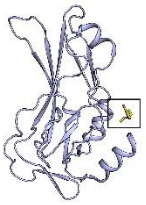 Pocket C |

| PDB | Ligand | Pocket | 2Fo-Fc map (1σ) | Location |
|---|---|---|---|---|

| PDB | Ligand | Pocket | 2Fo–Fc map (1σ) | Location |
|---|---|---|---|---|
| 5QSG | 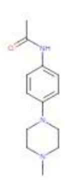 Z2856434903 | 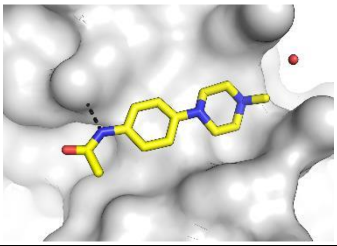 | 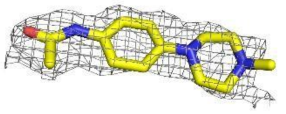 1.87 Å | 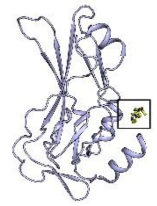 Pocket C |
| 5QSH | 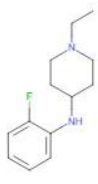 Z2856434868 | 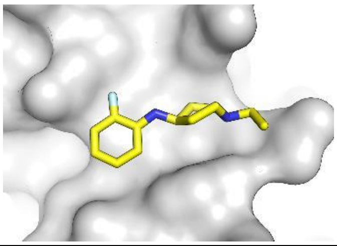 | 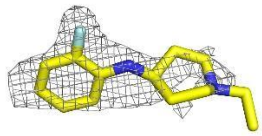 1.90 Å | 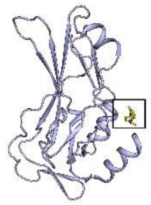 Pocket C |
| 5QSI | 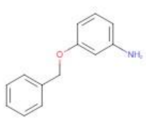 Z933326822 | 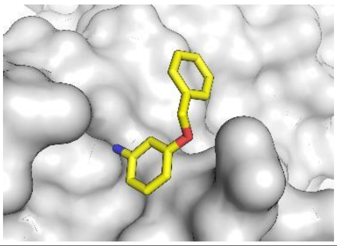 | 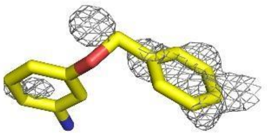 1.64 Å | 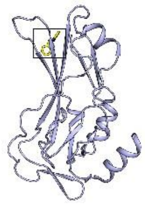 Pocket D |
| 5QSJ | 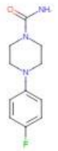 Z198194394 | 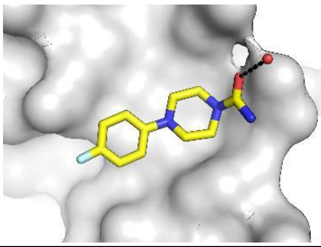 | 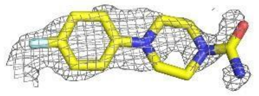 1.49 Å | 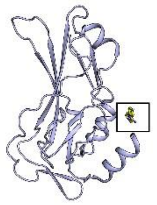 Pocket C |
| 5QSK | 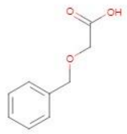 Z2856434906 | 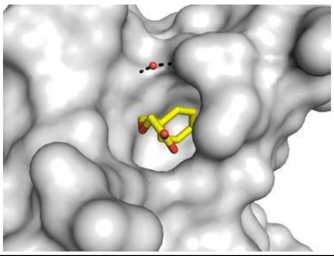 | 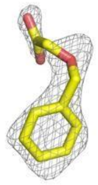 1.55 Å | 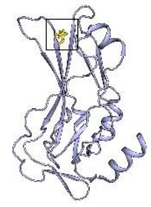 Pocket D |
| 5QSL | 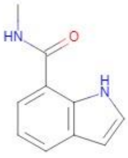 Z1273312153 | 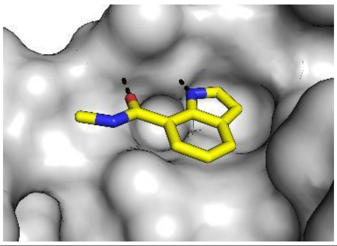 | 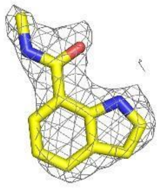 2.20 Å | 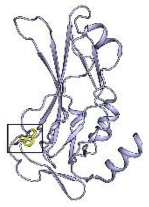 Pocket B |

| | | | | |
|---|---|---|---|---|
| 5QRF | 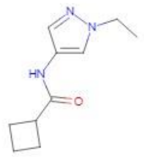 Z373768900 | 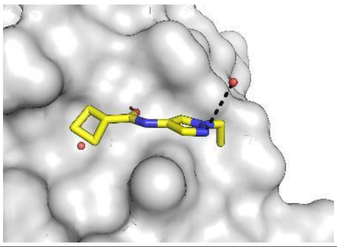 | 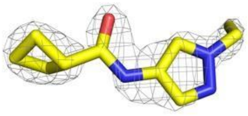 2.03 Å | 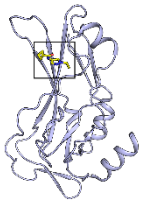 Pocket A |
| 5QRG | 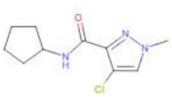 Z275151340 | 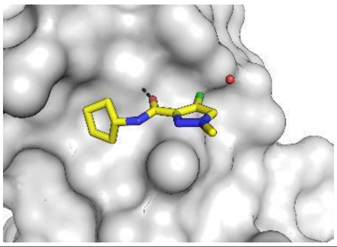 | 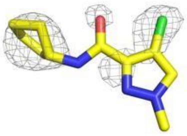 1.95 Å | 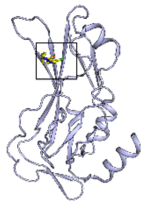 Pocket A |
| 5QRH | 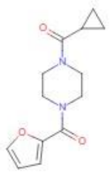 Z32327641 | 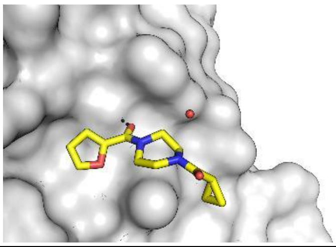 | 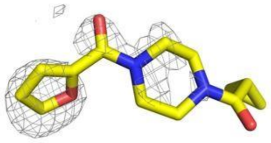 1.81 Å | 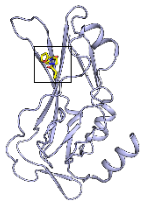 Pocket A |
| 5QRI | 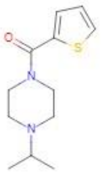 Z2856434826 | 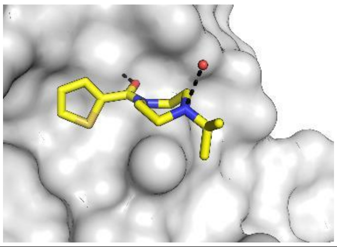 | 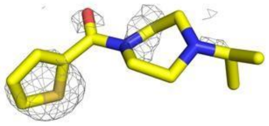 1.83 Å | 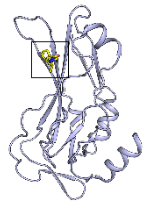 Pocket A |
| 5QRJ | 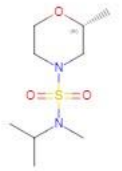 Z416341642 | 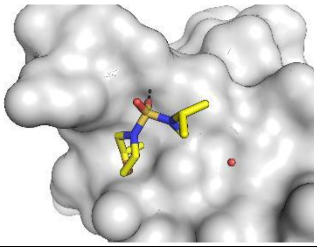 | 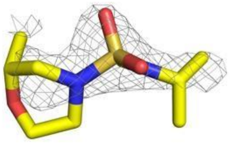 1.81 Å | 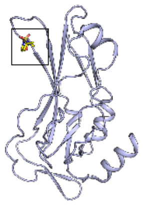 Pocket A |
| 5QRK | 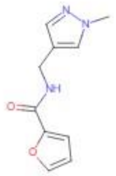 Z275179758 | 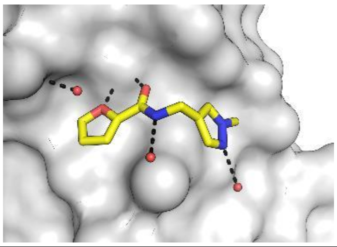 | 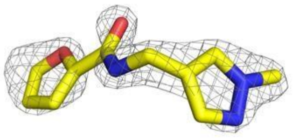 1.63 Å | 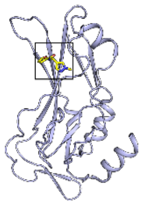 Pocket A |

| PDB | Ligand | Pocket | 2Fo-Fc map (1σ) | Location |
|---|---|---|---|---|
| 5QRL | 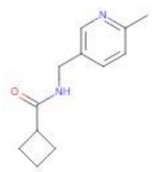 Z437516460 | 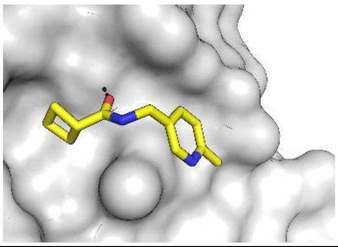 | 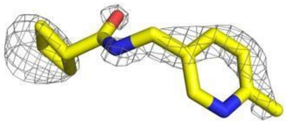 1.76 Å | 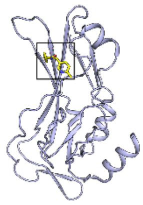 Pocket A |
| 5QRM | 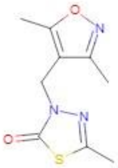 Z1899842917 | 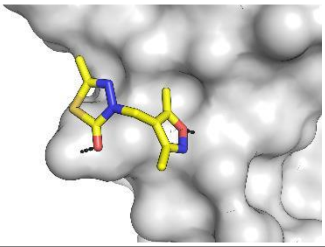 | 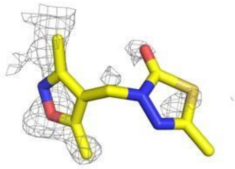 1.55 Å | 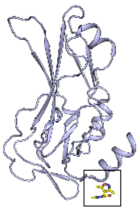 DNA Interface |
| 5QRN | 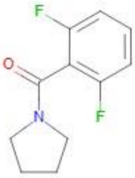 Z54226006 | 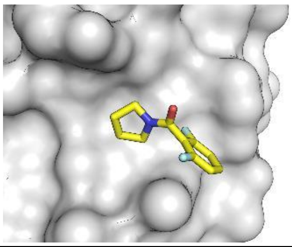 | 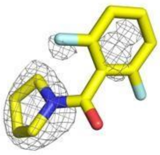 1.62 Å | 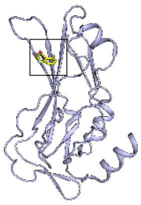 Pocket A |
| 5QRO | 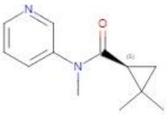 Z1506050651 | 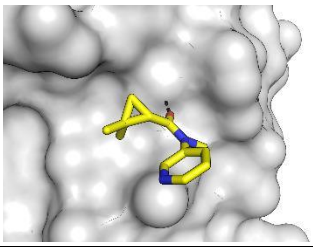 | 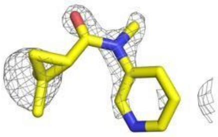 1.61 Å | 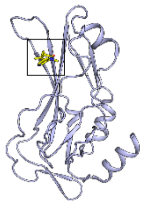 Pocket A |
| 5QRP | 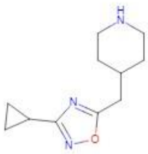 Z2442270563 | 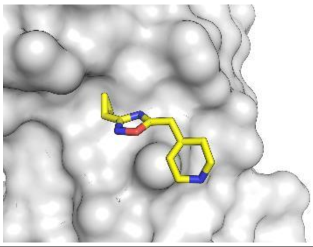 | 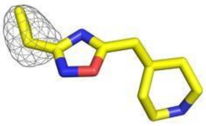 1.67 Å | 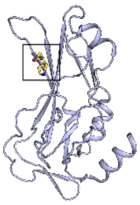 Pocket A |

| | | | | |
|---|---|---|---|---|
| 5QRQ | 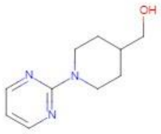 Z645232558 | 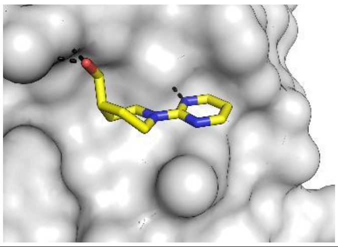 | 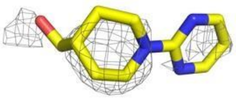 2.10 Å | 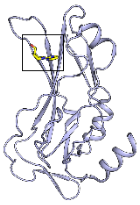 Pocket A |

| PDB | Ligand | Pocket | 2Fo-Fc map (1σ) | Location |
|---|---|---|---|---|
| 5QRR | 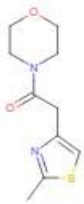 Z31720228 | 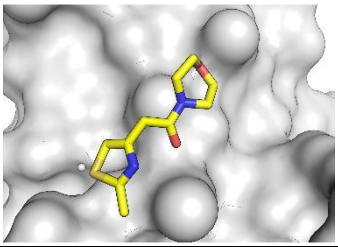 | 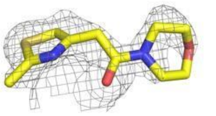 1.69 Å | 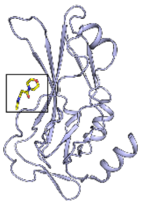 Pocket Other |
| 5QRS | 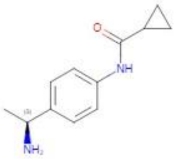 Z1891773393 | 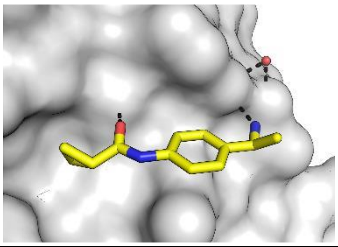 | 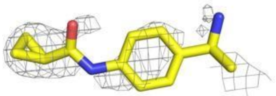 2.06 Å | 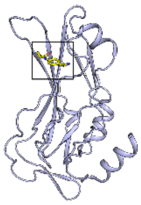 Pocket A |
| 5QRT | 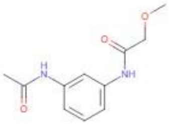 Z31735562 | 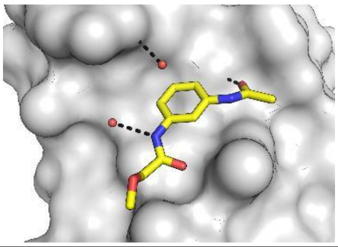 | 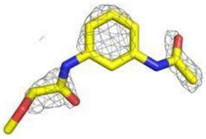 1.77 Å | 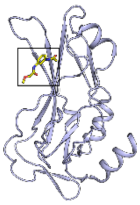 Pocket A |
| 5QRU | 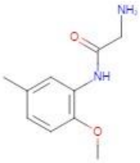 Z235341991 | 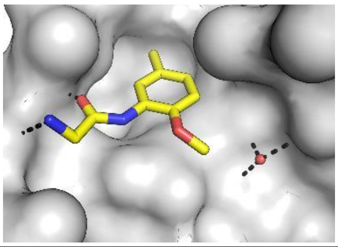 | 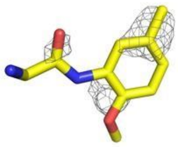 1.76 Å | 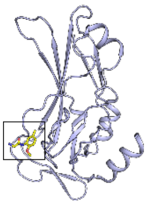 Pocket B |

| | | | | |
|---|---|---|---|---|
| 5QRV | 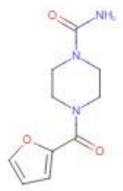 Z198194396 | 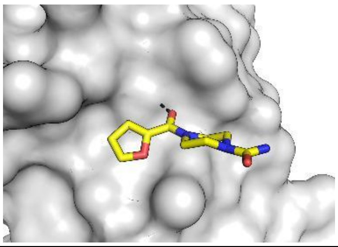 | 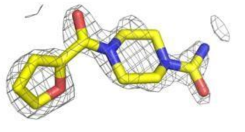 1.67 Å | 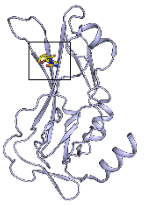 Pocket A |
| 5QRW | 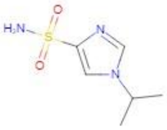 Z1509882419 | 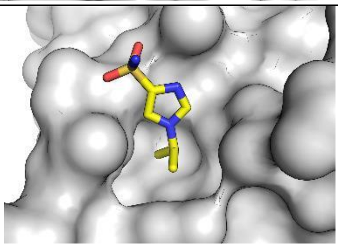 | 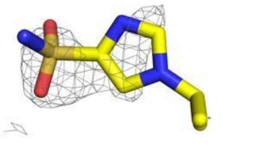 | 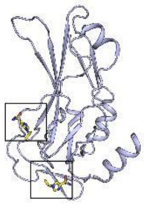 |

| PDB | Ligand | Pocket | 2Fo-Fc map (1σ) | Location |
|---|---|---|---|---|
| | | 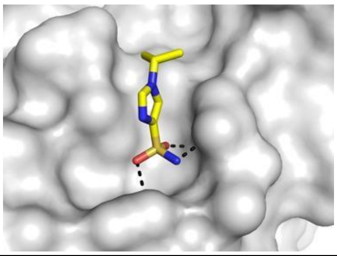 | 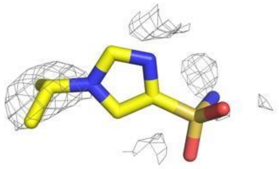 1.74 Å | Pocket B & DNA interface |
| 5QRX | 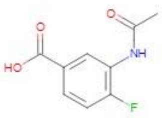 Z364328788 | 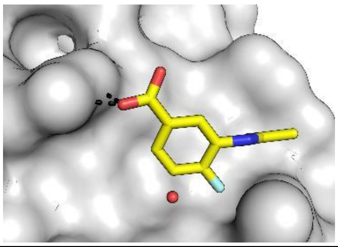 | 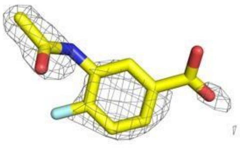 1.87 Å | 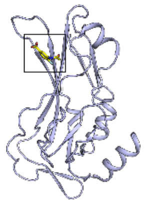 Pocket A |
| 5QRY | 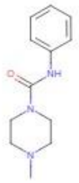 Z2856434890 | 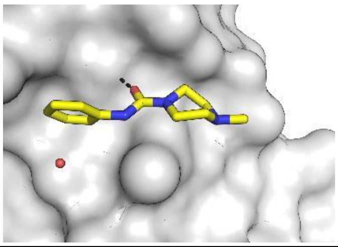 | 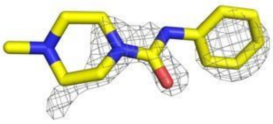 1.58 Å | 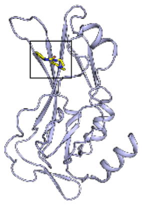 Pocket A |

| | | | | |
|---|---|---|---|---|
| 5QRZ | 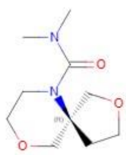 Z1998104358 | 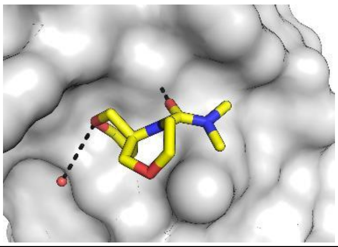 | 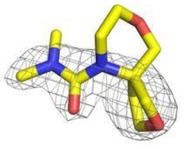 1.96 Å | 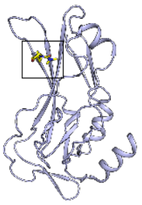 Pocket A |
| 5QS0 | 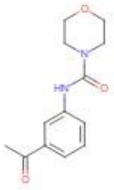 Z274555794 | 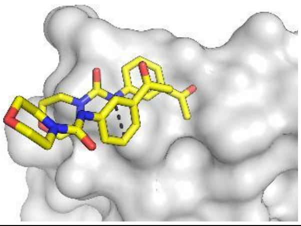 | 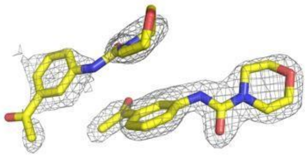 1.60 Å | 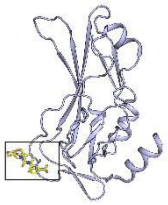 Other & Other |
| 5QS1 | 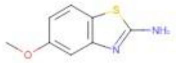 Z1954800564 | 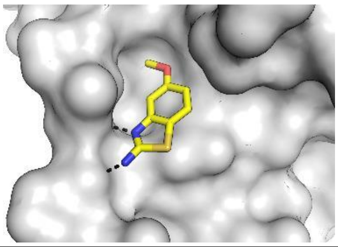 | 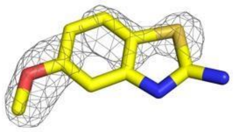 1.66 Å | 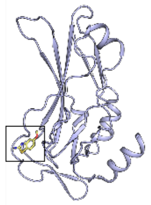 Pocket B |
| PDB | Ligand | Pocket | 2Fo-Fc map (1σ) | Location |
| 5QS2 | 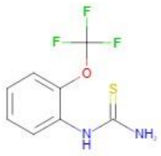 Z291279160 | 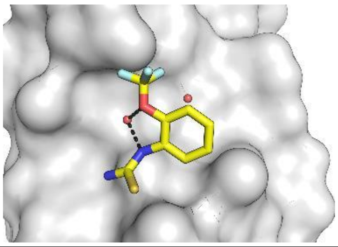 | 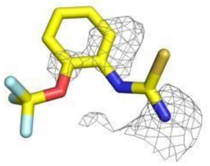 1.68 Å | 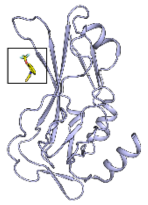 Pocket A |
| 5QS3 | 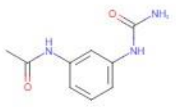 Z198195770 | 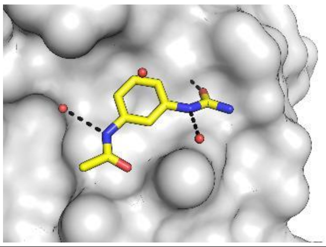 | 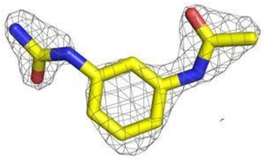 1.71 Å | 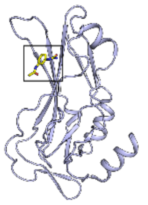 Pocket A |
| 5QS4 | 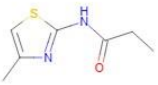 Z30820160 | 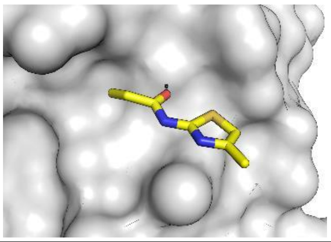 | 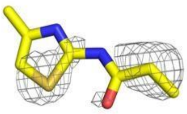 1.65 Å | 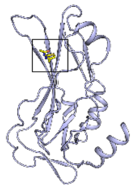 Pocket A |

| | | | | |
|---|---|---|---|---|
| 5QS5 | 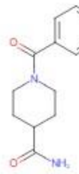 Z32400357 | 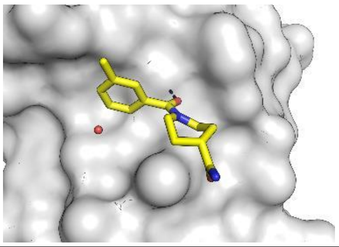 | 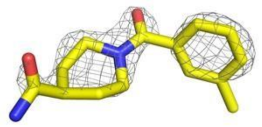 1.81 Å | 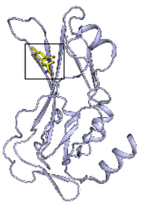 Pocket A |

Table S1 – Overview of the brachyury fragment screening hits. Each PDB entry is listed together with the chemical structure of the fragment, the fragment binding pocket, location, and electron density map. Maps shown are 2Fo-1Fc type calculated with the final refined coordinates and are shown to allow comparison of the relative strengths of the electron density for each fragment hit rather than the experimental evidence for fragment binding which is based on the PANDDA event maps.

| Entry | Structure | $K_D$ μM | Curve fit | Sensogram |
|---|---|---|---|---|
| 1a* | 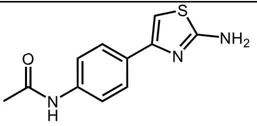 CSC000256427 | – | – | – |
| 1b* | 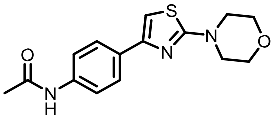 CSC02789850 | 316 | 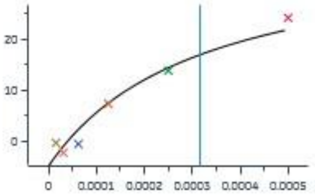 | 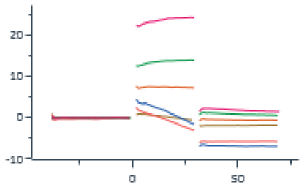 |
| 2 2-071 | 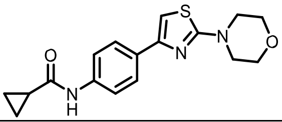 | 19 | 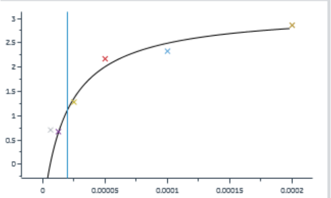 | 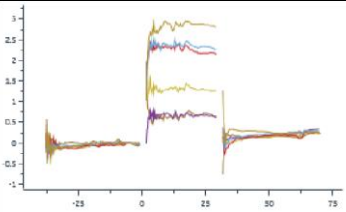 |
| 3 166-3 | 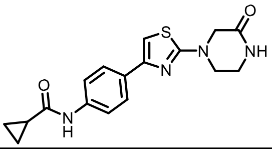 | 14 | 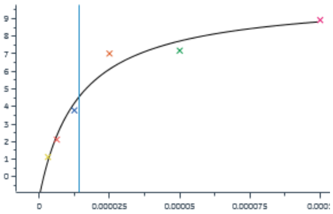 | 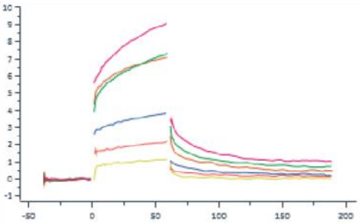 |
| 4 194-12 | 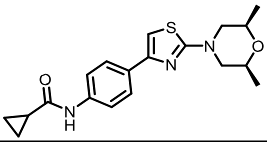 | 21 | 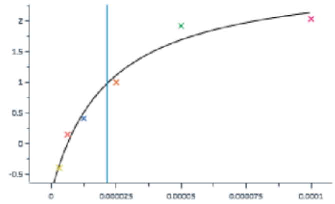 | 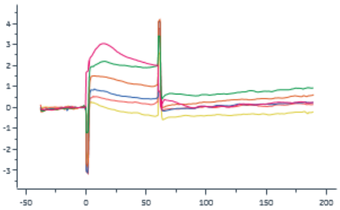 |
| 5 0290A | 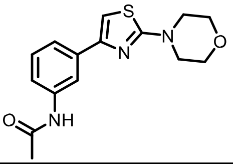 | 84 | 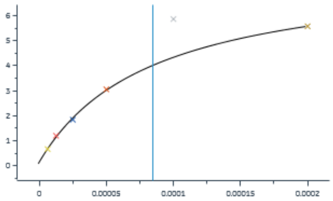 | 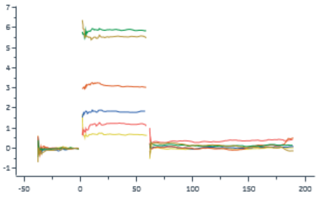 |
| 6 3007H | 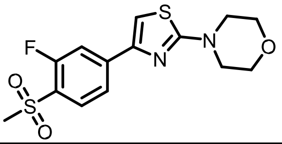 | 25 | 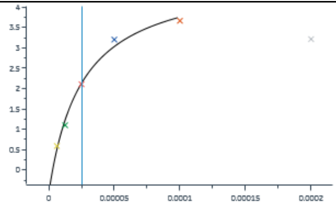 | 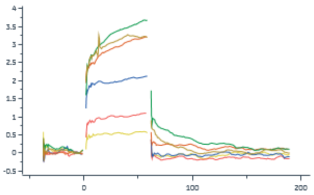 |
| 7 | 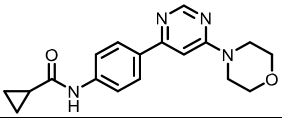 | 14 | 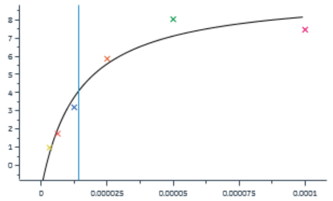 | 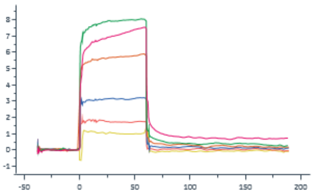 |

| | | | | |
|---|---|---|---|---|
| 8 | 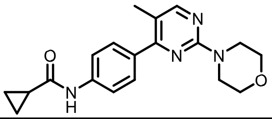 | 16 | 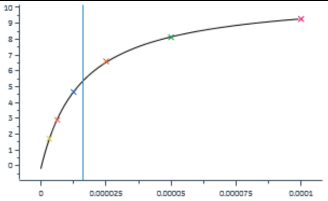 | 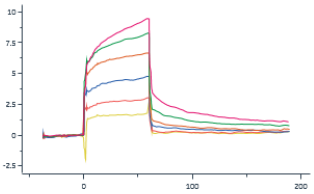 |
| 9 | 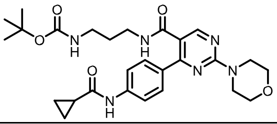 | 30 | 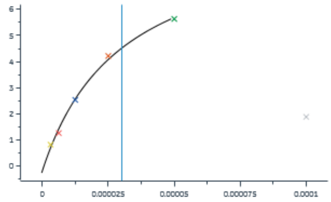 | 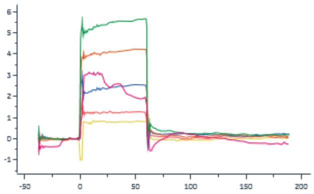 |
| 10 | 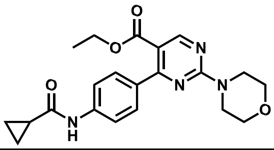 | 8 | 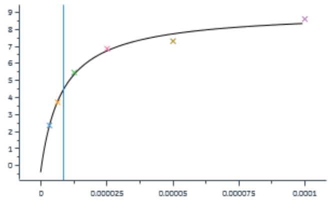 | 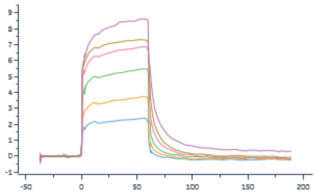 |
| 11 | 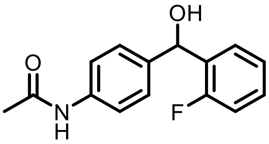 | 34<br>0 | 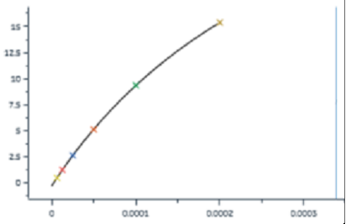 | 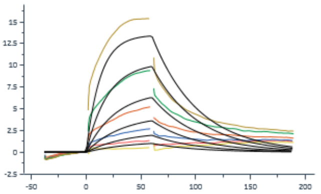 |
| 12 | 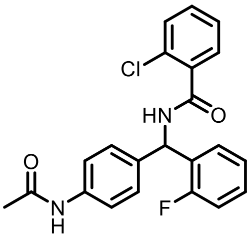 | 49 | 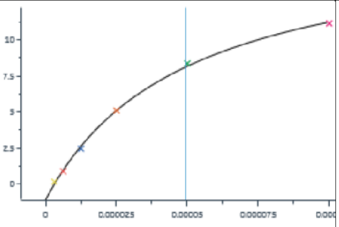 | 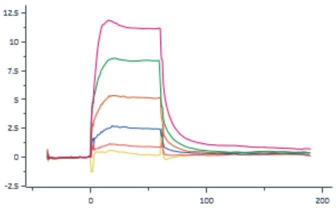 |
| 13 | 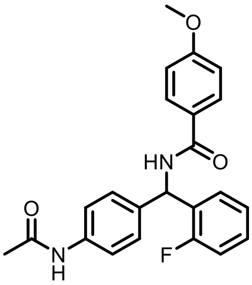 | 17 | 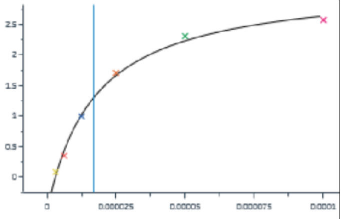 | 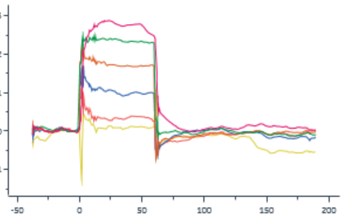 |

Table S2 – Summary of compounds binding to brachyury based on elaboration of fragment hits targeting pocket A’. SPR dose response fits are shown in the 4th column with response in RU as the Y axis and molar compound concentration as the X axis. The vertical line depicts the apparent dissociation constant using a dose response fit. Sensograms are shown in the 5th column plotted as response in RU over time in seconds.

General Chemistry Information

All reagents and solvents, unless specifically stated, were used as obtained from their commercial sources without further purification. Air and moisture sensitive reactions were performed under an inert atmosphere using nitrogen or argon in a previously oven-dried or flame-dried reaction flask, and addition of reagents were done using a syringe. All microwave (MW) reactions were carried out in a Biotage Initiator EXP US 400W microwave synthesizer. Thin layer chromatography (TLC) analyses were performed using 200 µm precoated sorbtech fluorescent TLC plates and spots were visualized using UV light. High resolution mass spectrometry samples were analyzed with a ThermoFisher Q Exactive HF-X (ThermoFisher, Bremen, Germany) mass spectrometer coupled with a H2Os Acquity H-class liquid chromatograph system. Column chromatography was undertaken with a Biotage Isolera One instrument. HPLC: Phenomenex Luna Phenyl-Hexyl (5 µm particle size, 100 Å pore size, 75 x 30 mm) column on an Agilent 1100 Series instrument equipped with an Agilent G1315B diode array detector measured at 220/254 nm. Nuclear magnetic resonance (NMR) spectrometry was run on a varian Inova 400 MHz or Bruker Avance III 700 MHz spectrometer equipped with a TCI HC/N-D 5 mm cryoprobe and data was processed using the MestReNova processor. Chemical shifts are reported in ppm with residual solvent peaks referenced as internal standard.

**Synthesis of *N*-(4-(2-morpholinothiazol-4-yl)phenyl)cyclopropanecarboxamide** (2)

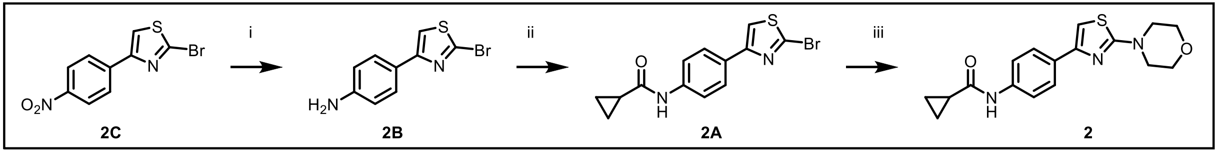


**Reaction Scheme S1**: i. hydrazine hydrate, EtOH, Raney N (excess), 50 °C, 2 hr, 86%; ii. cyclopropyl acyl chloride, pyridine, 0 °C, 24 hr, 54%; iii. Morpholine, DMA, 140 °C, 16 h, 65%.

**Synthesis of 4-(2-bromothiazol-4-yl)aniline (2B)**: To a suspension of 2-bromo-4-(4-nitrophenyl)thiazole (400 mg, 1.00 Eq, 1.40 mmol) in EtOH (30 mL) was added hydrazinehydrate (1.05 g, 1.02 mL, 15.00 Eq, 21.0 mmol) and the reaction mixture was heated to 50 °C for 15 min until the starting material was completely dissolved. Excess Raney Nickel (823 mg, 112 µL, 10.00 Eq, 14.0 mmol) was added to the reaction mixture and the reaction was maintained at 50 °C. When gas evolution ceased the reaction mixture was filtered through celite and concentrated to dryness to yield the desired product **2B** (310 mg, 86%) which was taken to the next step with no further purifications.

$^{1}H$ NMR (850 MHz, DMSO) δ 7.71 (s, 1H), 7.59 – 7.55 (m, 2H), 6.62 – 6.58 (m, 2H), 5.38 (s, 2H).
$^{13}C$ NMR (214 MHz, DMSO) δ 158.8, 152.48, 138.3, 130.3, 124.1, 116.9, 116.6.

**Synthesis of N-(4-(2-bromothiazol-4-yl)phenyl)cyclopropanecarboxamide (2A)**: To a 100 mL RBF was added 4-(2-bromothiazol-4-yl)aniline (350 mg, 1.0 eq, 1.37 mmol) and 20 mL of DCM. The reaction mixture was cooled to 0 °C using an ice bath and pyridine (217 mg, 2.74 mmol, 2.00 eq.) was added to the reaction mixture followed by cyclopropyl acyl chloride (172 mg, 1.65 mmol, 1.20 eq) dropwise over 30 min. The reaction mixture was stirred for 3.0-24 h at room temperature. The reaction mixture was poured into a separatory funnel and washed 3 x 10.0 mL of saturated $NaHCO_3$(aq) and

15.0 mL brine. The organic layer was dried over anhydrous $Na_2SO_4$ filtered and evaporated. The crude material waspurified by column chromatography on silica gel (30-50% Ethyl acetate in hexanes) to afford the product **2A** (240 mg, yield 54%, 97% pure).

$^1$H NMR (700 MHz, DMSO) δ 10.26 (s, 1H), 7.95 (s, 1H), 7.79 – 7.75 (m, 2H), 7.61 (d, J = 8.7 Hz, 2H), 1.73 (tt, J = 7.4, 4.9 Hz, 1H), 0.75 (ddd, J = 9.1, 6.1, 2.3 Hz, 4H).

$^{13}$C NMR (176 MHz, DMSO) δ 172.2, 154.8, 140.1, 136.3, 128.1, 126.9, 119.5, 117.6, 60.2, 15.1, 14.6, 7.8.

**Synthesis of N-(4-(2-morpholinothiazol-4-yl)phenyl)cyclopropanecarboxamide (2)**: To a small 4 mL dram vial was added N-(4-(2-bromothiazol-4-yl)phenyl)cyclopropanecarboxamide **2** (30 mg, 1 Eq, 93 μmol), to this was added morpholine (16 mg, 2 eq. 190 μmol) and DMA (0.5 mL). The reaction mixture was then heated to 140 °C overnight. After heating for 16 h the reaction mixture was taken to dryness and purified by HPLC 0.05% TFA in $H_2O$/MeOH. Product fractions were collected and dried to yield the product **2** (20 mg, yield 65%, 98% pure), m/z: 330.12722 [M+H]+.

$^1$H NMR (700 MHz, DMSO) δ 10.25 (s, 1H), 7.80 – 7.76 (m, 2H), 7.61 (d, J = 8.3 Hz, 2H), 7.19 (d, J = 1.3 Hz, 1H), 3.74 (t, J = 4.9 Hz, 4H), 3.44 (t, J = 4.8 Hz, 4H), 1.79 (s, 1H), 0.83 – 0.77 (m, 4H).

$^{13}$C NMR (101 MHz, cdcl$_3$) δ 129.1, 127.2, 126.7, 102.6, 77.3, 77.0, 76.7, 66.2, 16.2, 14.8, 11.1, 11.0, 8.0, 7.4.

**Synthesis of N-(4-(2-(3-oxopiperazin-1-yl)thiazol-4-yl)phenyl)cyclopropanecarboxamide (3)**: To a small 4 mL dram vial was added N-(4-(2-bromothiazol-4yl)phenyl)cyclopropanecarboxamide, **2A** (30 mg, 1 eq., 93 µmol), to this was added piperazin-2-one (28 mg, 3 eq., 280 µmol) and DMA (0.5 mL). The reaction mixture was then heated to 140 °C overnight. After heating for 16 h the reaction mixture was taken to dryness and purified by HPLC 0.05% TFA in $H_2O$/MeOH, product fractions were collected and dried to yield the product **3** (9.0 mg, yield 28%, 97% pure), m/z: 343.112 $[M+H]^+$.

$^1$H NMR (850 MHz, DMSO) δ 10.25 (s, 1H), 8.19 (t, J = 2.8 Hz, 1H), 7.81 – 7.77 (m, 2H), 7.64 – 7.59 (m, 2H), 7.19 (s, 1H), 4.01 (s, 2H), 3.70 – 3.66 (m, 2H), 3.37 (ddd, J = 7.0, 5.2, 2.9 Hz, 2H), 1.79 (tt, J = 7.8, 4.8 Hz, 1H), 0.80 (ddt, J = 12.8, 10.1, 4.0 Hz, 4H).

$^{13}$C NMR (214 MHz, DMSO) δ 174.7, 171.8, 169.1, 153.6, 142.0, 132.5, 129.3, 122.0, 104.6, 53.9, 40.6, 37.6, 24.5, 17.7, 10.30.

**Synthesis of N-(4-(2-((2R,6S)-2,6-dimethylmorpholino)thiazol-4-yl)phenyl)cyclopropanecarboxamide (4)**: To a small 4 mL dram vial was added N-(4-(2-bromothiazol-4-yl)phenyl)cyclopropanecarboxamide (30 mg, 1 Eq, 93 µmol), to this was added (2S,6R)-2,6-dimethylmorpholine (32 mg, 3 Eq. 280 µmol) and DMA (0.5 mL). The reaction mixture was then heated to 140 °C overnight. After heating for 16 h the reaction mixture was taken to dryness and purified by HPLC 0.05% TFA in $H_2O$/MeOH, product fractions were collected and dried to yield the product **4** (10.3 mg, yield 31%, 96% pure), m/z: 358.1584 [M+H]+.

$^1$H NMR (850 MHz, DMSO) δ 10.25 (s, 1H), 7.78 – 7.76 (m, 2H), 7.64 – 7.59 (m, 2H), 7.14 (s, 1H), 4.08 (pd, J = 6.4, 3.5 Hz, 2H), 3.55 (dd, J = 12.3, 3.5 Hz, 2H), 3.16 (dd, J = 12.3, 6.1 Hz, 2H), 1.20 (d, J = 6.5 Hz, 7H), 0.84 – 0.76 (m, 4H), 1.79 (tt, J = 7.7, 4.7 Hz, 1H).

$^{13}C$ NMR (214 MHz, DMSO) δ 174.7, 174.1, 153.5, 141.9, 132.6, 129.3, 122.0, 104.1, 68.1, 55.8, 20.69, 17.68, 10.29.

**Synthesis of N-(3-(2-morpholinothiazol-4-yl)phenyl)acetamide (5):**

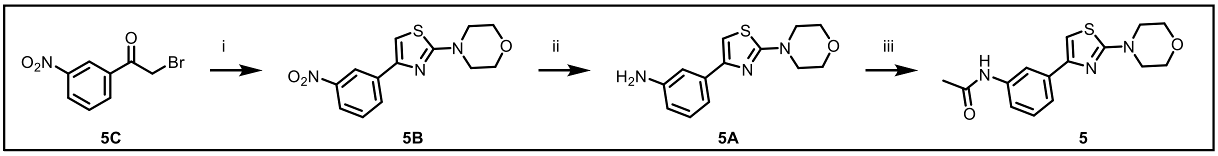


**Reaction Scheme S2**: i. thiourea, EtOH, reflux, 3 hr; ii. hydrazine hydrate, EtOH, Raney N (excess), 50 °C, 2 hr, 89%; iii. acetyl chloride, pyridine, THF, 0 °C to rt 2 hr then rt overnight, 82%.

**Synthesis of 4-(4-(3-nitrophenyl)thiazol-2-yl)morpholine (5B)**: To a 100 mL RBF was added thiourea (1.52 g, 20.0 mmol) and 2-bromo 4′-bromoacetophenone (5.56 g, 20.0 mmol) in EtOH (40 mL). The suspension was refluxed for 3 h. After filtration, the precipitate was washed with EtOH to get [4-(4-bromophenyl)-1,3-thiazol-2-yl]amino hydrobromide (1:1) as a solid, **5B** (6.05 g, yield 90%) which was used for the next step without purification. $^{1}H$ NMR (850 MHz, DMSO) δ 8.63 (q, J = 2.3 Hz, 1H), 8.29 (dt, J = 7.7, 1.3 Hz, 1H), 8.13 (ddd, J = 8.1, 2.4, 1.0 Hz, 1H), 7.68 (t, J = 8.0 Hz, 1H), 7.62 (s, 1H), 3.77 – 3.73 (m, 4H), 3.48 – 3.45 (m, 4H).

$^{13}C$ NMR (214 MHz, DMSO) δ 174.16, 151.09, 139.28, 135.04, 133.68, 125.28, 122.92, 108.55, 68.45, 50.98, 43.14.

**Synthesis of 3-(2-morpholinothiazol-4-yl)aniline (5A)**: In a 100 mL RBF was added 4-(4-(3-nitrophenyl)thiazol-2-yl)morpholine (2.5 g, 8.6 mmol, 1 eq.) in EtOH (50 mL) was added hydrazinehydrate (6.4 g, 6.3 mL, 15.00 Eq, 130 mmol) and the reaction mixture was heated to 50 °C for 15 min until the starting material dissolved completely. Excess of Raney Nickel (10 g, 20.00 Eq, 170 mmol) was added to the reaction mixture and the reaction was maintained at 50 °C. When gas evolution ceased, the reaction mixture was was filtered over celite and concentrated to give 3-(2morpholinothiazol-4-yl)aniline, **5A** (2.0 g, yield 89%, 96% purity) The product was taken to the next step with no further purifications.

**Synthesis of N-(3-(2-morpholinothiazol-4-yl)phenyl)acetamide (5)**: To a 100 mL RBF was added 3-(2-morpholinothiazol-4-yl)aniline (65.3 mg, 0.250 mmol, 1 eq.) in 5 mL of THF. The reaction mixture was cooled to 0 °C using an ice bath. To this was added pyridine (39.6 mg, 40.5 µL, 0.5 mmol, 2 eq.) followed by dropwise addition of acetyl chloride (23.5 mg, 0.30 mmol, 1.20 eq.) over 10 min. The reaction mixture was stirred at room temperature until complete by TLC. The reaction mixture was poured into a separatory funnel, washed 3 x 10.0 mL of saturated $NaHCO_3$(aq) and 15.0 mL brine. The organic layer was dried over $Na_2SO_4$, filtered and concentrated in vacuo. The crude product was purified by silica column chromatography (30%-50% Ethyl acetate in hexanes) to afford the product N-(3-(2-morpholinothiazol-4yl)phenyl)acetamide, **5** (62 mg, yield 82%, 96% purity), m/Z: 304.11139 $[M+H]^+$.

$^1H$ NMR (400 MHz, $CD_3OD$) δ 8.01 (t, J = 1.9 Hz, 1H), 7.55 (dt, J = 7.9, 1.4 Hz, 1H), 7.48 (ddd, J = 8.1, 2.2, 1.1 Hz, 1H), 7.29 (t, J = 7.9 Hz, 1H), 6.99 (s, 1H), 3.84 – 3.77 (m, 4H), 3.52 – 3.46 (m, 4H), 2.12 (s, 3H).

$^{13}C$ NMR (101 MHz, $CD_3OD$) δ 128.47, 121.41, 118.99, 117.48, 102.02, 65.80,
22.41.

**Synthesis of 4-(4-(3-fluoro-4-(methylsulfonyl)phenyl)thiazol-2-yl)morpholine (6)**: To a 4 mL microwave vial was added 4-(4-bromothiazol-2-yl)morpholine (50.00 mg, 1.00 Eq, 200.7 µmol), 3-Fluoro-4-(methylsulfonyl)phenylboronic acid (56.88 mg, 1.30 Eq, 260.9 µmol) and tetrakis (11.60 mg, 0.05 Eq, 10.03 µmol) ($Pd(P(Ph)_3)_4$ (5 mol%)] were dissolved in a mixed solution of 1,4-dioxane (3.0 mL)/$H_2O$ (0.3 mL) under argon. After degassing for 5 minutes, $Cs_2CO_3$ (98.09 mg, 1.50 Eq, 301.0 µmol) was added and the reaction mixture was heated in a microwave at 120 °C for 30 min.

The reaction mixture was allowed to cool to room temperature and reaction mixture was concentrated in vacuo. To the reaction mixture H2O was added and the solution was acidified to pH=2 using 2N HCl. The white solid precipitate was collected by filtration and dried to give the desired compound 4-(4-(3-fluoro-4-(methylsulfonyl)phenyl)thiazol-2-yl)morpholine, **6** (52.0 mg, yield 75.67%, purity >95%), m/Z: 343.05826 $[M+H]^+$.

$^1$H NMR (700 MHz, DMSO) δ 7.98 – 7.91 (m, 2H), 7.88 (t, J = 7.7 Hz, 1H), 7.72 (d, J = 1.3 Hz, 1H), 3.75 (t, J = 4.8 Hz, 4H), 3.48 (t, J = 4.8 Hz, 4H), 3.34 (s, 3H).

$^{13}$C NMR (214 MHz, CDCl3) δ 173.96, 163.06, 161.88, 151.23, 145.26, 145.22, 132.53, 129.30, 129.23, 124.41, 124.39, 117.24, 117.13, 108.33, 68.79, 68.68, 51.28, 50.61, 46.68, 46.66.

**Synthesis of N-(4-(6-morpholinopyrimidin-4-yl)phenyl)cyclopropanecarboxamide (7)**:

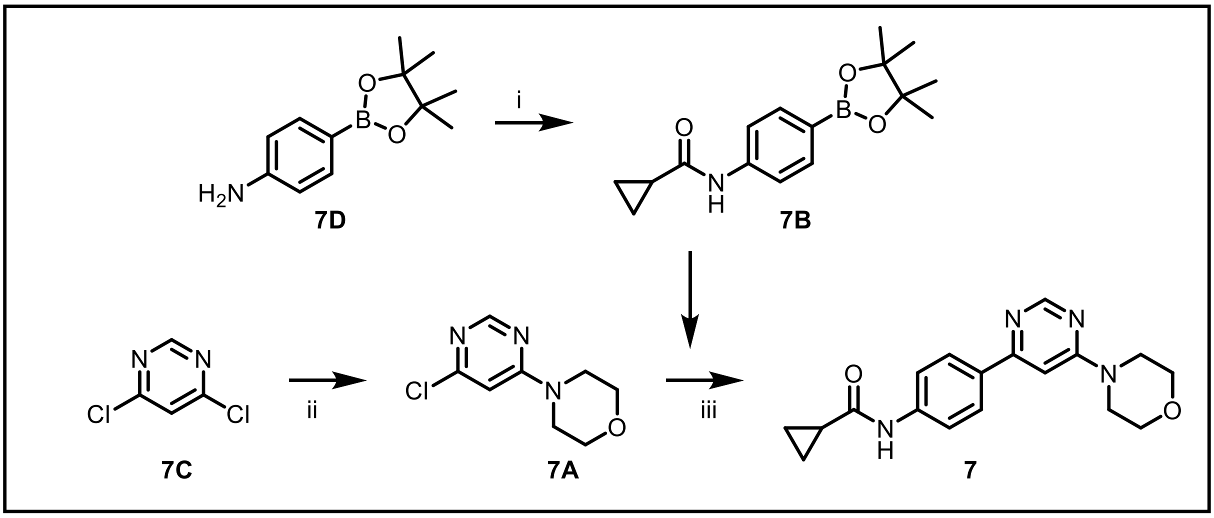


**Reaction scheme S3**: i. HATU, DIPEA, DMF, RT, 2 hr, 44%; ii. DIPEA, CAN, RT, 12 hr, 42%; iii. $PdCl_2(PPh_3)_2$, $K_2CO_3$, dioxane, water, 90 °C, 12 hr, 9%.

**Synthesis of N-(4-(4,4,5,5-tetramethyl-1,3,2-dioxaborolan-2-yl)phenyl)cyclopropanecarboxamide (7B)**: To a stirred solution of cyclopropane carboxylic acid (0.3 g, 3.48 mmol, 1.0 eq.) in DMF (2.0 mL) was added HATU (1.81 g, 5.23 mmol, 1.5 eq.) followed by addition of DIPEA (0.67

g, 5.23 mmol, 1.5 eq.) and stirred at room temperature for 10 min. To this reaction mixture was added 4-(4,4,5,5tetramethyl-1,3,2-dioxaborolan-2-yl)aniline (**7D**) (0.7 g, 3.48 mmol, 1.0 eq.) and stirred at room temperature for 2 h. Reaction progress was monitored by TLC, after completion, the reaction mixture was diluted with $H_2O$ (20 mL) and extracted with EtOAc (3 x 30 mL). The organic layer was washed with brine solution, dried over $Na_2SO_4$, filtered and concentrated to give crude compound which was purified by flash column chromatography using a 5-10% EtOAc/n-hexane. The product fractions were collected and dried to yield N-(4-(4,4,5,5-tetramethyl-1,3,2-dioxaborolan-2-yl)phenyl)cyclopropanecarboxamide (**7B**) (0.4 g, 44%) as an off white solid. m/z: 288.19 $[M+H]^+$ .

$^1$H NMR (400 MHz, DMSO-d6) δ 10.32 (s, 1H), 7.60 (s, 4H), 1.83 – 1.76 (m, 1H), 1.28 (s, 12H), 0.80 (d, J = 5.0 Hz, 4H).

**Synthesis of 4-(6-chloropyrimidin-4-yl)morpholine (7A)**: To a stirred solution of 4, 6-dichloropyrimidine (**7C**) 0.5 g (3.37 mmol, 1.0 eq.) in MeCN (10 mL) was added morpholine 0.32 g (3.71 mmol, 1.1 eq.), followed by the addition of DIPEA (5.19 g, 5.05 mmol, 1.5 eq.). and stirred at room temperature for 12 h. Reaction progress was monitored by TLC, after completion, the mixture was diluted with $H_2O$ (10 mL), solid formed was filtered and dried to give 4-(6-chloropyrimidin-4-yl) morpholine (**7A**) (0.28 g, 42%) as a white solid. m/z 200.03 $[M+H]^+$ .

$^1$H NMR (400 MHz, DMSO-d6) δ 8.37 (s, 1H), 6.98 (s, 1H), 3.40-3.90 (m, 8H).

**Synthesis of N-(4-(6-morpholinopyrimidin-4-yl)phenyl)cyclopropanecarboxamide (7)**: To a stirred solution of 4-(6-chloropyrimidin-4-yl)morpholine (**7A**) (0.25 g, 1.25 mmol, 1 eq.) in 1,4-

dioxane (5.0 mL) was added N-(4-(4,4,5,5tetramethyl-1,3,2-dioxaborolan-2-yl) phenyl)cyclopropanecarboxamide (**7B**) (0.43 g, 1.50 mmol, 1.2 eq.). To this mixture was added $K_2CO_3$ (0.69 g, 1.87 mmol, 1.5 eq.) in $H_2O$ (2.5 mL). The reaction mixture was degassed with nitrogen for 20 min and $(PPh_3)_2PdCl_2$ (0.051 g, 0.06 mmol, 0.05 eq.) was added. The reaction mixture was heated at 90 $^oC$ for 12 h. Reaction progress was monitored by TLC and LCMS, after completion, the mixture was diluted with $H_2O$ (10 mL) and extracted with EtOAc (3 x 20 mL). The organic layer was washed with brine solution, dried over $Na_2SO_4$, filtered and concentrated to give crude compound. The resulting crude was purified by prep. HPLC using 0.1% formic acid in $H_2O$/MeCN. The product fractions were collected and concentrated to give N-(4-(6-morpholinopyrimidin-4-yl)phenyl)cyclopropane carbox-amide, **7** (0.012 g, 9%) as a white solid. m/z $[M+H]^+$ 325.20.

$^1H$ NMR (400 MHz, DMSO-d6) δ 10.41 (s, 1H), 8.56 (s, 1H), 8.13 (d, J = 8.6 Hz, 2H), 7.71 (d, J = 8.6 Hz, 2H), 7.28 (s, 1H), 3.59- 3.79 (m, 8H), 1.85 – 1.77 (m, 1H), 0.86 – 0.80 (m, 4H).

$^{13}C$ NMR (176 MHz, DMSO) δ 172.37, 162.96, 161.73, 158.36, 141.71, 132.01, 127.91, 119.07, 97.95, 66.33, 44.39, 15.13, 7.85

**Synthesis of N-(4-(5-methyl-2-morpholinopyrimidin-4-yl)phenyl)cyclopropanecarboxamide (8)**:

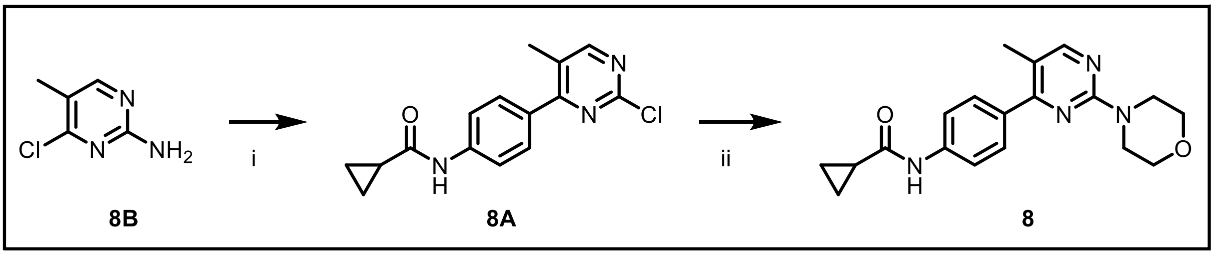


**Reaction scheme S4**: i. $PdCl_2(PPh_3)_2$, $K_2CO_3$, dioxane, water, 90 °C, 12 hr, 59%; ii. DIPEA, DMF, 80 °C, 5 hr, 17%.

**Synthesis of N-(4-(2-chloro-5-methylpyrimidin-4-yl)phenyl)cyclopropane carboxamide (8A)**: To a stirred solution of 2,4-dichloro-5-methylpyrimidine (**8B**) (0.25 g, 1.53 mmol, 1.0 eq.) in 1,4-dioxane (4.0 mL) was added N-(4-(4,4,5,5-tetramethyl1,3,2-dioxaborolan-2-yl)phenyl) cyclo-propanecarboxamide (**7B**) (0.52 g, 1.84 mmol, 1.1 eq.) and $K_2CO_3$ (0.31 g, 2.29 mmol, 1.5 eq.) in $H_2O$ (4.0 mL) was added. The reaction mixture was degassed with nitrogen for 20 min and $(PPh_3)_2PdCl_2$ (0.1 g, 0.016 mmol, 0.1 eq.) was added. The reaction mixture was then heated at 90 °C for 12 h. Reaction progress was monitored by TLC and LCMS. After completion, the mixture was diluted with $H_2O$ (10 mL) and extracted using EtOAc (3 x 20 mL). The organic layer was washed with brine solution, dried over Na2SO4, filtered and concentrated to give crude compound. The resulting crude was purified by flash column chromatography using 30 % EtOAc/n-hexane. The product fractions were collected and concentrated under reduced pressure to give N-(4-(2-chloro-5methylpyrimidin-4-yl)phenyl)cyclopropane-carboxamide (**8A**) (0.25 g, 59%) as an off white solid. m/z: 288.16 $[M+H]^+$

**Synthesis of N-(4-(5-methyl-2-morpholinopyrimidin-4-yl)phenyl)cyclopropane carboxamide (8)**: To a stirred solution of N-(4-(2-chloro-5-methylpyrimidin-4-yl)phenyl)cyclopropane-carboxamide (**8A**) (0.25 g, 0.87 mmol, 1.0 eq.) in DMF (5.0 mL) was added morpholine (0.12 g,

1.30 mmol, 1.5 eq.), followed by the addition of DIPEA (0.16 g, 1.30 mmol, 1.5 eq.). The reaction mixture was heated at 80 $^{o}$C for 5 h. Reaction progress was monitored by TLC, after completion, the mixture was diluted with $H_2O$ (20 mL) and extracted using EtOAc (3 x 30 mL). The organic layer was washed with brine solution, dried over $Na_2SO_4$, filtered and concentrated to give crude compound. The resulting crude was purified by HPLC using 0.1% formic acid in $H_2O$/MeCN. Product fractions were collected and lyophilized to give N-(4-(5-methyl-2morpholinopyrimidin-4-yl)phenyl) cyclopropane carboxamide (**8**) (0.05 g, 17%) as a white solid. m/z: 339.22 $[M+H]^+$

$^1$H NMR (400 MHz, DMSO-d6) δ 10.40 (s, 1H), 8.29 (s, 1H), 7.74 – 7.60 (m, 4H), 3.67 (s, 8H), 2.20 (s, 3H), 1.85 – 1.77 (m, 1H), 0.83 (d, J = 4.3 Hz, 4H).

$^{13}$C NMR (176 MHz, DMSO) δ 172.34, 164.06, 160.90, 160.25, 140.63, 133.36, 129.90, 118.76, 116.24, 66.47, 44.59, 16.51, 15.09, 7.80.

Synthesis of tert-butyl (3-(4-(4-(cyclopropanecarboxamido)phenyl)-2-morpholinopyrimidine-5-carboxamido)propyl)carbamate (9):

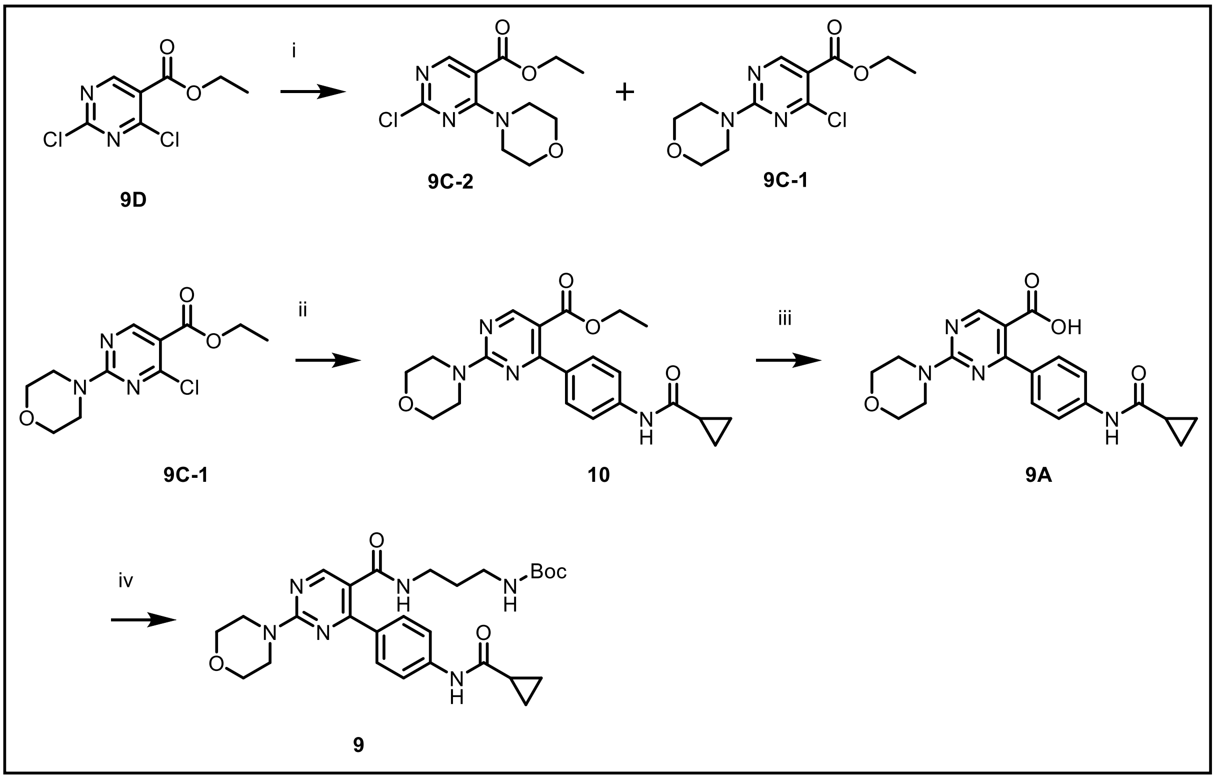


**Reaction scheme S5**: i. morpholine, DIPEA, NMP, RT, 3 hr, 12-19%; ii. $PdCl_2(PPh_3)_2$, $K_2CO_3$, dioxane, water, 90 °C, 12 hr, 8%; iii. NaOH, THF, MeOH, Water, RT, 12 hr, 45%; iv. EDC.HCl, HOBt, DIPEA, DMF, RT, 12 hr, 23%.

**Synthesis of ethyl 2-chloro-4-morpholinopyrimidine-5-carboxylate (9C-2)**:

To a stirred solution of ethyl 2,4-dichloropyrimidine-5-carboxylate (Int-1) (0.50 g, 2.26 mmol, 1.0 eq.) in NMP (5.0 mL) was added morpholine (0.18 g, 2.26 mmol, 1.2 eq.). followed by DIPEA (0.43 g, 3.39 mmol, 1.5 eq.). Reaction mixture was stirred at room temperature for 3 h. Reaction progress was monitored by TLC, after completion, the mixture was diluted with $H_2O$ (10 mL) and extracted using EtOAc (2 x 30 mL). The organic layer was washed with brine solution, dried over $Na_2SO_4$, filtered and concentrated to give crude compound. The resulting crude was purified by flash column chromatography eluting with 15 % EtOAc/n-hexane. Two fractions were

isolated, characterised by $^1H$ NMR and compared with reported value. Ethyl 2-chloro-4-
morpholinopyrimidine-5-carboxylate, **9C-2** (0.07 g, yield 11%) was isolated as a white solid. m/z: 272.11 $[M+H]^+$

$^1H$ NMR (400 MHz, Chloroform-d) δ 8.57 (s, 1H), 4.36 (q, J = 7.1 Hz, 2H), 3.83 – 3.75 (m, 4H), 3.68 – 3.61 (m, 4H), 1.39 (t, J = 7.1 Hz, 3H).

**Ethyl 4-chloro-2-morpholinopyrimidine-5-carboxylate, 9C-1** (0.12 g yield 19%) was isolated as a white solid. m/z: 272.08 $[M+H]^+$

$^1H$ NMR (400 MHz, Chloroform-d) δ 8.84 (s, 1H), 4.37 (q, J = 7.0 Hz, 2H), 3.96 – 3.89 (m, 4H), 3.78 (d, J = 4.7 Hz, 4H), 1.39 (t, J = 7.1 Hz, 3H).

**Synthesis of ethyl 4-(4-(cyclopropanecarboxamido)phenyl)-2morpholino-pyrimidine-5-carboxylate** (**10**): To a stirred solution of ethyl 4-chloro-2-morpholinopyrimidine-5carboxylate, **9C-1** (0.25 g, 0.92 mmol, 1.0 eq.) in 1,4-dioxane (5.0 mL) was added N-(4-(4,4,5,5-tetramethyl-1,3,2-dioxaborolan-2-yl)phenyl)cyclopropanecarboxamide (**7B**) (0.29 g, 1.01 mmol, 1.1 eq.) and $K_2CO_3$ (0.19 g, 1.38 mmol, 1.5 eq.) in $H_2O$ (5.0 mL) was then added. The reaction mixture was degassed with nitrogen for 20 min and $(PPh_3)_2PdCl_2$ (0.003 g, 0.046 mmol, 0.05 eq.) was added to it. The reaction mixture was stirred at 90 °C for 12 h. Reaction progress was monitored by TLC and LCMS, after completion, the mixture was diluted with $H_2O$ (10 mL) and extracted using EtOAc (3 x 20 mL). The organic layer was washed with brine solution, dried over Na2SO4, filtered and concentrated to give crude compound. The resulting crude was purified by flash column chromatography eluting the product with 30 % EtOAc/n-hexane. The pure fraction was collected and concentrated under reduced pressure to give ethyl 4-(4-(cyclopropanecarboxamido) phenyl)-2-morpholinopyrimidine-5-carboxylate (**10**) (0.03 g, 8%) as an off white solid. m/z: 397.23 $[M+H]^+$

$^1H$ NMR (400 MHz, DMSO-d6) δ 10.40 (s, 1H), 8.74 (s, 1H), 7.66 (d, J = 8.3 Hz, 2H), 7.50 (d, J = 8.4 Hz, 2H), 4.11 (q, J = 6.7 Hz, 2H), 3.85 (s, 4H), 3.68 (s, 4H), 1.81 (d, J = 6.2 Hz, 1H), 1.08 (t, J = 6.9 Hz, 3H), 0.83 (s, 4H).

$^{13}C$ NMR (176 MHz, DMSO) δ 172.36, 166.71, 166.34, 161.12, 141.33, 132.96, 129.99, 118.39, 112.41, 66.39, 60.84, 44.37, 15.11, 14.24, 7.85.

**Synthesis of 4-(4-(cyclopropanecarboxamido)phenyl)-2morpholinopyrimidine-5-carboxylic acid (9A)**: To a stirred solution of ethyl 4-(4-(cyclopropanecarboxamido)phenyl)-2morpholino pyrimidine-5-carboxylate (**9B**) (0.10 g, 0.25 mmol, 1.0 eq.) in THF (2.0 mL) and EtOH (1.0 mL) was added NaOH (0.015 g, 0.38 mmol, 1.5 eq) in $H_2O$ (1.0 mL). The reaction mixture was stirred at room temperature for 12 h. Reaction progress was monitored by TLC and LCMS, after completion, the mixture was diluted with $H_2O$ (10 mL) and extracted using EtOAc (1 x 10 mL). The aqueous layer was acidified using dilute hydrochloride, solid formed was filtered and dried to give 4-(4-(cyclopropane-carboxamido)phenyl)-2morpholinopyrimidine-5-carboxylic acid (**9A**) (0.042 g. 45%) as a white solid. m/z: 369.19 $[M+H]^+$

$^1H$ NMR (400 MHz, DMSO-d6) δ 10.49 (s, 1H), 8.75 (s, 1H), 7.66 (d, J = 8.5 Hz, 2H), 7.52 (d, J = 8.4 Hz, 2H), 3.84 (s, 4H), 3.67 (s, 4H), 1.89 – 1.81 (m, 1H), 0.82 (s, 4H).

13C NMR (176 MHz, DMSO) δ 172.35, 167.56, 166.85, 161.24, 161.09, 141.21, 133.21, 130.07, 118.35, 113.05, 66.41, 44.35, 15.09, 7.82.

**Synthesis of tert-butyl (3-(4-(4-(cyclopropane carboxamide) phenyl)2-morpholinopyrimidine-5-carboxamido) propyl) carbamate (9)**: To a stirred solution of 4-(4-(cyclopropanecarboxamido)phenyl)-2morpholinopyrimidine-5-carboxylic acid (**9A**) (0.15 g, 0.40 mmol, 1.0 eq.) in DMF (3.0 mL) was added EDC.HCl (0.15 g, 0.61 mmol, 1.5 eq), HOBt (0.027 g, 0.2 mmol, 0.5 eq) and DIPEA (0.08 g, 0.60 mmol, 1.5 eq) at 0 $^{o}$C. After stirring for 15 mins at the same temperature tert-butyl (3-aminopropyl)carbamate (0.08 g, 0.44 mmol, 1.1 eq) was added to above mixture and the reaction was allowed to warm to room temperature and stir for an additional 12 h. Reaction progress was monitored by TLC and LCMS, after completion, the mixture was diluted with $H_2O$ (10 mL) and extracted using EtOAc (3 x 10 mL). The organic layer was washed with brine solution, dried over $Na_2SO_4$, filtered and concentrated to give crude compound. The resulting crude was purified by flash column chromatography eluting the product with 40 % EtOAc/n-hexane. The desired fractions were collected and concentrated to give tert-butyl (3-(4-(4-(cyclopropanecarboxamide) phenyl)2-morpholinopyrimidine-5-carboxamido) propyl) carbamate (**9**) (0.05 g) as a white solid. (yield-23%). m/z: 525.4 $[M+H]^+$

$^1$H NMR (400 MHz, DMSO-d6) δ 10.39 (s, 1H), 8.37 (s, 1H), 8.23 (s, 1H), 7.64 (s, 4H), 6.77 (s, 1H), 3.80 (s, 4H), 3.34 (s, 4H), 3.15 – 3.08 (m, 2H), 2.89 (s, 2H), 1.80 (s, 1H), 1.52 (s, 2H), 1.37 (s, 9H), 0.82 (s, 4H).

$^{13}$C NMR (176 MHz, DMSO) δ 172.35, 167.42, 141.44, 129.71, 119.18, 118.73, 77.95, 66.47, 44.40, 40.49, 40.35, 40.23, 40.11, 40.00, 39.88, 39.76, 39.64, 38.03, 37.28, 29.68, 28.72, 15.08, 7.85.

**Synthesis of N-(4-((2-fluorophenyl)(hydroxy)methyl)phenyl)acetamide** (11):

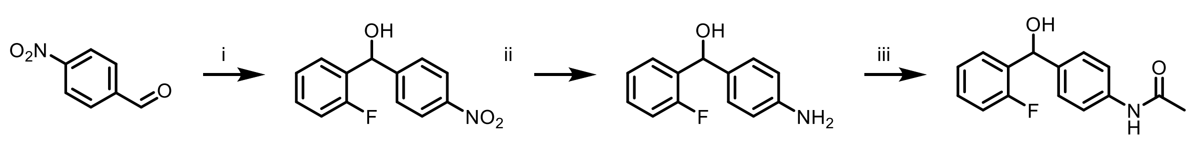


**Reaction scheme S6**: i. (2-fluorophenyl)boronic acid, tri-1-naphthyl phosphine, $K_2CO_3$, $Pd(OAc)_2$, [bmim][$PF_6$], water, 65 °C, 16 h, 42%; ii. Fe, $NH_4Cl$, EtOH, water, 70 °C, 4 hr, 56%; iii. Acetyl chloride, $Na_2CO_3$, dioxane, water, RT, 3 hr, 33.5%.

**Synthesis of (4-aminophenyl)(2-fluorophenyl)methanol (11B)**: A mixture of 4-nitrobenzaldehyde (Int-1) (7.0 g, 19 mmol, 1 eq.) in THF (70 mL) and added (2-fluorophenyl)boronic acid (6.5 g, 26 mmol, 1.2 eq.) and K2CO3 (8.22 g, 59 mmol, 3 eq.). The reaction was degassed with nitrogen for 10 min, added Tri-1-naphthylphosphine (2.4 g, 10 mmol, 0.05 eq.) and palladium(II) chloride (1.16 g, 19 mmol, 0.05 eq.). It was stirred at 65 °C for 3 h. The reaction was monitored on TLC. After completion of the reaction, it was quenched with H2O (200 mL) and extracted with EtOAc (2 X 100 mL), dried over sodium sulfate and concentrated under reduced pressure to give the crude compound which was purified by flash column chromatography, eluting the desired product with 20% EtOAc: n-hexane to give (2-fluorophenyl)(4nitrophenyl) EtOH (**11B**) (7.0 g, 61.10%) as a cream solid. m/z: 246.0 $[M+H]^+$.

$^1H$ NMR (400 MHz, DMSO-$d_6$) δ 8.20 (d, J = 8.6 Hz, 2H), 7.63 (d, J = 8.5 Hz, 2H), 7.55 (t, J = 7.0 Hz, 1H), 7.36 – 7.28 (m, 1H), 7.27 – 7.11 (m, 2H), 6.38 (d, J = 4.3 Hz, 1 OH), 6.08 (d, J = 4.0 Hz, 1H).

**Synthesis of (4-aminophenyl)(2-fluorophenyl)methanol (11A)**: To a mixture of (2-fluorophenyl)(4-nitrophenyl) EtOH (Int-3) (0.14 g, 0.64 mmol, 1 eq.) in EtOH (1 mL) and H2O (1 mL), were added Fe powder (0.361 g, 6.4 mmol, 10.0 eq) and ammonium chloride (0.341 g, 6.4 mmol, 10.0 eq) and heated to reflux for 4 h. After completion of the reaction, the reaction was filtered through celite-pad and concentrated under reduced pressure to give the desired product as (4-aminophenyl)(2-fluorophenyl)methanol (**11A**) (0.07 g, 56.9 %) as a solid. m/z: 218.0 $[M+H]^+$

$^1$H NMR (400 MHz, DMSO-$d_6$) δ 7.58 (t, J = 7.4 Hz, 1H), 7.21 (dt, J = 24.8, 7.2 Hz, 2H), 7.12 – 7.02 (m, 1H), 6.95 (d, J = 8.0 Hz, 2H), 6.47 (d, J = 8.1 Hz, 2H), 5.76 (d, J = 3.5 Hz, 1H), 5.65 (d, J = 4.1 Hz, 1H), 4.97 (s, 2H).

**Synthesis of N-(4-((2-fluorophenyl)(hydroxy)methyl)phenyl)acetamide (11)**: To a mixture of (4-aminophenyl)(2-fluorophenyl) EtOH (Int-4) (0.1 g, 0.53 mmol, 1 eq.) in 1,4-dioxane (2 mL) and H2O (2 mL), was added sodium bicarbonate (0.085 g, 0.8 mmol, 1.5 eq.), cooled to 0 $^o$C and acetyl chloride (0.05 g, 0.64 mmol, 1.2 eq.) was added to stir for 3 h at room temperature. After completion of the reaction, the reaction mixture quenched with H2O (20 mL) and extracted with EtOAc (2 X 20 mL), dried over sodium sulfate and concentrated under reduced pressure to give the crude compound which was purified with RP-HPLC purification using (i) 0.1% formic acid in H2O (ii) MeCN to give to N-(4-((2-fluorophenyl)(hydroxy)methyl)phenyl)acetamide (**11**) (0.04 g, 33.51 %) as a white solid. m/z: 260.0 $[M+H]^+$.

$^1$H NMR (400 MHz, DMSO-$d_6$) δ 9.90 (s, 1H), 7.57 (t, J = 1.6 Hz, 1H), 7.50 (t, J = 8.8 Hz, 2H), 7.25-7.20 (m, 5H), 5.94-5.89 (m, 2H), 2.02 (s, 3H).

$^{13}$C NMR (176 MHz, DMSO) δ 168.62, 160.14, 139.37, 138.64, 133.05, 129.16, 128.16, 127.05, 124.80, 119.26, 115.52, 68.08, 24.41.

**Synthesis of N-((4-acetamidophenyl)(2-fluorophenyl)methyl)-2-chlorobenzamide (12)**:

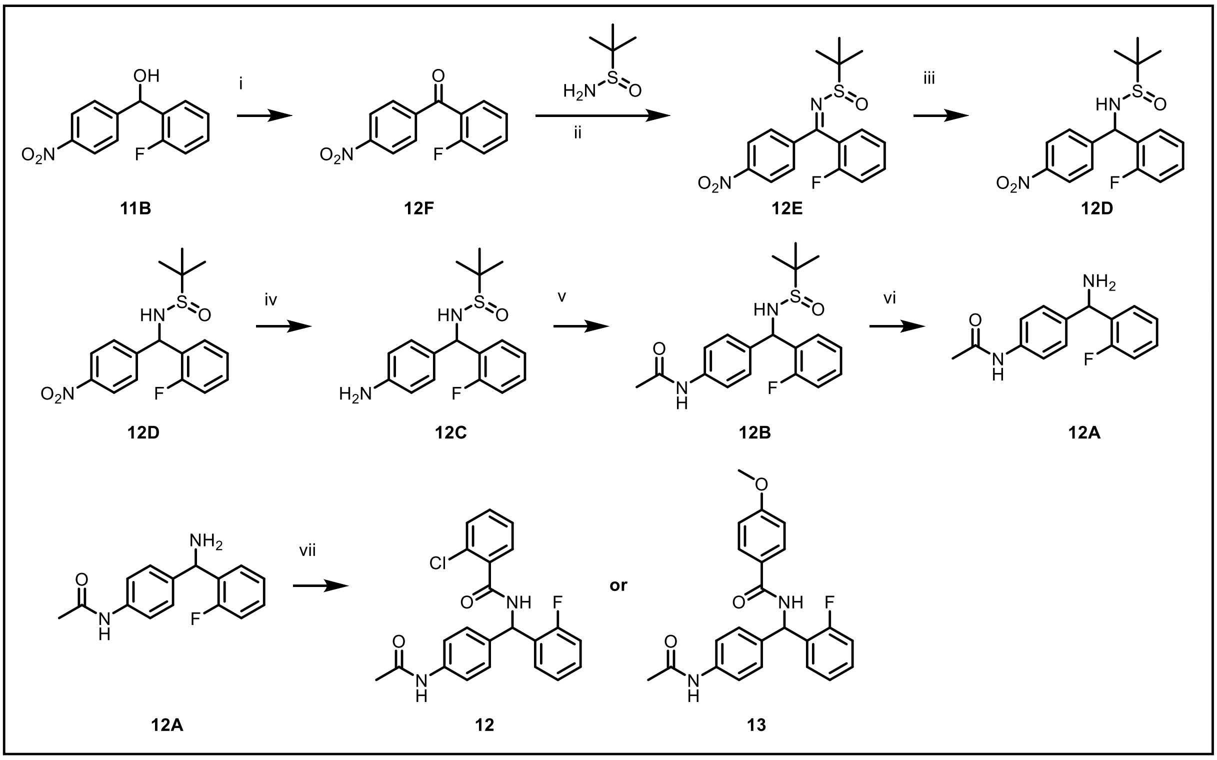


**Reaction scheme S7**: i. PCC, DCM, RT, 12 hr, 50%; ii. Titanium ethoxide, dioxane, 100 °C, 3 hr, 75%; iii. $NaBH_4$, MeOH, 0 °C, 30 min, 78%; iv. Pd/C, $H_2$, MeOH, RT, 5 hr, 81%; v. acetic anhydride, AcOH, RT, 2 hr, 55%; vi. HCl (4 M in dioxane), RT, 2 hr, 19%; vii. 2-chlorobenzoic acid or 4-methoxybenzoic acid, HATU, DIPEA, THF, RT, 2 hr, 7-35%.

**Synthesis of (4-aminophenyl)(2-fluorophenyl)methanol (11B)**: A mixture of 4-nitrobenzaldehyde (**11C**) (7.0 g, 19 mmol, 1 eq.) in THF (70 mL) and added (2-fluorophenyl)boronic acid (6.5 g, 26 mmol, 1.2 eq.) and K2CO3 (8.22 g, 59 mmol, 3 eq.). The reaction was degassed with nitrogen for 10 min, added Tri-1-naphthylphosphine (2.4 g, 10 mmol, 0.05 eq.) and palladium(II) chloride (1.16 g, 19 mmol, 0.05 eq.). It was stirred at 65 °C for 3 h. The reaction was monitored on TLC. After completion of the reaction, it was quenched with $H_2O$ (200 mL) and extracted with EtOAc (2 X 100 mL), dried over sodium sulfate and concentrated under reduced pressure to give the crude compound which was purified by flash column chromatography, eluting the desired product with 20% EtOAc: n-hexane to give (4-

aminophenyl)(2-fluorophenyl)methanol (**11B**) (7.0 g,  61.10%) as a cream solid. m/z: 246.0 [M-H]$^-$,

$^1$H NMR (400 MHz, DMSO-$d_6$) δ 8.20 (d, J = 8.6 Hz, 2H), 7.63 (d, J = 8.5 Hz, 2H), 7.55 (t, J = 7.0 Hz, 1H), 7.36 – 7.28 (m, 1H), 7.27 – 7.11 (m, 2H), 6.38 (d, J = 4.3 Hz, 1 OH), 6.08 (d, J = 4.0 Hz, 1H).

**Synthesis of (2-fluorophenyl)(4-nitrophenyl)methanone (12F)**: To a mixture of (4-aminophenyl)(2-fluorophenyl)methanol (**11B**) (7.0 g, 27 mmol, 1 eq.) in dichloromethane (20 mL) at 0 °C, was added  pyridinium chlorochromate (8.7 g, 40 mmol, 1.5 eq.) and stirred for 1h.  Reaction was monitored on TLC. After completion of the reaction, the reaction mixture was quenched with H2O (200 mL) and extracted with EtOAc (2 X 100 mL), dried over sodium sulfate and concentrated under reduced pressure to give the crude compound **12F** which was forwarded for the next step without purification. m/z: 246.22 [M+1].

**Synthesis of (Z)-N-((2-fluorophenyl)(4-nitrophenyl)methylene)-2-methyl propane-2-sulfinamide (12E)**: To a mixture of (2-fluorophenyl)(4-nitrophenyl)methanone (**12F**) (6.0 g, 24 mmol, 1 eq.) in 1,4-dioxane (60 mL), was added titanium(IV) ethoxide (16.75 g, 73 mmol, 3 eq.) and 2-methylpropane-2-sulfinamide (3.58 g, 29 mmol, 1.2 eq.) and stirred 100° C for 4 h.  The reaction was monitored on TLC. After completion of the reaction, the reaction mixture was quenched with H2O (400 mL) and extracted with EtOAc (2 X 100 mL), dried over sodium sulfate and concentrated under reduced pressure to give the crude compound which was purified by flash column chromatography, eluting the desired product with 30% EtOAc: n-hexane to give (Z)-N-((2fluorophenyl)(4-nitrophenyl)methylene)-2-methyl propane-2-sulfinamide (**12E**)  (5.9 g, 69.22%) as a solid. m/z: 349.1 [M+H]$^+$.

$^1$H NMR (400 MHz, DMSO-$d_6$) δ 8.35 (d, J = 8.6 Hz, 2H), 7.85 (d, J = 8.3 Hz, 2H), 7.63 (d, J = 6.2 Hz, 1H), 7.41 (dd, J = 18.8, 10.0 Hz, 3H), 1.27 (s, 9H).

**Synthesis of N-((2-fluorophenyl)(4-nitrophenyl)methyl)-2methylpropane-2-sulfinamide (12D)**: To a mixture of (Z)-N-((2-fluorophenyl)(4-nitrophenyl)methylene)-2methylpropane-2-sulfinamide (**12E**) (5.9 g, 16 mmol, 1 eq.) in EtOH (60 mL), cooled at 0°C added sodium borohydride (0.576 g, 16 mmol, 1 eq.) then stirred reaction mass at 1h. Reaction was monitoring on TLC. After completion of the reaction, the reaction mixture quenched with H2O (200 mL) and extracted with EtOAc (2 X 100 mL), dried over sodium sulfate and concentrated under reduced pressure to give the crude compound which was purified by flash column chromatography, eluting the desired product with 40% EtOAc: n-hexane to give N-((2-fluorophenyl)(4-nitrophenyl)methyl)2-methylpropane-2-sulfinamide (**12D**) (5.8 g, 97.72%) as a cream solid. m/z: 351 $[M+H]^+$

**Synthesis of N-((4-aminophenyl)(2-fluorophenyl)methyl)-2methylpropane-2-sulfinamide (12C)**: To a mixture of N-((2-fluorophenyl)(4-nitrophenyl)methyl)-2-methylpropane-2-sulfinamide (**12D**) (5.8 g, 17 mmol, 1 eq.) in EtOH (120 mL), degassed with nitrogen, added Pd/C (0.7% with 50% moisture) ( 0.7 g) and stirred under hydrogen balloon pressure for 1 h. The reaction mixture was filtered through celite-pad, washed with EtOH, the filtrate was concentrated under reduced pressure to give N-((4-aminophenyl)(2fluorophenyl)methyl)-2-methylpropane-2-sulfinamide (**12C**) (5.3 g, 98.22%) as a solid. m/z: 320.49 $[M+H]^+$
$^1$H NMR (400 MHz, DMSO-$d_6$) δ 7.76 (t, J = 7.7 Hz, 1H), 7.45 (d, J = 8.3 Hz, 2H), 7.22 (dd, J = 20.4, 7.8 Hz, 2H), 7.06 – 6.96 (m, 1H), 6.50-6.46 (m, 2H), 5.73 (s, 1H), 5.08 (m, 2H, $NH_2$), 1.14 – 1.05 (m, 9H). (Mixture of atropisomers)

**Synthesis of N-(4-(((tert-butylsulfinyl)amino)(2-fluorophenyl)methyl)phenyl)acetamide (12B)**: To a mixture of N-((4-aminophenyl)(2-fluorophenyl)methyl)-2methylpropane-2-sulfinamide (**12C**) (5.3 g, 16 mmol 1 eq.) in AcOH (10 mL) at 0 °C, was added acetic anhydride (2.56 g, 25 mmol, 1.5 eq.) and stirred for 1h. The reaction was monitored on LC-MS. After completion of the reaction, the reaction mixture was quenched with H2O (200 mL) and extracted with EtOAc (2 X 100 mL), dried over sodium sulfate and concentrated under reduced pressure to give the crude compound which was purified by flash column chromatography, eluting the desired product with 40% EtOAc: n-hexane to give N-(4-(((tert-butylsulfinyl)amino)(2-fluorophenyl)methyl)phenyl)acetamide (**12B**) (5.9 g, 98.39%) as a solid. m/z: 363.0 $[M+H]^+$

$^1$H NMR (400 MHz, DMSO-$d_6$) δ 9.94 (d, J = 3.6 Hz, 1H), 7.66 – 7.58 (m, 3H), 7.54 – 7.42 (m, 2H), 7.22 (ddt, J = 32.5, 26.0, 9.1 Hz, 4H), 6.04 (dd, J = 21.4, 6.4 Hz, 1H), 5.79 – 5.66 (m, 1H), 2.00 (d, J = 8.0 Hz, 3H), 1.13 (s, 9H). (Mixture of atropisomers).

**Synthesis of N-(4-(amino(2-fluorophenyl)methyl)phenyl)acetamide (12A)**: To a solution of N-(4-(((tert-butylsulfinyl)amino)(2-fluorophenyl)methyl)phenyl) acetamide (**12B**) (5.9 g, 16 mmol, 1 eq.) in 1,4-dioxane (30 mL), was added HCl in 1,4-dioxane (4.0 M, 30 mL) and stirred for 1h. The reaction mixture was concentrated to give crude which was purified with RP-HPLC using 0.1% formic acid in $H_2O$/ MeCN. Product fractions were collected and concentrated to give N-(4-(amino(2fluorophenyl)methyl)phenyl) acetamide (**12A**) (3.01 g, 73.39%) as a cream solid. m/z: 242.1 [M-16]

$^1$H NMR (400 MHz, DMSO-$d_6$) δ 10.00 (s, 1H), 7.66 (t, J = 7.2 Hz, 1H), 7.53 (d, J = 8.4 Hz, 2H), 7.31 (p, J = 9.9, 8.5 Hz, 4H), 7.27 – 7.12 (m, 1H), 5.49 (s, 1H), 2.02 (s, 3H).

$^{13}$C NMR (176 MHz, DMSO) δ 168.73, 163.70, 160.41, 159.02, 139.03, 129.68, 128.52, 127.80, 125.05, 119.43, 115.89, 51.86, 24.40.

**Synthesis of N-((4-acetamidophenyl)(2-fluorophenyl)methyl)-2chloro benzamide (12)**: To a mixture of N-(4-(amino(2-fluorophenyl)methyl)phenyl)acetamide (Int-9) (0.13 g, 0.5 mmol, 1 eq.) in THF (2 mL), was added 2-chlorobenzoic acid (0.094 g, 0.6 mmol, 1.2 eq.), stirred for 10 min, added HATU (0.3 g, 0.75 mmol, 1.5 eq.) and DIPEA (0.2 g, 1.5 mmol, 3 eq.) and stirred for 3 h.

After completion of the reaction, the reaction mixture was quenched with $H_2O$ (20 mL) and extracted with EtOAc (2 X 10 mL), dried over $Na_2SO_4$ and concentrated under reduced pressure to give the crude compound which was purified by flash column chromatography, eluting the desired product with 80% EtOAc/n-hexane and purified further with HPLC purification using 0.1% TFA in $H_2O$/MeCN. Product fractions were collected and dried to give to N-((4-acetamidophenyl)(2-fluorophenyl)methyl)-2-chloro benzamide (12) as a solid (0.06 g, 30.0%). m/z: 397.3 $[M+H]^+$

$^1H$ NMR (400 MHz, DMSO-$d_6$) δ 9.96 (s, 1H), 9.45 (d, J = 8.3 Hz, 1H), 7.55-7.30 (m, 8H), 7.25-7.19 (m, 4H), 6.52 (d, J = 8.4 Hz, 1H), 2.02 (s, 2H).

$^{13}C$ NMR (176 MHz, DMSO) δ 168.70, 166.10, 160.72, 159.33, 138.90, 137.17, 135.57, 131.23, 130.39, 129.95, 129.68, 129.56, 129.48, 129.38, 129.10, 128.08, 127.51, 124.96, 124.94, 119.48, 115.81, 115.69, 50.08, 50.06, 24.40.

**Synthesis of N-((4-acetamidophenyl)(2-fluorophenyl)methyl)-4methoxy benzamide (13)**: To a mixture of N-(4-(amino(2-fluorophenyl)methyl)phenyl)acetamide (**12A**) (0.15 g, 0.4 mmol, 1 eq.) in THF (2 mL), was added 4methoxybenzoic acid (0.087 g, 0.4 mmol, 1.2 eq.), stirred for 10 min, added HATU (0.178 g, 0.75 mmol, 1.5 eq.) and DIPEA (0.26 g, 2.2 mmol, 3 eq.) and stirred for 3 h. After completion of the reaction, the reaction mixture was quenched with $H_2O$ (20 mL) and extracted with EtOAc (2 X 10 mL), dried over $Na_2SO_4$ and concentrated under reduced pressure to give the crude compound which was purified by flash column chromatography, eluting the desired product with 80% EtOAc: n-hexane and further with RP-HPLC purification using 0.1% formic acid in $H_2O$/MeCN to give to N-((4acetamidophenyl)(2-fluorophenyl)methyl)-4-methoxybenzamide (**13**) (0.016 g,  7.04%) as a solid. m/z: 393.43$[M+H]^+$

$^1$H NMR (400 MHz, DMSO-$d_6$) δ 9.96 (s, 1H), 9.08 (d, J = 8.3 Hz, 1H), 7.92 (d, J = 8.7 Hz, 2H), 7.50 (dd, J = 21.2, 7.8 Hz, 3H), 7.35 (d, J = 6.9 Hz, 1H), 7.20 (q, J = 8.1 Hz, 4H), 7.00 (d, J = 8.7 Hz, 2H), 6.59 (d, J = 8.4 Hz, 1H), 3.81 (s, 3H), 2.02 (s, 2H).

$^{13}$C NMR (176 MHz, DMSO) δ 168.69, 165.70, 162.19, 138.85, 135.96, 130.00, 129.63, 129.36, 128.27, 126.76, 124.92, 119.48, 115.63, 113.89, 55.85, 24.41.

### 1H and 13C NMR of Compound 2

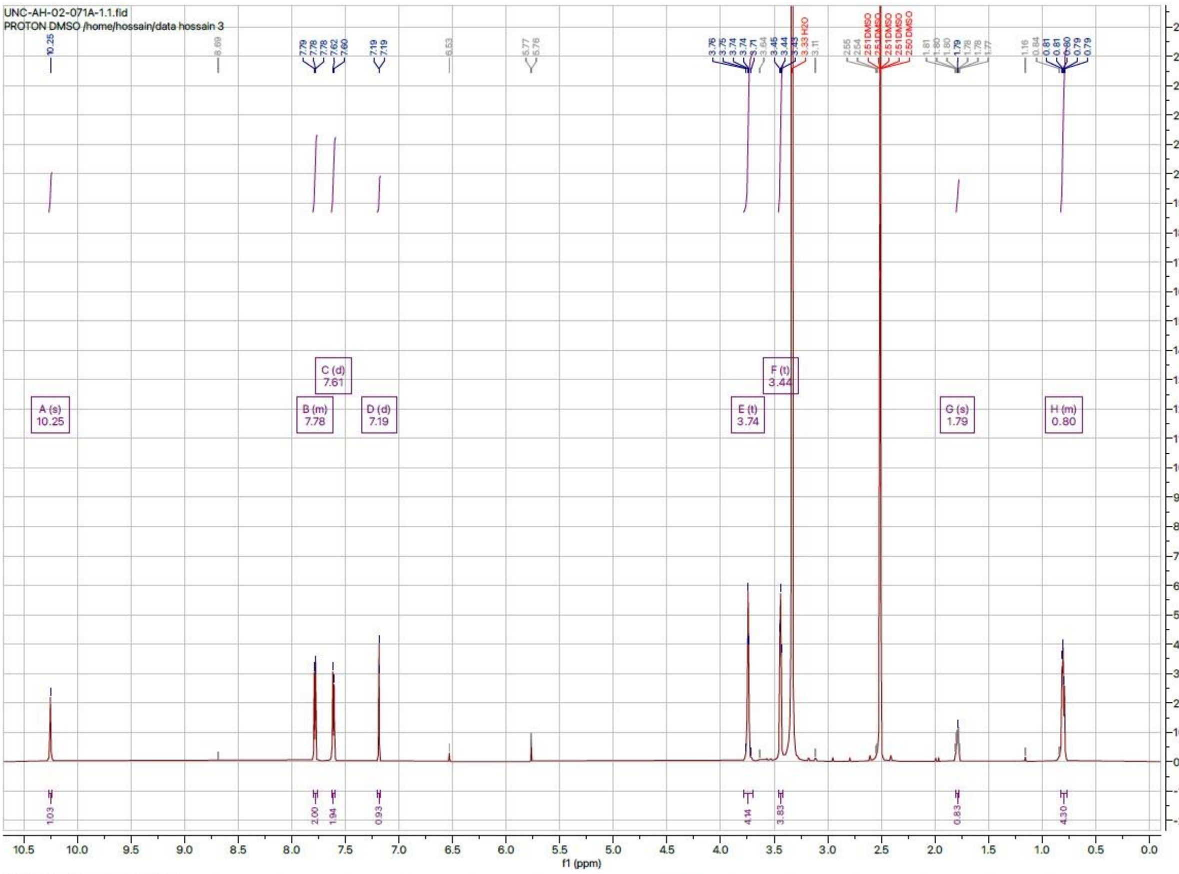


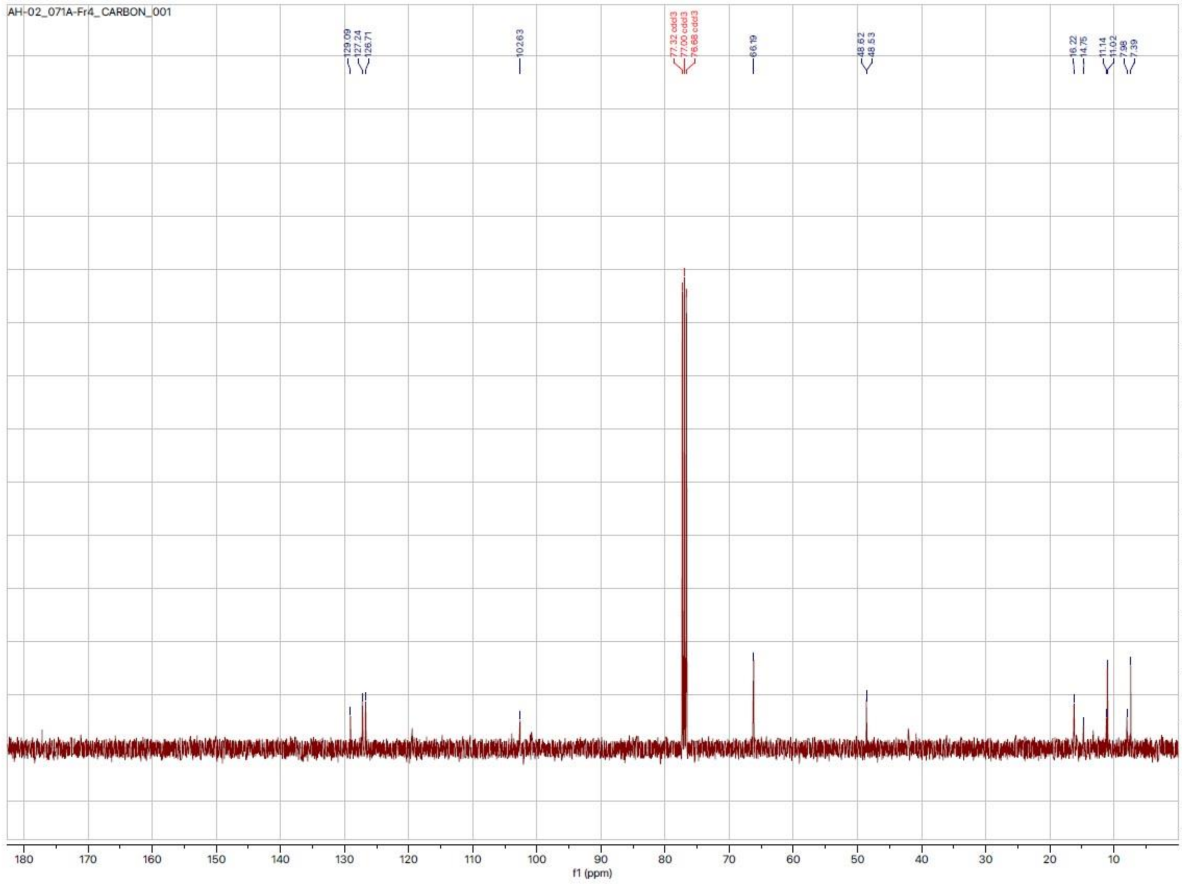

### 1H and 13C NMR of Compound 3

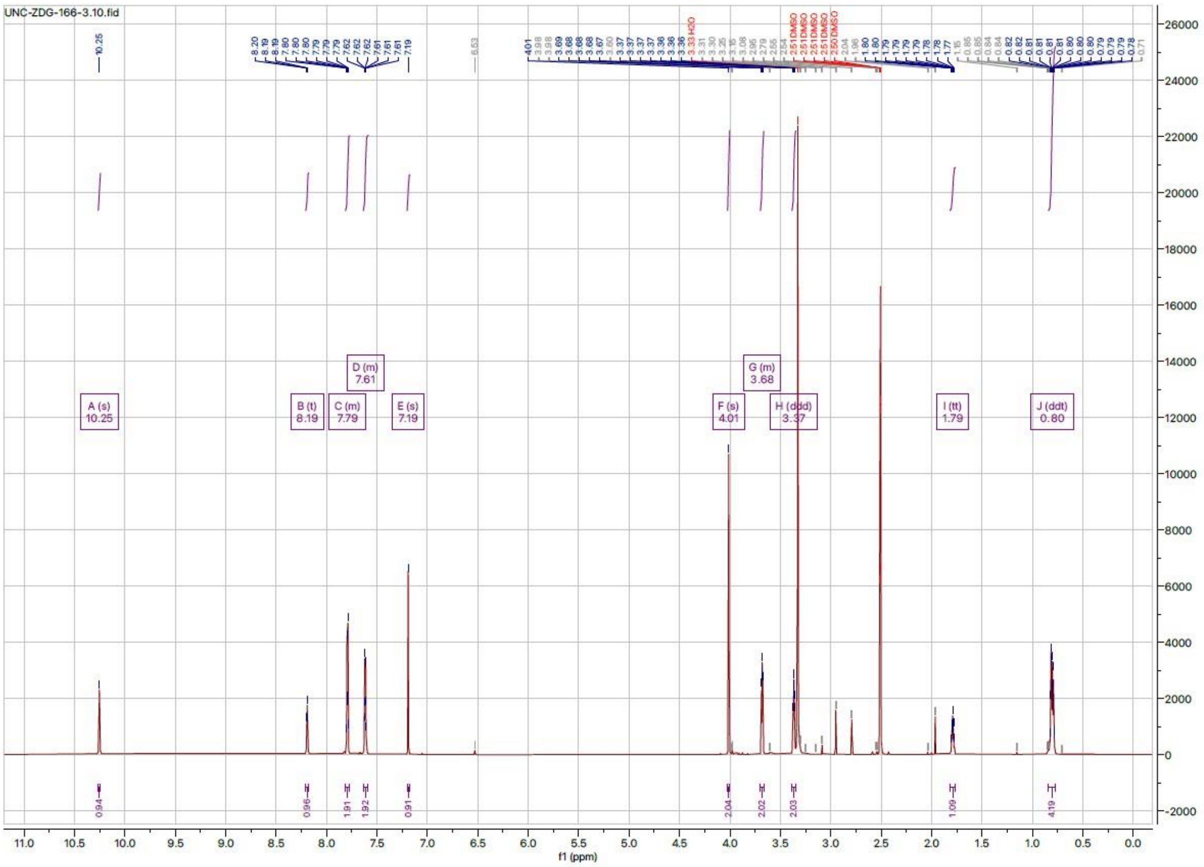


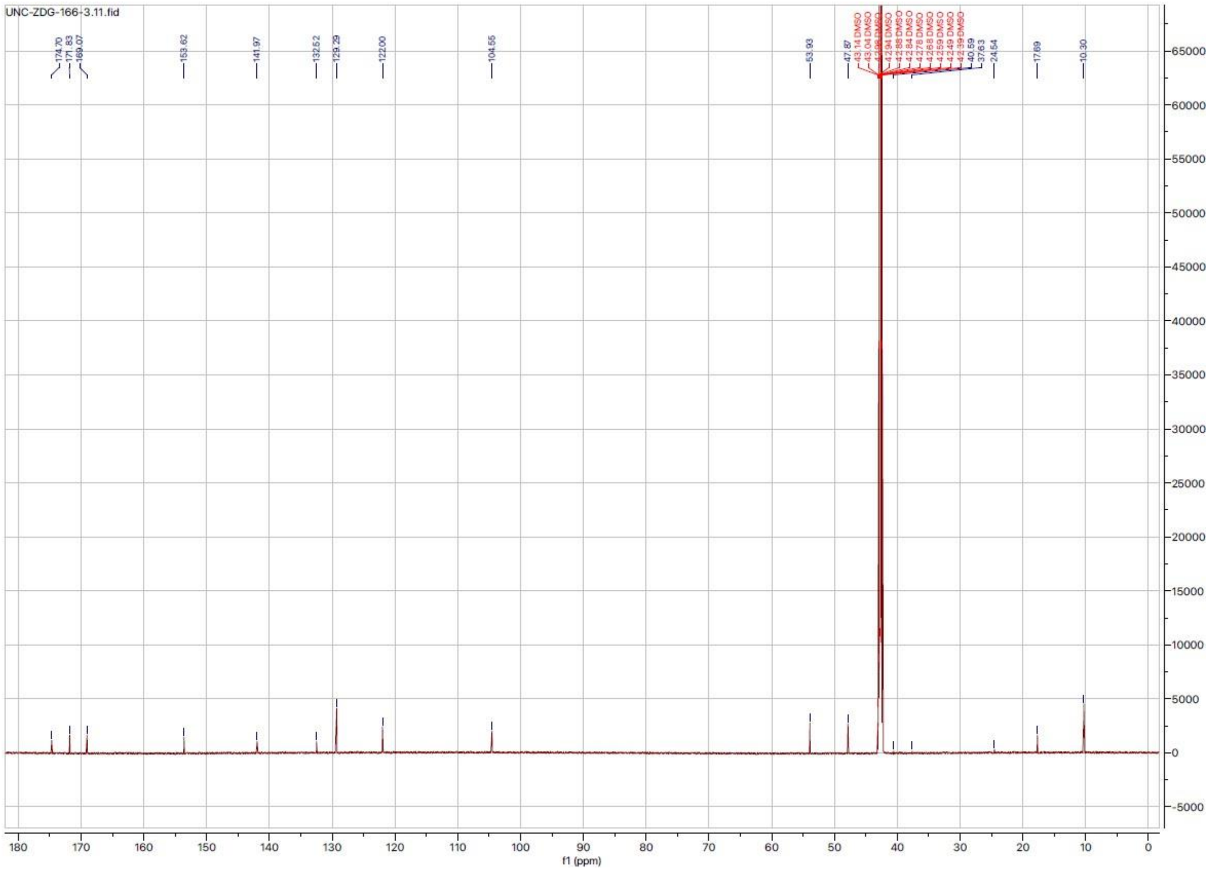

### 1H and 13C NMR of Compound 4

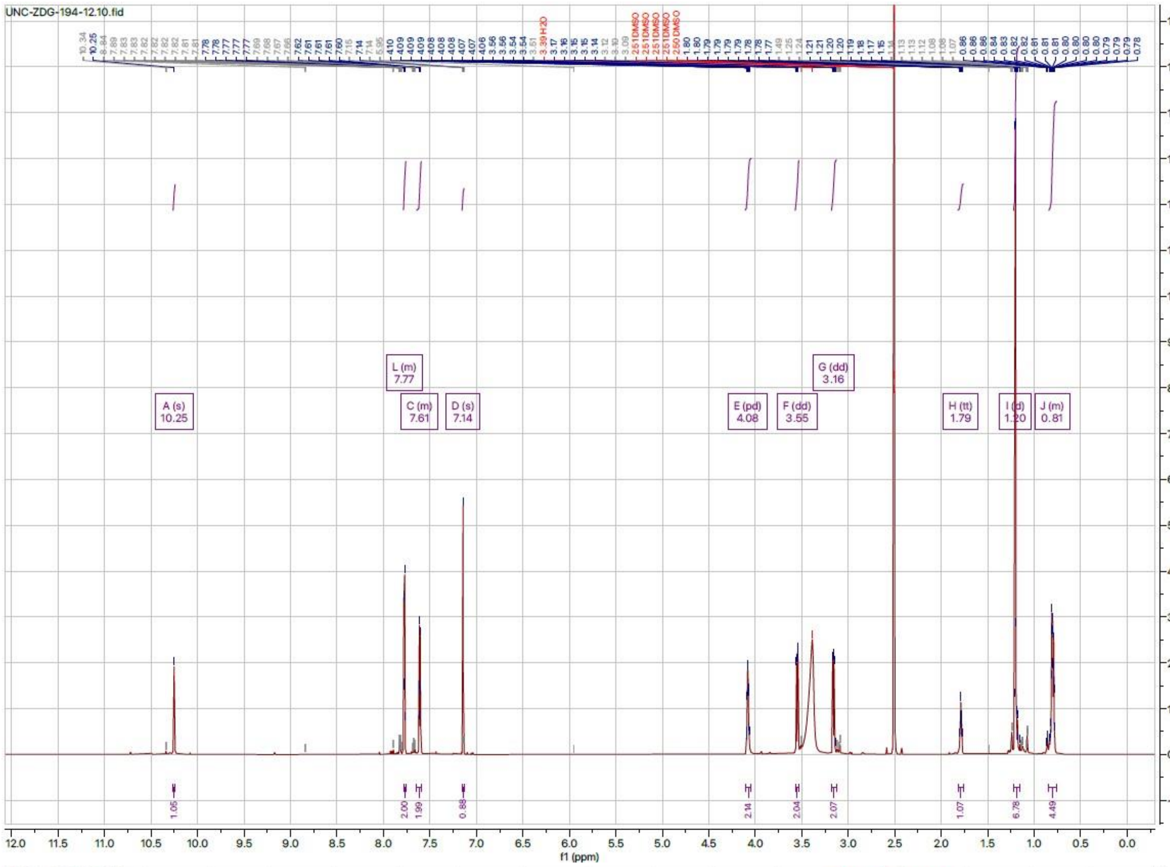

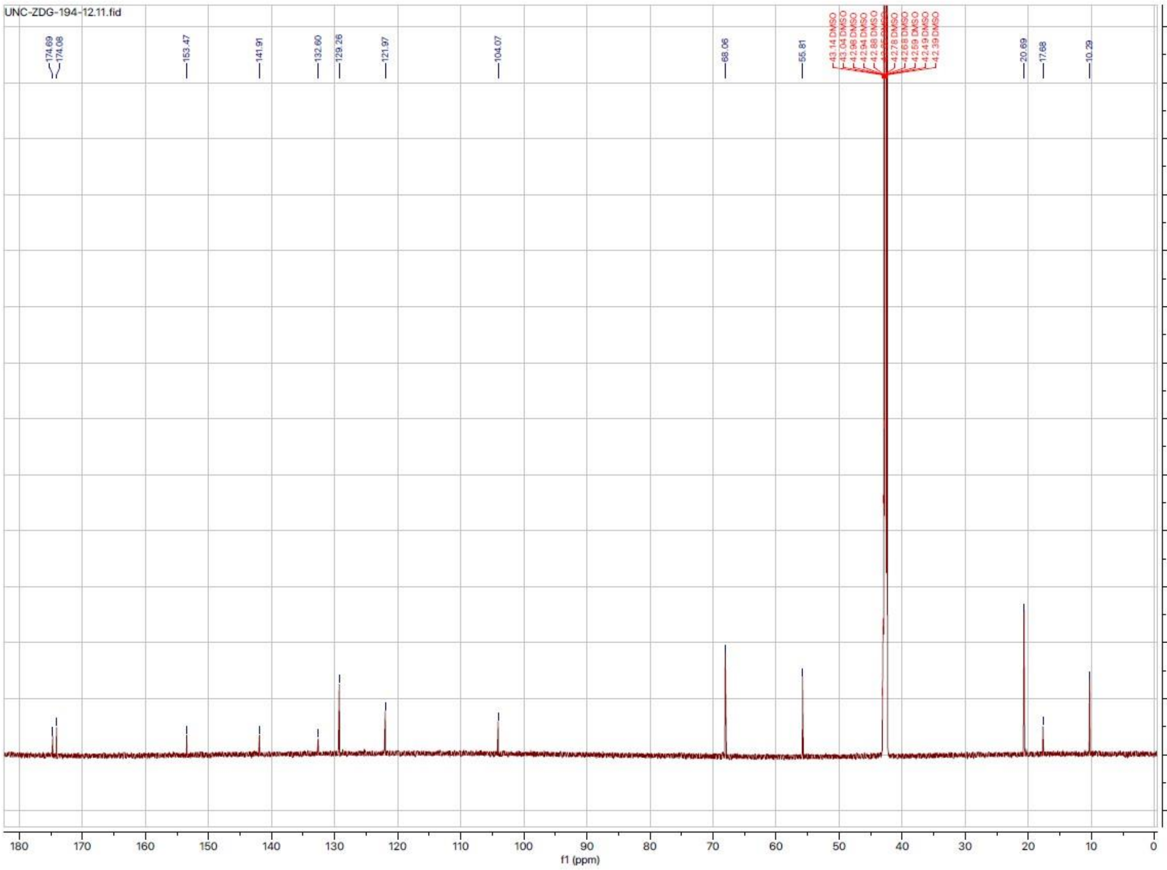

### 1H and 13C NMR of Compound 5

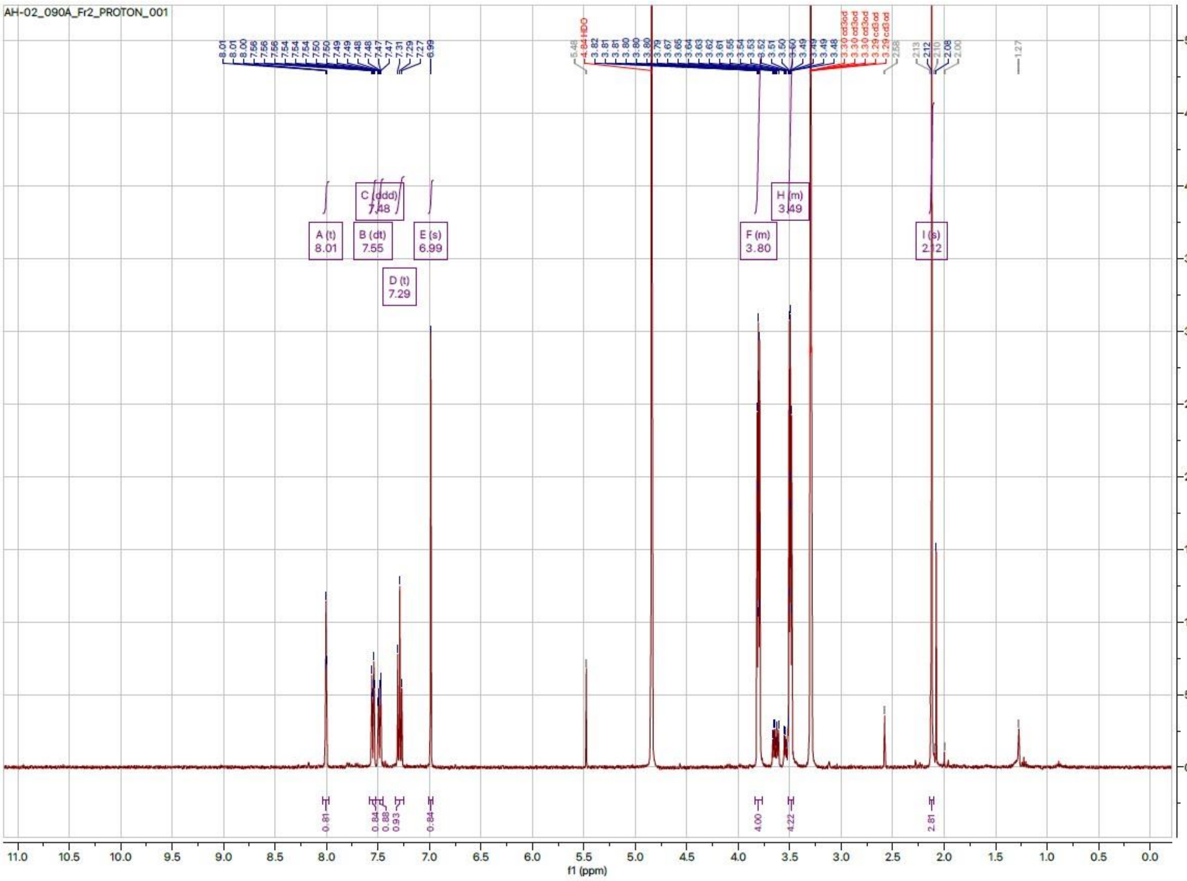


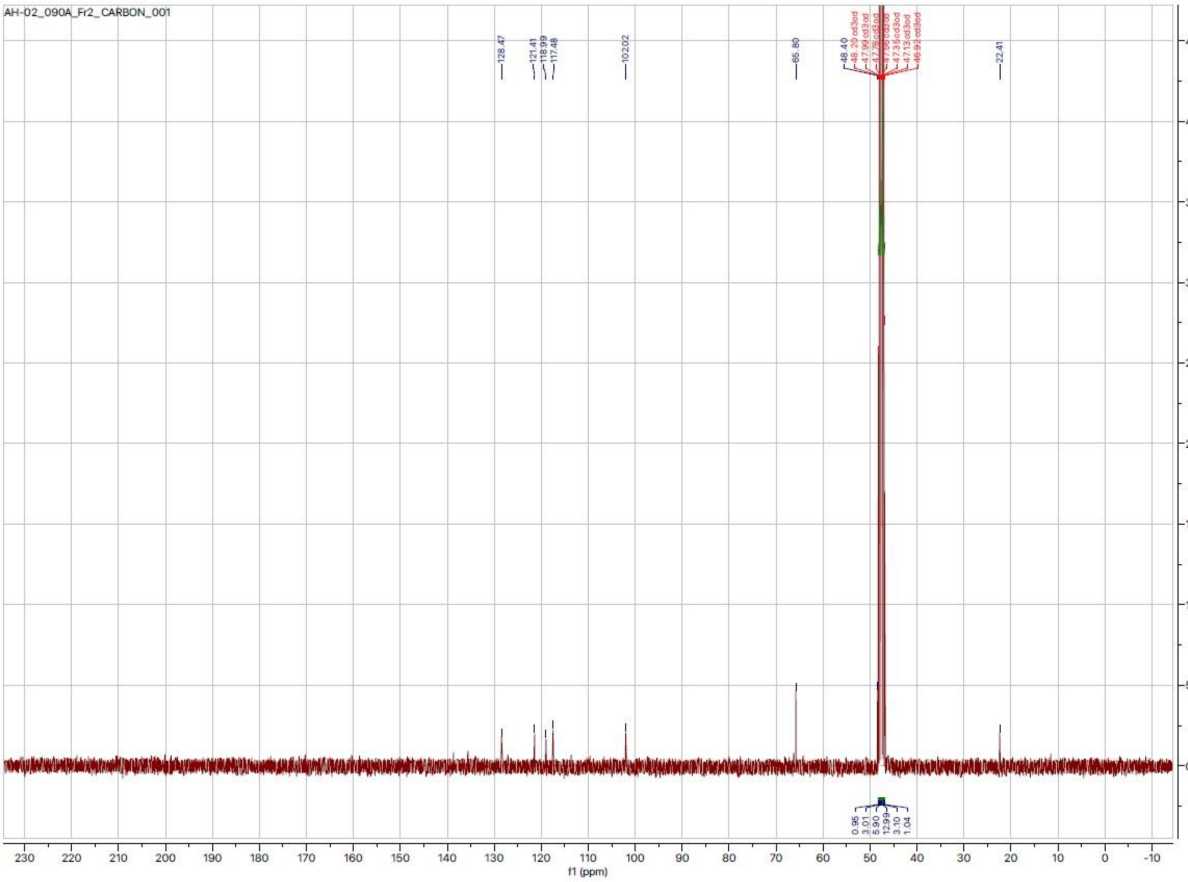

### 1H and 13C NMR of Compound 6

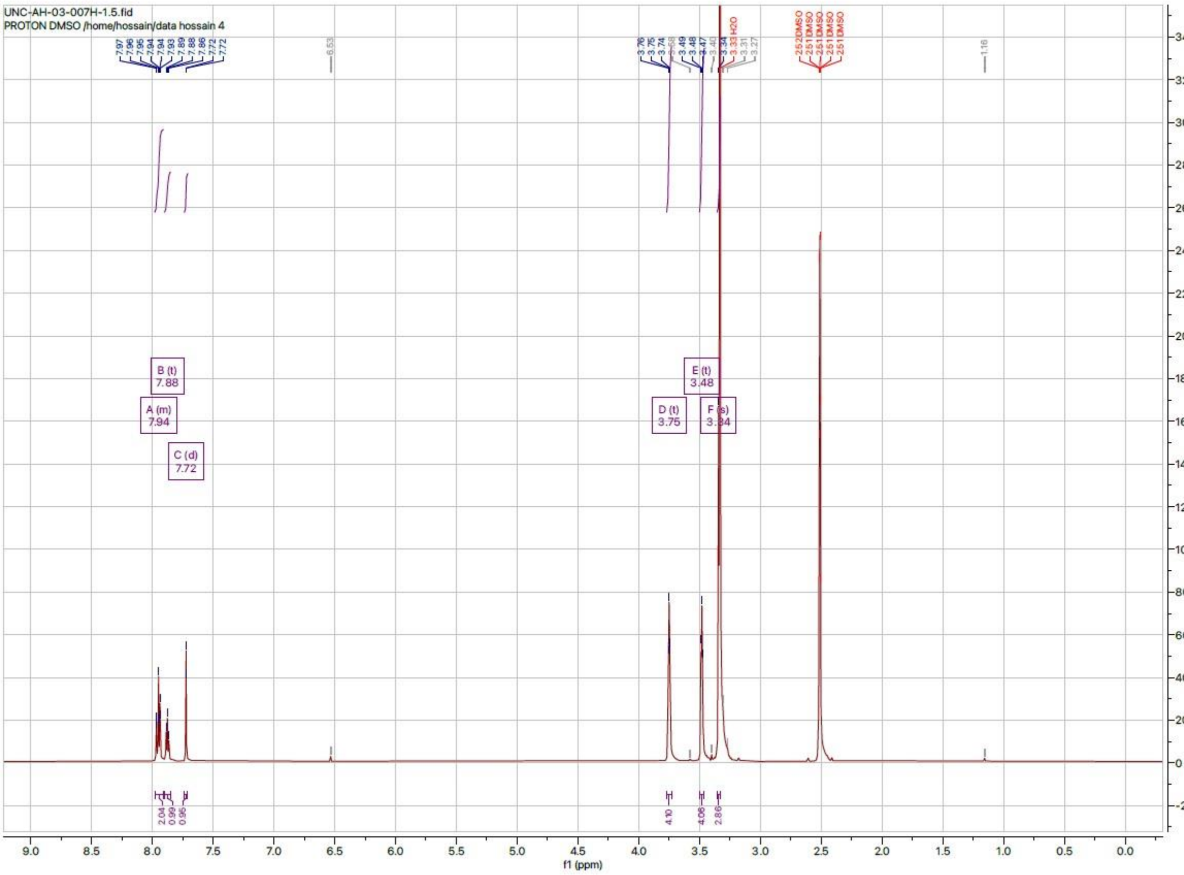


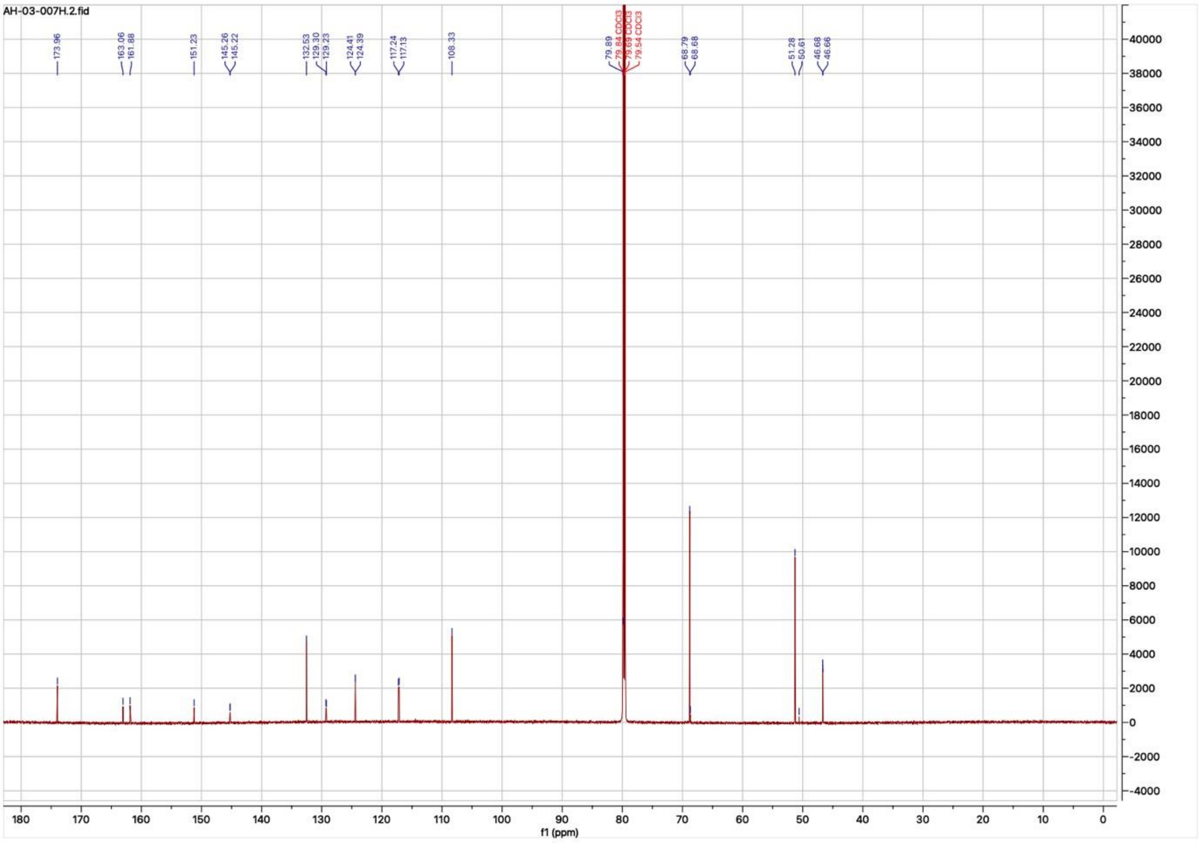

1H and 13C NR of Compound 7

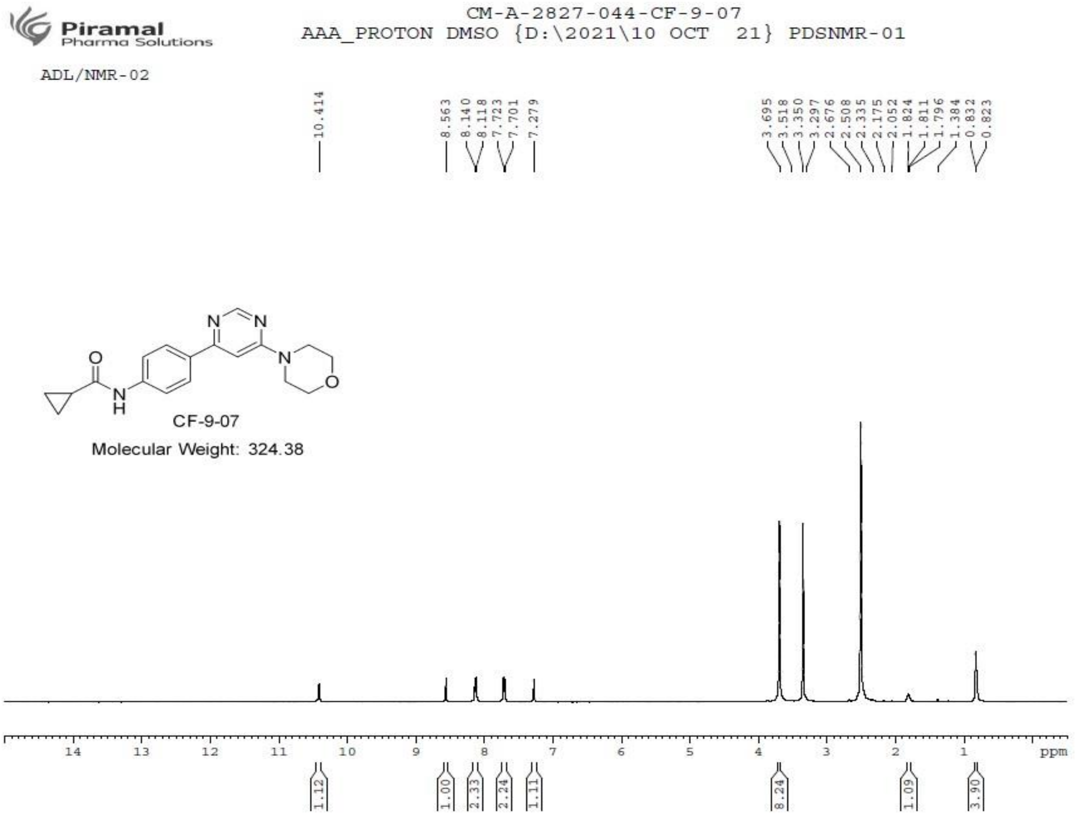


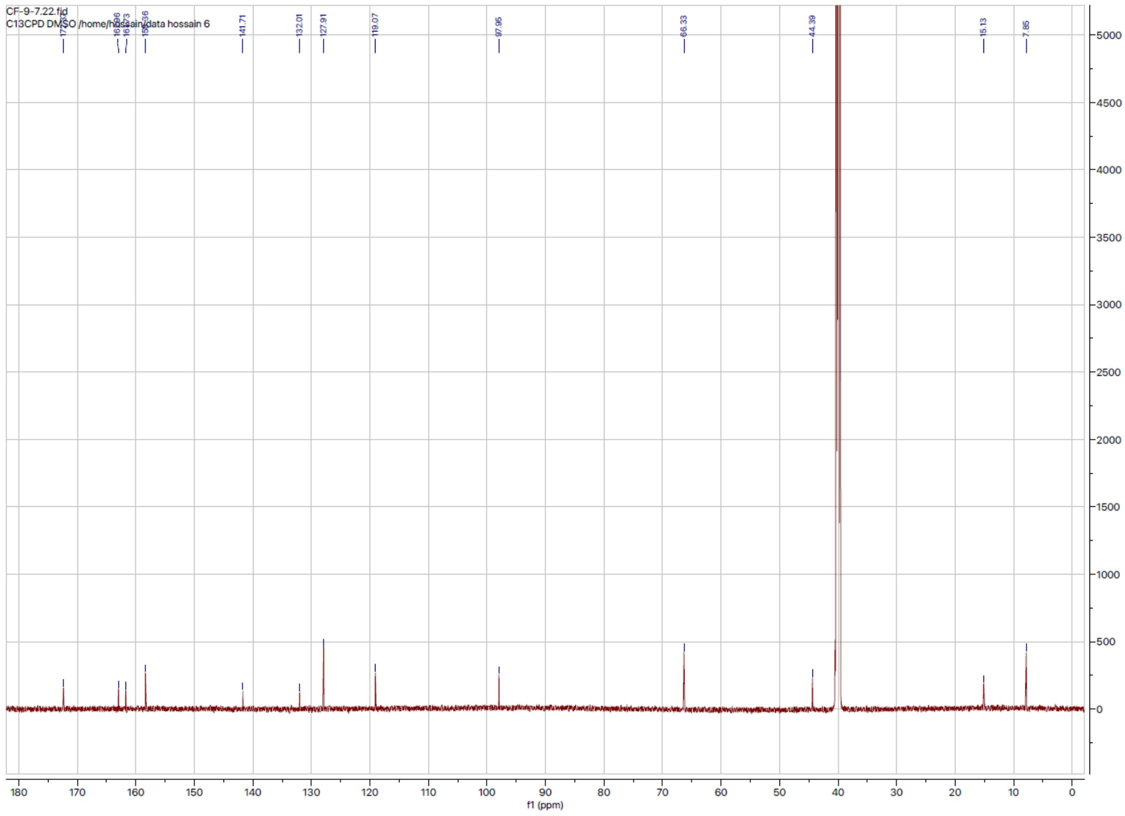

1H and 13C NMR of Compound 8

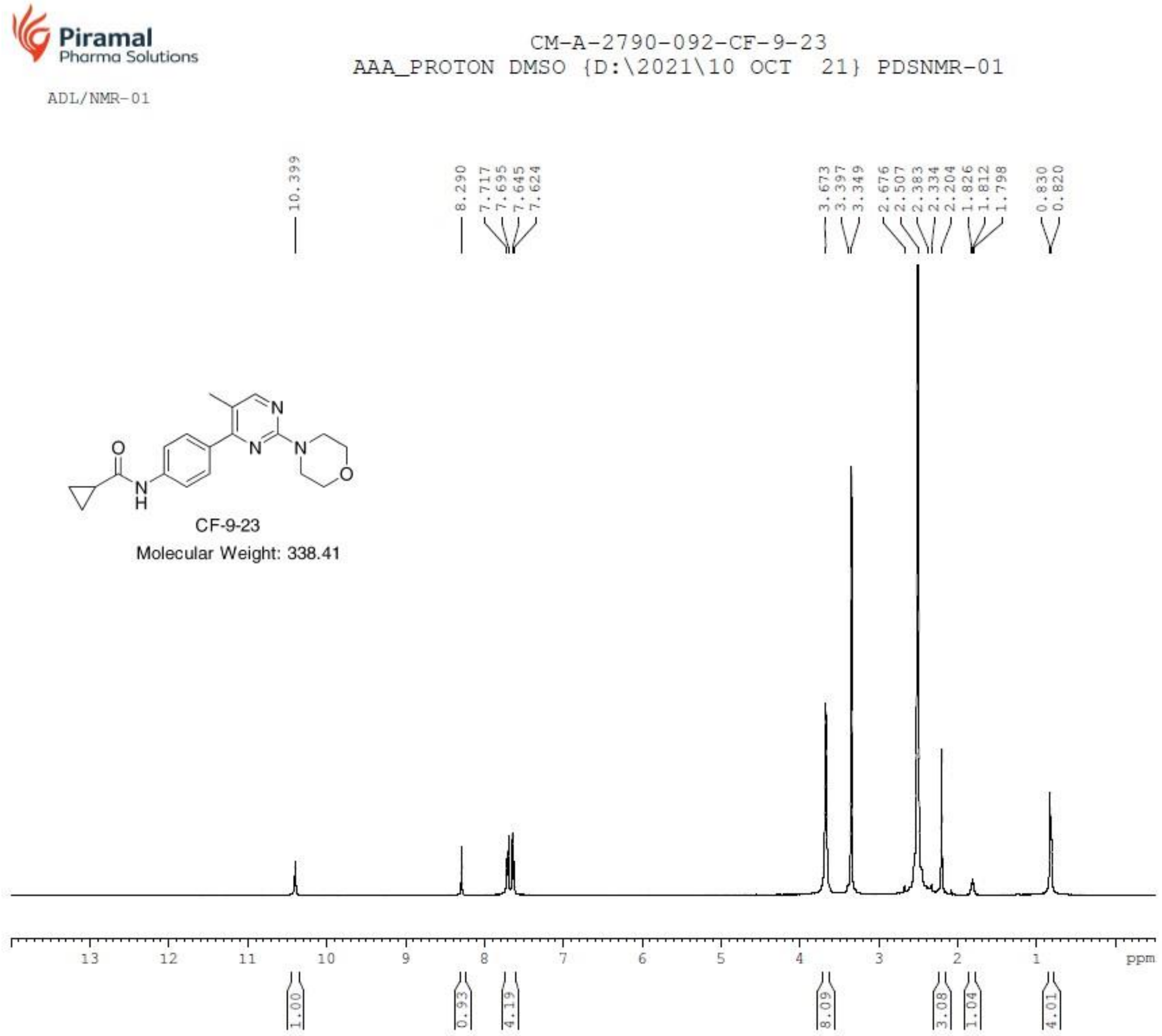

### 1H and 13C NMR of Compound 9

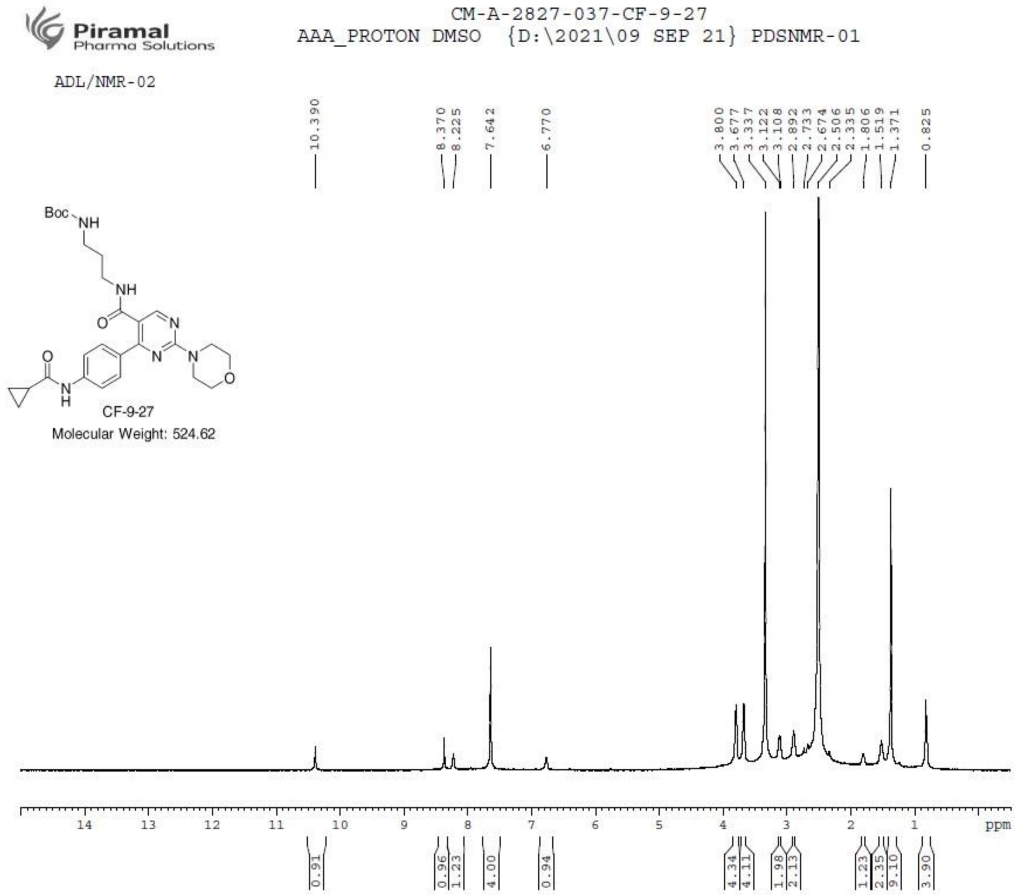


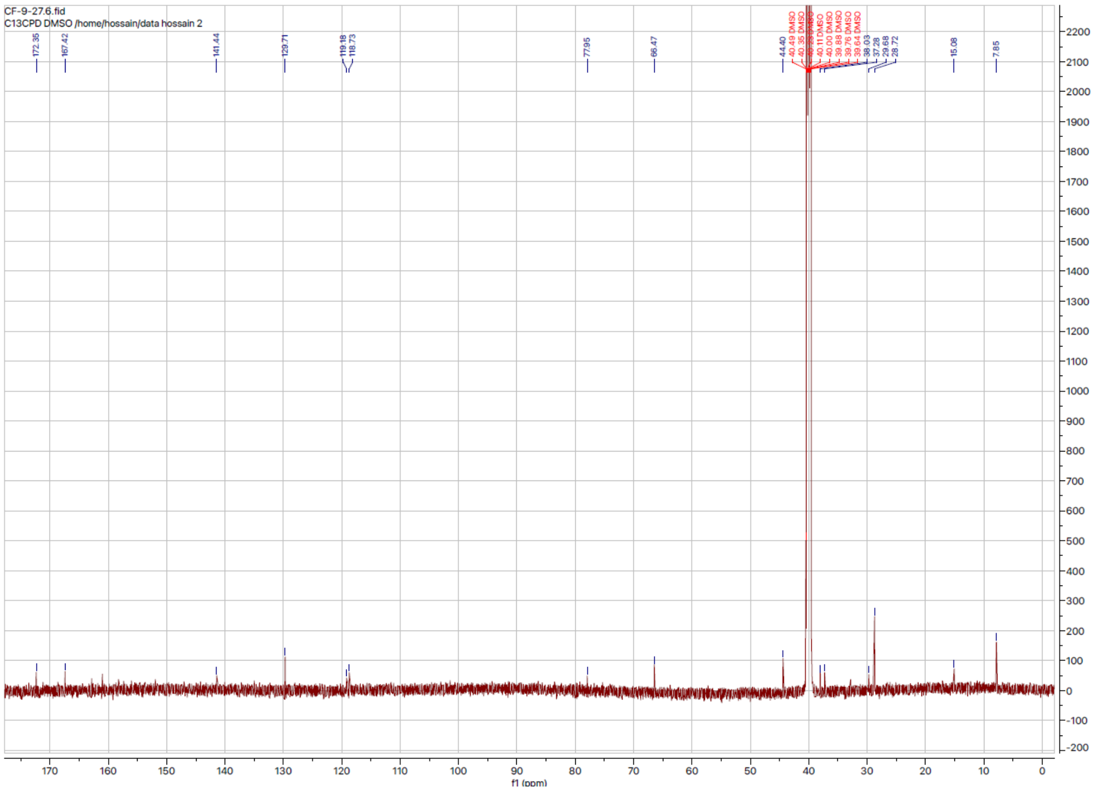

1H and 13C NMR of Compound 10

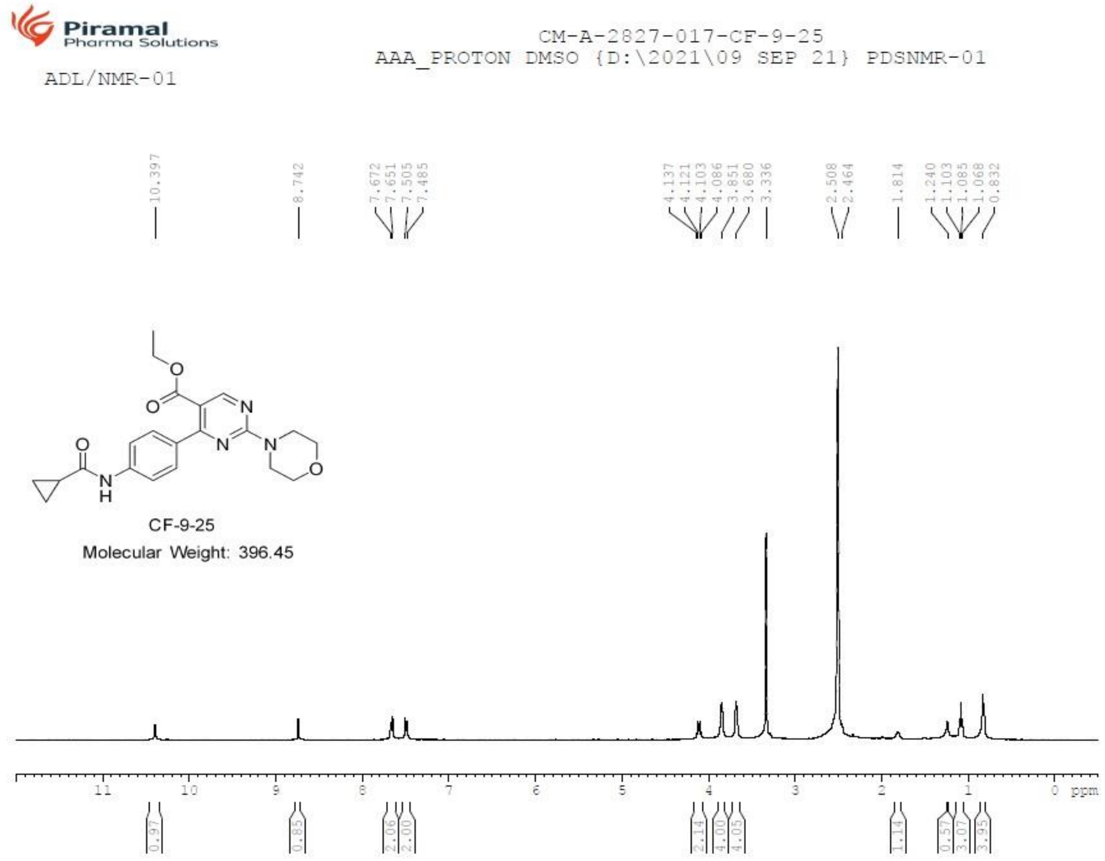


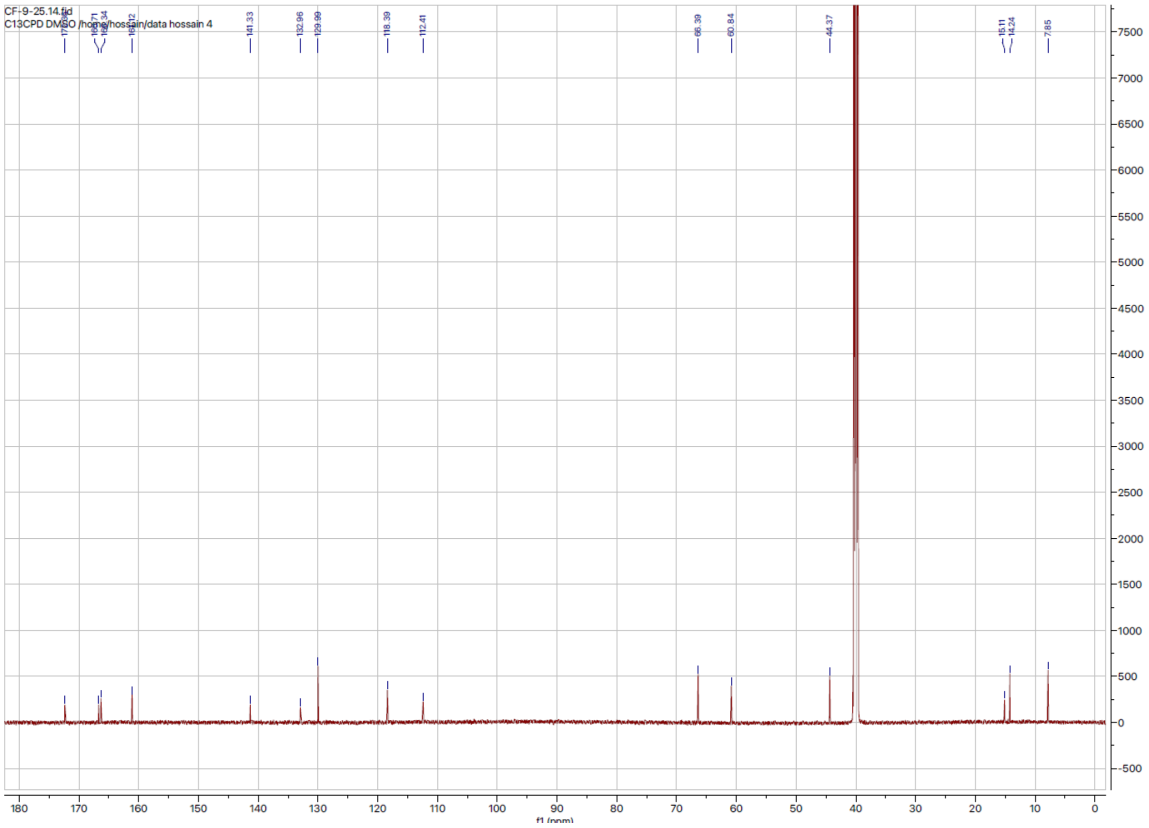

### 1H and 13C NMR of Compound 11

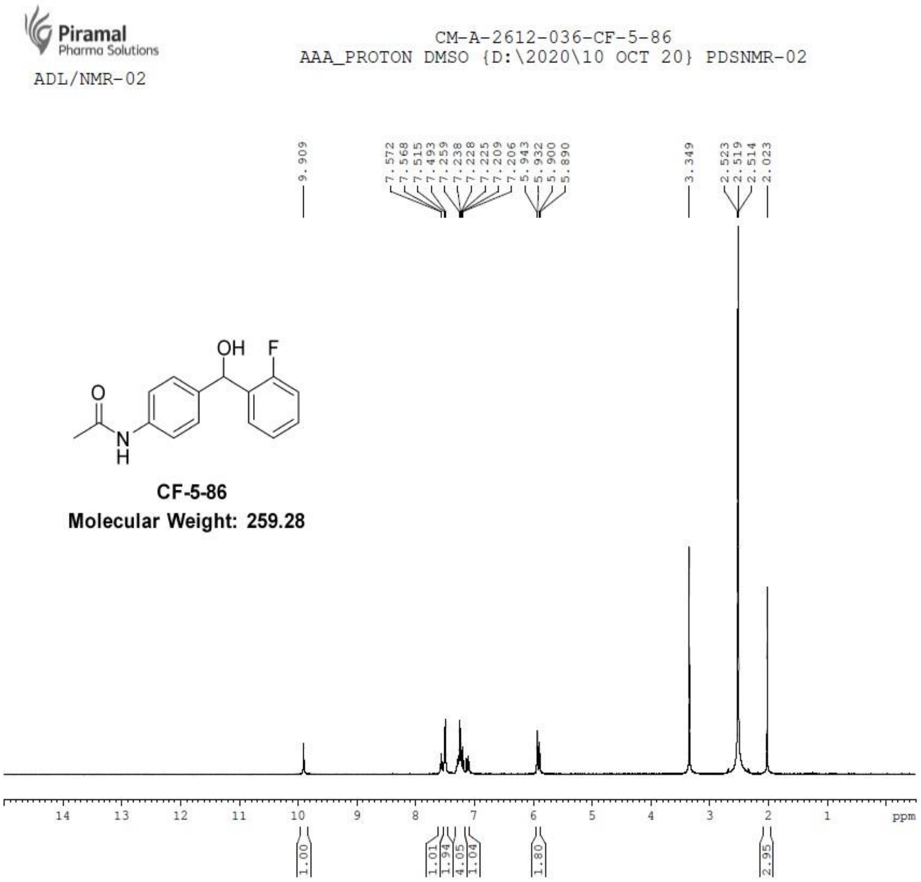


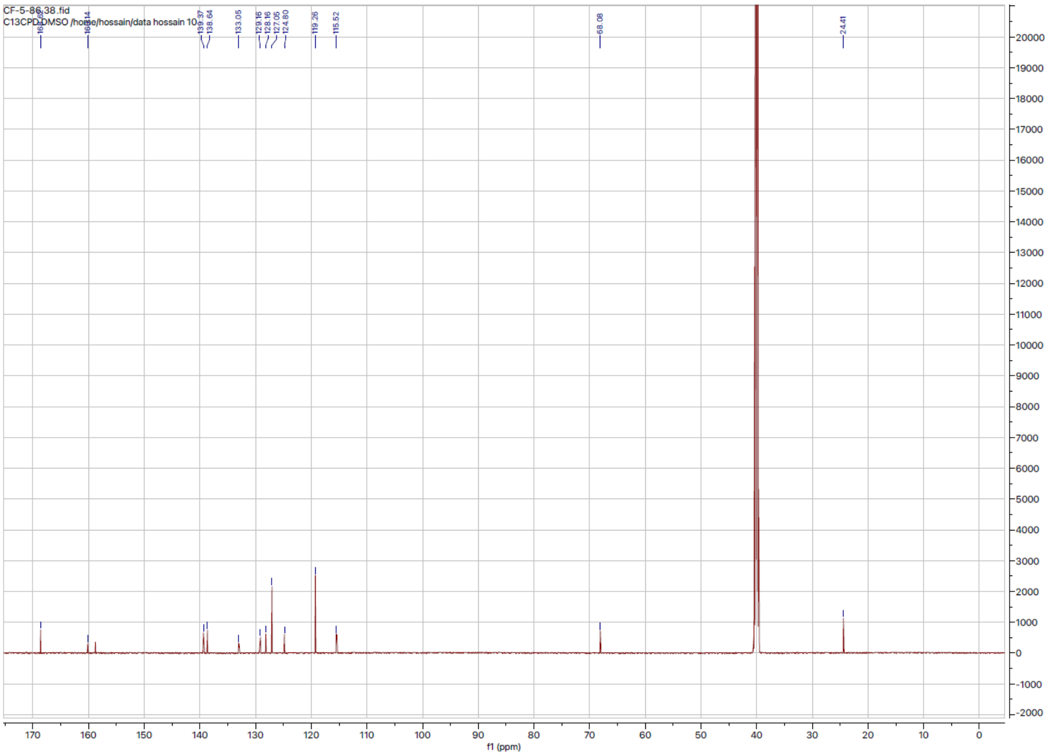

### 1H and 13C NMR of Compound 12

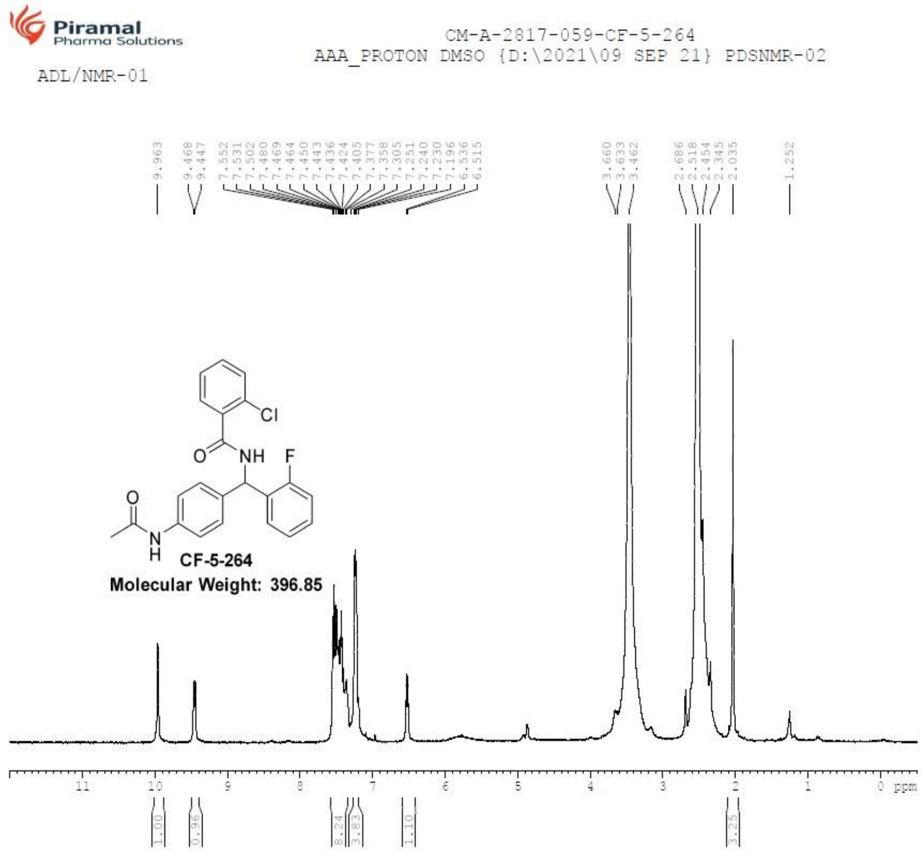


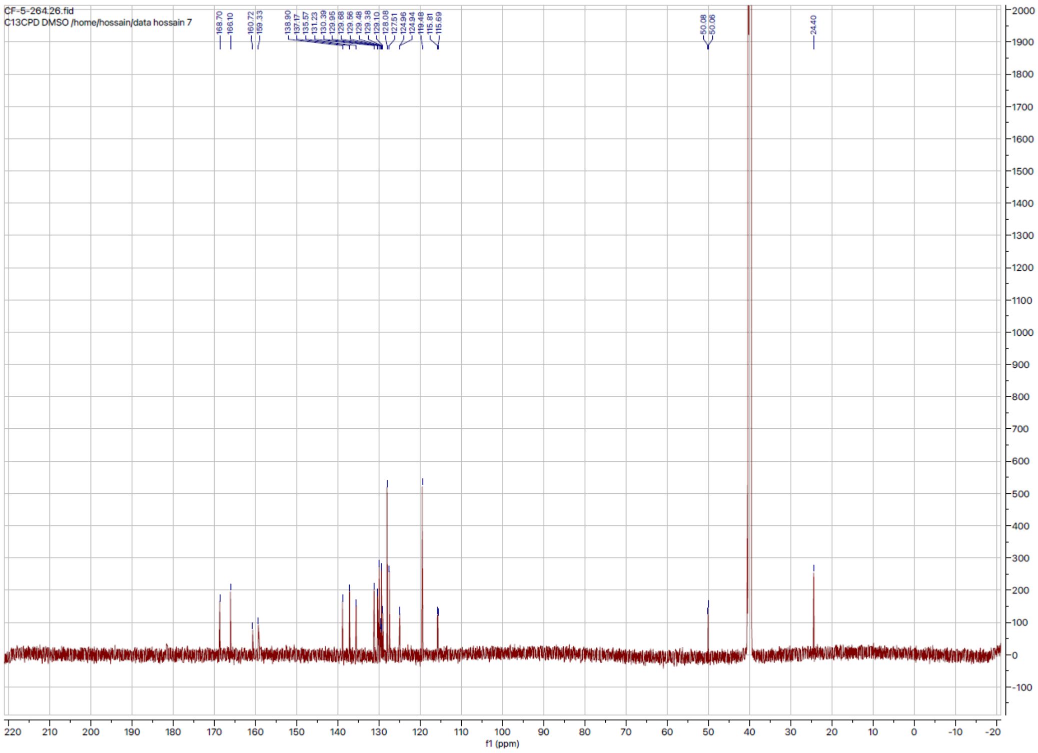

### 1H and 13C NMR of Compound 13

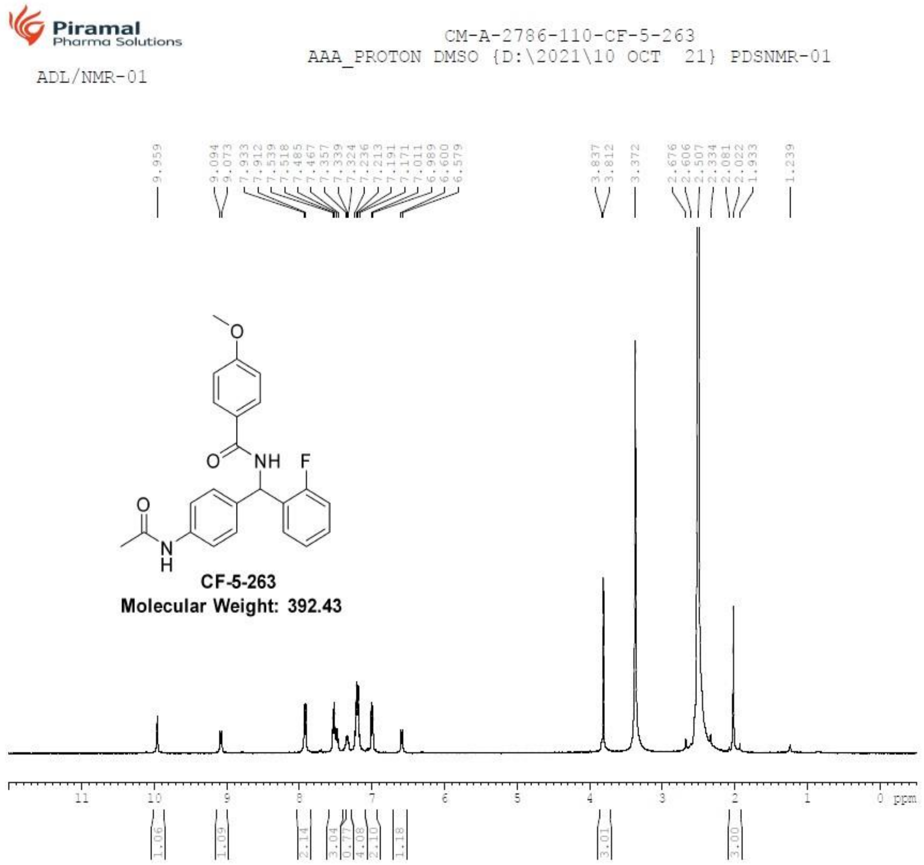


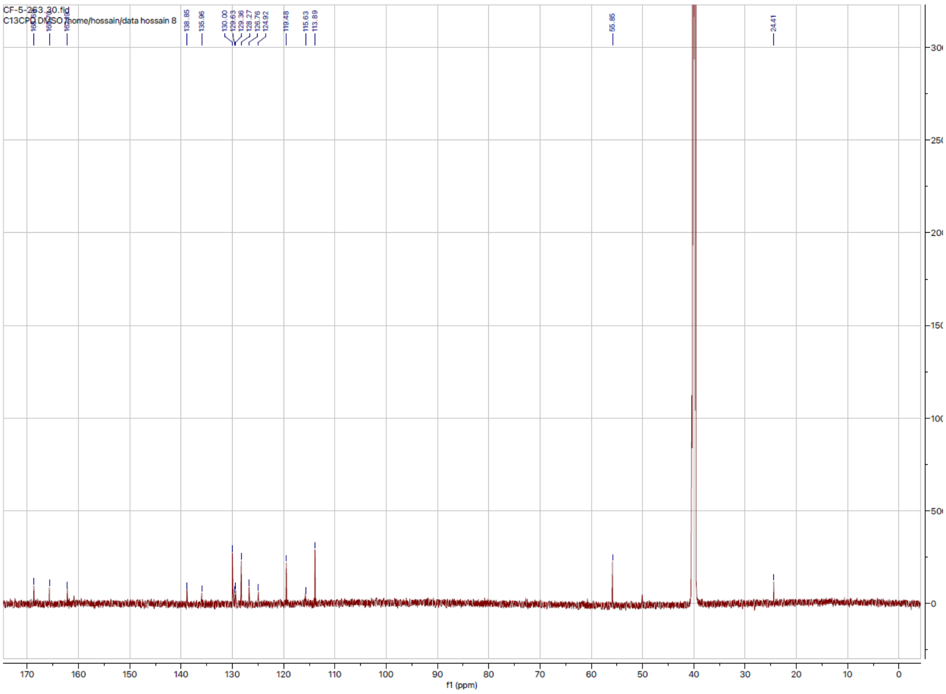
